# N-terminal acetylation shields proteins from degradation and promotes age-dependent motility and longevity

**DOI:** 10.1101/2022.09.01.505523

**Authors:** Sylvia Varland, Rui Duarte Silva, Ine Kjosås, Alexandra Faustino, Annelies Bogaert, Maximilian Billmann, Hadi Boukhatmi, Barbara Kellen, Michael Costanzo, Adrian Drazic, Camilla Osberg, Katherine Chan, Xiang Zhang, Amy Hin Yan Tong, Simonetta Andreazza, Juliette J. Lee, Lyudmila Nedyalkova, Matej Ušaj, Alexander J. Whitworth, Brenda J. Andrews, Jason Moffat, Chad L. Myers, Kris Gevaert, Charles Boone, Rui Gonçalo Martinho, Thomas Arnesen

## Abstract

Most eukaryotic proteins are N-terminally acetylated, but the functional impact on a global scale has remained obscure. Using genome-wide CRISPR knockout screens in human cells, we reveal a strong genetic dependency between a major N-terminal acetyltransferase and specific ubiquitin ligases. Biochemical analyses uncover that both the ubiquitin ligase complex UBR4-KCMF1 and the acetyltransferase NatC recognize proteins bearing an unacetylated N-terminal methionine followed by a hydrophobic residue. NatC KO-induced protein degradation and phenotypes are reversed by UBR knockdown, demonstrating the central cellular role of this interplay. We reveal that loss of *Drosophila* NatC is associated with male sterility, reduced longevity, and age-dependent loss of motility due to developmental muscle defects. Remarkably, muscle-specific overexpression of UbcE2M, one of the proteins targeted for NatC KO mediated degradation, suppresses defects of NatC deletion. In conclusion, NatC-mediated N-terminal acetylation acts as a protective mechanism against protein degradation, which is relevant for increased longevity and motility.

**In Brief:** Varland, Silva *et al*. define that a major cellular role of N-terminal acetylation is shielding proteins from proteasomal degradation by specific ubiquitin ligases. The human N-terminal acetyltransferase NatC protects the neddylation regulator UBE2M from degradation, while overexpression of *Drosophila* UBE2M/UbcE2M rescues the longevity and motility defects of NatC deletion.

**Highlights:**

- N-terminal acetylation by NatC protects proteins from degradation, including UBE2M
- UBR4-KCMF1 targets unacetylated N-terminal Met followed by a hydrophobic residue
- *Drosophila* NatC is required for adult longevity and motility in elderly
- Overexpression of UBE2M/UbcE2M suppresses *Drosophila* NatC deletion phenotypes

## INTRODUCTION

The N-terminal protein landscape is a hotspot for modifications influencing protein interactions and homeostasis (Varland et al., 2015). N-terminal acetylation (Nt-acetylation) is undoubtedly one of the most common protein modifications in eukaryotes, with the α-amino group of ~80-90% of human proteins being susceptible to acetylation by N-terminal acetyltransferases (NATs) (Arnesen et al., 2009; Ree et al., 2018). Acetylation transforms a charged protein N-terminus into a hydrophobic segment, thereby affecting key protein properties such as folding, polymerization, stability, interactions, and localization. Given the prevalence of Nt-acetylation, NATs regulate a wide range of cellular processes, ranging from metabolism, cell proliferation and migration to differentiation and stress response (Aksnes et al., 2019). Dysregulation of Nt-acetylation can cause developmental disorders, by affecting brain and heart development, and contribute to cancer development (Aksnes et al., 2019; Cheng et al., 2018; Morrison et al., 2021; Muffels et al., 2021; Rope et al., 2011; Ward et al., 2021).

The human NAT family comprises five ribosome-associated members (NatA-NatE) that acetylate nascent polypeptides during translation (Aksnes et al., 2016), as well as the Golgi-associated NAA60/NatF (Aksnes et al., 2015) and the actin-specific NAA80/NatH (Drazic et al., 2018) which both act post-translationally. The NATs mainly recognize the first two amino acids at the N-terminus, but local sequence context may affect substrate binding and the degree of acetylation. NatA acetylates small N-terminal residues (Arnesen et al., 2009), which are exposed after the initiator methionine has been removed by methionine aminopeptidases. Proteins harboring a N-terminal methionine can be modified by NatB/C/E/F. In this substrate class, NatB acetylates the α-amino group of methionine followed by a negatively charged residue (Van Damme et al., 2012), while NatC, NatE and NatF acetylates α-amino group of methionine followed by a hydrophobic or amphipathic residue (Aksnes et al., 2016; Van Damme et al., 2016).Structural studies of NatC have shown that the first four amino acids contribute to substrate recognition, and presumably NatC, NatE and NatF act on different substrates classes *in vivo* (Deng et al., 2021; Grunwald et al., 2020; Tercero et al., 1993).

The evolutionarily conserved NatC complex consists of the catalytic subunit NAA30, the ribosomal anchor NAA35, and the auxiliary subunit NAA38, all of which are required for normal enzymatic activity (Grunwald et al., 2020; Polevoda and Sherman, 2001; Starheim et al., 2009). NatC-mediated acetylation increases the affinity of the two NEDD8-conjugating enzymes UBE2M/UBC12 and UBE2F to E3 ligases, promoting cullin neddylation (Monda et al., 2013; Scott et al., 2011). Structural analysis showed that Ac-UBE2M is buried within a hydrophobic pocket of DCN1, which enhances cullin neddylation (Scott et al., 2011), and this acetylation-dependent interaction can be antagonized by inhibitors (Scott et al., 2017). Acetylation by NatC can be crucial for subcellular targeting, partly by mediating protein interactions. The GTPases ARL8B and ARFRP1 (Arl3 in yeast) rely on acetylation of their N-terminal methionine for correct targeting to the lysosomes and Golgi, respectively (Behnia et al., 2004; Setty et al., 2004; Starheim et al., 2009). The latter example is driven by an interaction between the acetylated N-terminus and the membrane protein SYS1. Several studies have also linked NatC activity to development, stress response and longevity (Cao et al., 2019; Pesaresi et al., 2003; Warnhoff et al., 2014; Wenzlau et al., 2006), whereas the catalytic subunit of NatC, NAA30, was shown to regulate cancer cell viability and tumorigenesis of glioblastoma initiating cells (Mughal et al., 2015; Starheim et al., 2009; Varland et al., 2018b).

The N-terminus of a protein can, depending on its nature, act as a degradation signal (N-degron) that is recognized by N-degron pathways and targeted for degradation by the 26S proteasome or lysosomes (via autophagy). In fact, most cellular proteins can be targeted for degradation by different N-degron pathways (Chen et al., 2017; Dong et al., 2020; Timms et al., 2019; Varshavsky, 2019). The Ac/N-degron pathway targets proteins harboring an acetylated N-terminal residue (mainly Met, Ala, Val, Ser, Thr and Cys) and is mediated by the E3 ubiquitin ligases Doa10 (MARCH6 in mammals) or Not4 (Hwang et al., 2010; Oh et al., 2017). The Arg/N-degron pathway targets specific unacetylated N-terminal residues, termed type 1 (basic; Arg, Lys and His) and type 2 (bulky hydrophobic; Leu, Phe, Tyr, Trp, or Ile), that are exposed after proteolytic processing (Bachmair et al., 1986; Varshavsky, 2019). The Arg/N-degron pathway also recognizes unacetylated N-terminal methionine when followed by a bulky hydrophobic residue in yeast (Kim et al., 2014). While the E3 ubiquitin ligase Ubr1 acts in the yeast Arg/N-degron pathway, in mammals there are at least four N-recognins, UBR1, UBR2, UBR4, and UBR5, whose individual contributions for protein degradation have not been extensively explored (Tasaki et al., 2005; Tasaki et al., 2009). Many proteins are conditional N-degron substrates that are only available under certain circumstances. For example, natural Ac/N-degrons can be shielded within protein complexes (Shemorry et al., 2013). Consequently, the N-degron pathways act as important control mechanisms to ensure protein quality and subunit stoichiometry.

Acetylation of protein N-termini underlies several key biological processes. Although NatC acetylates hundreds of different substrates, we know very little about the functional and physiological importance of these acetylation events. While the genes encoding the human NatA and NatB subunits are mostly essential (Blomen et al., 2015; Hart et al., 2015; Wang et al., 2015), human NatC knockout cell models enabled us to investigate the overall role of Nt-acetylation. To this end we performed unbiased genome-wide CRISPR knockout screens in human HAP1 cells. NatC perturbation appears to regulate vesicle trafficking and organelle morphology. Moreover, we uncover a key role of Nt-acetylation in protecting hydrophobic N-termini against targeted degradation by the Arg/N-degron pathway. The organismal relevance of NatC was investigated in *Drosophila melanogaster*. Altogether, our findings reveal how NatC-mediated acetylation promotes healthy proteostasis by regulating protein quality control, and impacts motility among elderly and longevity in flies.

## RESULTS

### NatC subunits show strong genetic interactions with components of the Arg/N-degron pathway

Genetic interactions (GIs) occur when the combination of mutations in different genes leads to an unexpected phenotype, considering the effects of the individual gene mutations, and can reveal functional relationships between genes and pathways (Costanzo et al., 2019; Costanzo et al., 2016). Mapping GI profiles has thus become a powerful approach for deciphering gene function. To systematically identify GIs for NatC, we performed genome-wide CRISPR knockout (KO) screens in human HAP1 WT and *NAA30*-KO, *NAA35*-KO or *NAA38*-KO cell lines (collectively referred to as NatC KO) (**Figure S1A**) using the TKOv3 guide RNA (gRNA) library targeting ~18,000 protein-coding genes (**Figure 1A**). The relative abundance of a specific gRNA between the start and end timepoint of a screen provides an estimate of single mutant fitness in WT cells and double-mutant fitness in KO cells. We have previously developed a quantitative GI (qGI) score that compares the double-mutant fitness effects in a query KO cell line with the single-mutant fitness effects in a panel of HAP1 WT control screens and corrects for various experimental artifacts (Aregger et al., 2020). In this context, a negative GI occurs when the simultaneous knockout of two genes leads to a more severe cell fitness decrease than expected by considering the individual effects of both genes. In our screen, this is identified by reduced gRNA abundance in the KO cell line relative to the WT control cell line. Conversely, a positive GI occurs when the combined disruption of two genes promotes cell fitness, and results in increased gRNA abundance in the KO cells compared to WT cells.

**Figure 1.**
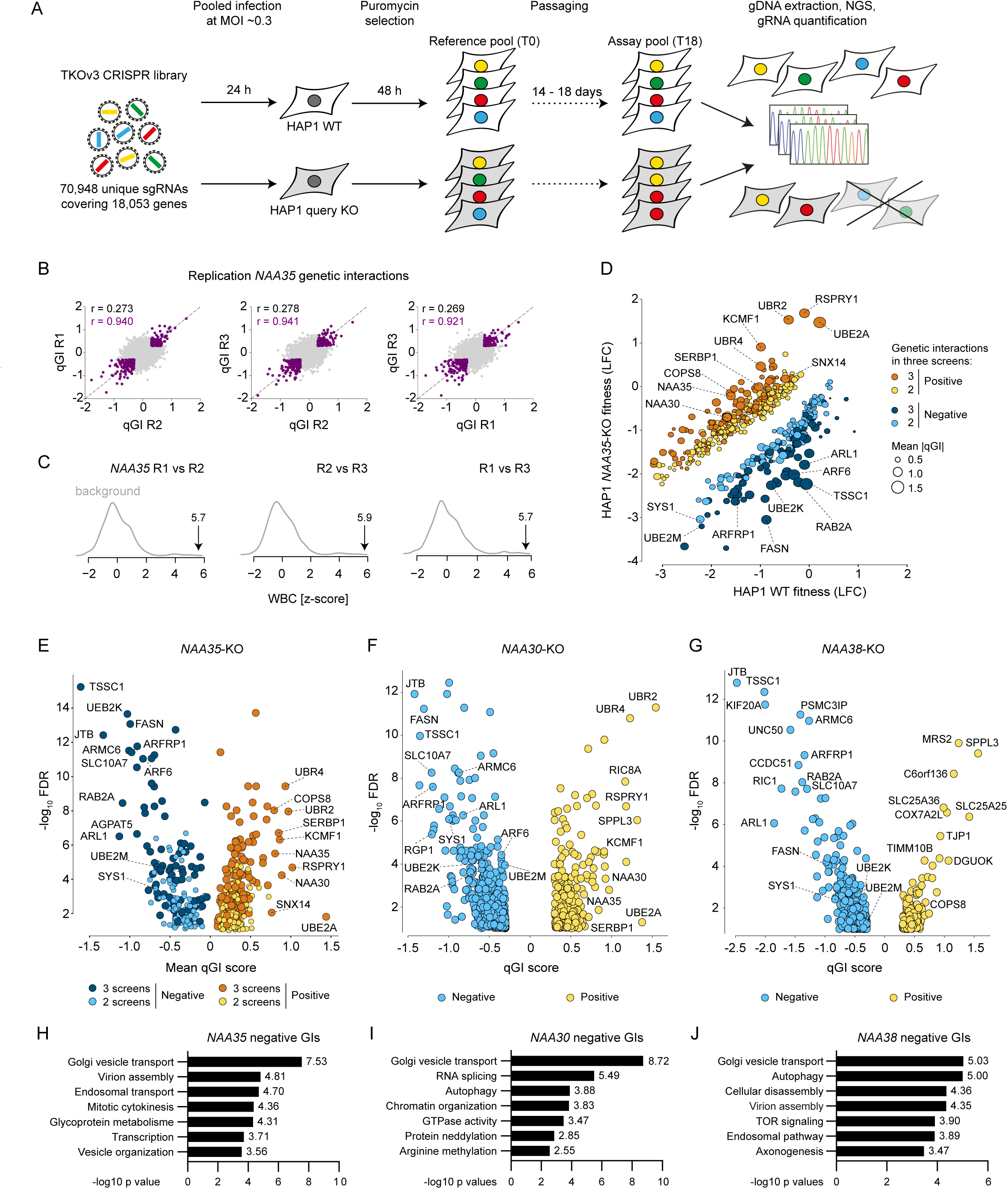
Genome-wide mapping of genetic interactions with human NatC. **(A)** Schematic of genome-wide CRISPR screens to identify genetic interactions (GI) with NatC. HAP1 WT, *NAA30*-KO, *NAA35*-KO, and *NAA38*-KO cells were transduced with a pooled lentiviral genome-wide CRISPR knockout library (TKOv3). The cells were selected for viral integration and passaged over time to allow gene-editing to take place. gRNA regions were amplified by PCR from genomic DNA extracted from cells collected at the start (T0) and at the endpoint of the screen (T14-18). gRNA abundance was determined by next-generation sequencing (NGS). **(B-C)** Reproducibility of *NAA35* qGI scores. **(B)** qGI scores were determined by comparing the LFC for every gene represented in the TKOv3 library in a *NAA35*-KO cell line with those observed in a panel of WT control screens. Pearson correlation coefficient (*r*) was calculated using all qGI scores (*r* in black, calculated from all data points) or using a stringent cut-off for the GIs (|qGI| > 0.3, FDR < 0.10) in both screens (*r* and datapoints marked in purple). **(C)** The Pearson correlation coefficients of the qGI scores measured in two replicated screens was adjusted to the similarity of a *NAA35*-KO screen to a panel of HAP1-KO screens. The resulting Within vs Between replicate Correlation (WBC) score provides a confidence of reproducibility that can be interpreted as a z-score. **(D)** Negative and positive GIs of *NAA35*. Scatterplot showing the fitness effect (LFC) of 486 genes in *NAA35-*KO versus WT cell line, which showed a significant GI in at least two *NAA35* screens (|qGI| > 0.3, FDR < 0.10). Negative (blue) and positive (yellow) *NAA35* GIs are shown. Darker color indicates interactions were called in all three replicate screens. Node size corresponds to strength of the mean absolute GI score derived from three independent screens. Selected GIs are indicated. **(E**-**G)** Volcano plots displaying qGI scores and associated significance (log10 values) for the genes targeted by the TKOv3 library in **(E)** *NAA35*-KO, **(F)** *NAA30*-KO and **(G)** *NAA38*-KO screens. Negative and positive GIs are marked in blue and yellow, respectively. Selected GIs are indicated. **(H-J)** Negative GIs of NatC indicate a role in Golgi vesicle transport. Pathway enrichment analysis of genes exhibiting a negative GI with **(H)** *NAA35*, **(I)** *NAA30* or **(J)** *NAA38* (identified in at least two *NAA35* screens; for all screens |qGI| > 0.3, FDR < 0.1). The p-value for each gene ontology term is indicated.

We performed three independent screens using a *NAA35*-KO query cell line to establish a high-confidence GI dataset for the major NatC auxiliary subunit. The reproducibility of the double-mutant fitness effects (log2-fold change, LFC) between the three *NAA35* screens were strongly correlated (pairwise *r* = 0.81) (**Figure S1B**). Furthermore, we observed only a modest correlation between the genome-wide qGI scores of the *NAA35* replicate screens (*r* = 0.27-0.28) (**Figure 1B**), which was expected given the relative sparsity of GIs (Aregger et al., 2020). When restricting our correlation analysis to significant GIs in pairs of replicate *NAA35* screens (|qGI| > 0.30; FDR < 0.10) the pairwise correlation increased considerably (*r* = 0.92-0.94), suggesting that significant GIs were reproducible. Indeed, the qGI scores measured in the *NAA35* screens were highly reproducible with a Within vs Between Correlation (WBC) score (adjusted z-score) between 5.7 and 5.9 (**Figure 1C**) (Billmann et al., 2022).

We next generated a combined set of *NAA35* GIs by mean-summarizing the gGI scores of the three replicate screens (**Figures 1D-E**). Negative GIs with *NAA35* included *TSSC1/EIPR1*, *JTB*, *ARL1*, *RAB2A*, *FASN*, *SLC10A7*, *ARF6* and *ARFRP1* possibly reflecting NatC’s impact on the secretory pathway and organelles (Starheim et al., 2009; Starheim et al., 2017; Van Damme et al., 2016). Indeed, negative *NAA35* GIs were enriched for genes involved in Golgi vesicle transport, virion assembly, and endosomal transport (FDR < 0.20) (**Figure 1H**, **Table S2**). The role of NatC in vesicle trafficking is most likely evolutionarily conserved as many of the same NatC GIs were observed in the global yeast genetic network (*ARL1*, *SYS1*, *ARFRP1*, *RIC1/KIAA1432*, *RGP1*, *COG5*, and *COG7*) (Costanzo et al., 2010; Costanzo et al., 2016). Arl3/ARFRP1 is a small GTPase which recruits Arl1 and its effectors to the *trans*-Golgi. The targeting of Arl3 to the Golgi requires NatC-mediated acetylation and the membrane protein Sys1 (Behnia et al., 2004; Setty et al., 2004). COG5 and COG7 are components of the conserved oligomeric Golgi (COG) complex, which acts in *intra*-Golgi protein transport (Gillingham and Munro, 2016). We also observed that several members of the Rab family of small GTPases (*RAB1A*, *RAB1B*, *RAB2A*, *RAB14*) regulating membrane trafficking (Homma et al., 2021) have negative GIs with *NAA35*, emphasizing the role of NatC in intracellular transport. The strongest positive qGI scores were also highly reproducible and included components of the Arg/N-degron pathway *UBE2A, UBR2*, *UBR4*, *KCMF1*, a component of the COP9 signalosome promoting cullin deneddylation *COPS8*, as well as the NatC genes *NAA30* and *NAA35* confirming the responsiveness of the screen. Moreover, *NAA35* positive GIs were enriched for genes with annotated roles in ribosome biogenesis, RNA processing and translation (**Table S2**), suggesting that loss of NatC might be buffered by translational perturbations.

To better understand the genetic dependencies of the NatC complex, we next performed genome-wide CRISPR KO screens in *NAA30*-KO and *NAA38*-KO cell lines (**Figures 1F-G**, **Table S1**). Each query screen was performed in technical triplicates, and we used the same confidence threshold as for the *NAA35* screens (|qGI| > 0.30, FDR < 0.10). Notably, both *NAA30* and *NAA38* negative GI profiles revealed genes involved in Golgi vesicle transport (**Figures 1I-J**). We next compared the NatC KO screens. Overall, 63 and 65 genes in the *NAA30* screen also significantly interacted with *NAA35* in 2 or 3 screens, respectively (128 genes in total). The *NAA38* GI profile contains 137 and 75 genes that also significantly interacted with *NAA30* and *NAA35* in at least 2 screens, respectively. We observed that the positive GIs were less shared between *NAA30*, *NAA35,* and *NAA38*, most likely caused by the *NAA38*-KO cells having increased proliferation rate compared to the other cell lines (**Figures S1C-D**). Taken together, the negative GIs of NatC suggest cellular roles related to the secretory pathway and organellar biology, while the positive GIs propose a functional link to the N-degron pathway and proteostasis.

### NAA30, NAA35, and NAA38 are essential for cellular Nt-acetylation and proteostasis of NatC-type substrates

To explore the dependencies of the NAA30, NAA35, and NAA38 subunits for NatC-mediated Nt-acetylation in human cells, we performed N-terminal proteomics using the same cell models. All three subunits were important for Nt-acetylation of different NatC-type substrates, including the IST1 protein harboring a Met-Leu-(ML) N-terminus (**Figures 2A-B**, **Table S3**). While IST1 was >90% Nt-acetylated in WT cells, it was <10% Nt-acetylated in all three NatC KO cell lines. Also, endogenous IST1 protein levels were significantly reduced in all NatC KO cell lines as compared to WT cells (**Figure 2C**). For several other proteins, such as the previously defined NatC substrates UBE2M (MI-) (Scott et al., 2011) and ATXN2L (ML-) (Van Damme et al., 2016) and the putative substrates DIS3 (ML-), ADNP (MF-), ADNP2 (MF-) and ITM2A (MV-), the Nt-acetylation levels could only be determined in WT cells and not in NatC KO cells, indicating that these proteins were less abundant (**Table S3**). To further address this issue, we performed label-free quantitative (LFQ) proteomics on HAP1 WT and NatC KO cells. We identified 440 proteins that were differentially expressed between WT and NatC KOs using multiple ANOVA testing (log2 transformed LFQ values, FDR = 0.01, S0 = 0) (**Table S4**). A pairwise comparison between the individual query KO cell lines with the WT control cell line identified 961 proteins that were differentially expressed (two-sample t test, FDR = 0.01, S0 = 0.1), of which 37% were enriched while 63% were depleted compared to WT. Notably, the NatC KO cell lines had different effects on the proteome and most deregulated proteins were identified in the *NAA35*-KO cells. However, most proteins that were identified in the *NAA30*-KO and *NAA38*-KO cells were also identified in the *NAA35*-KO cells, indicating a shared NatC effect. In total, 12 depleted and 10 enriched proteins were shared among the NatC KO cells, of which CAPNS1 (MF-), DARS (MY-), and FLNC (MM-) have NatC/E/F-type N-termini (**Table S4**).

**Figure 2.**
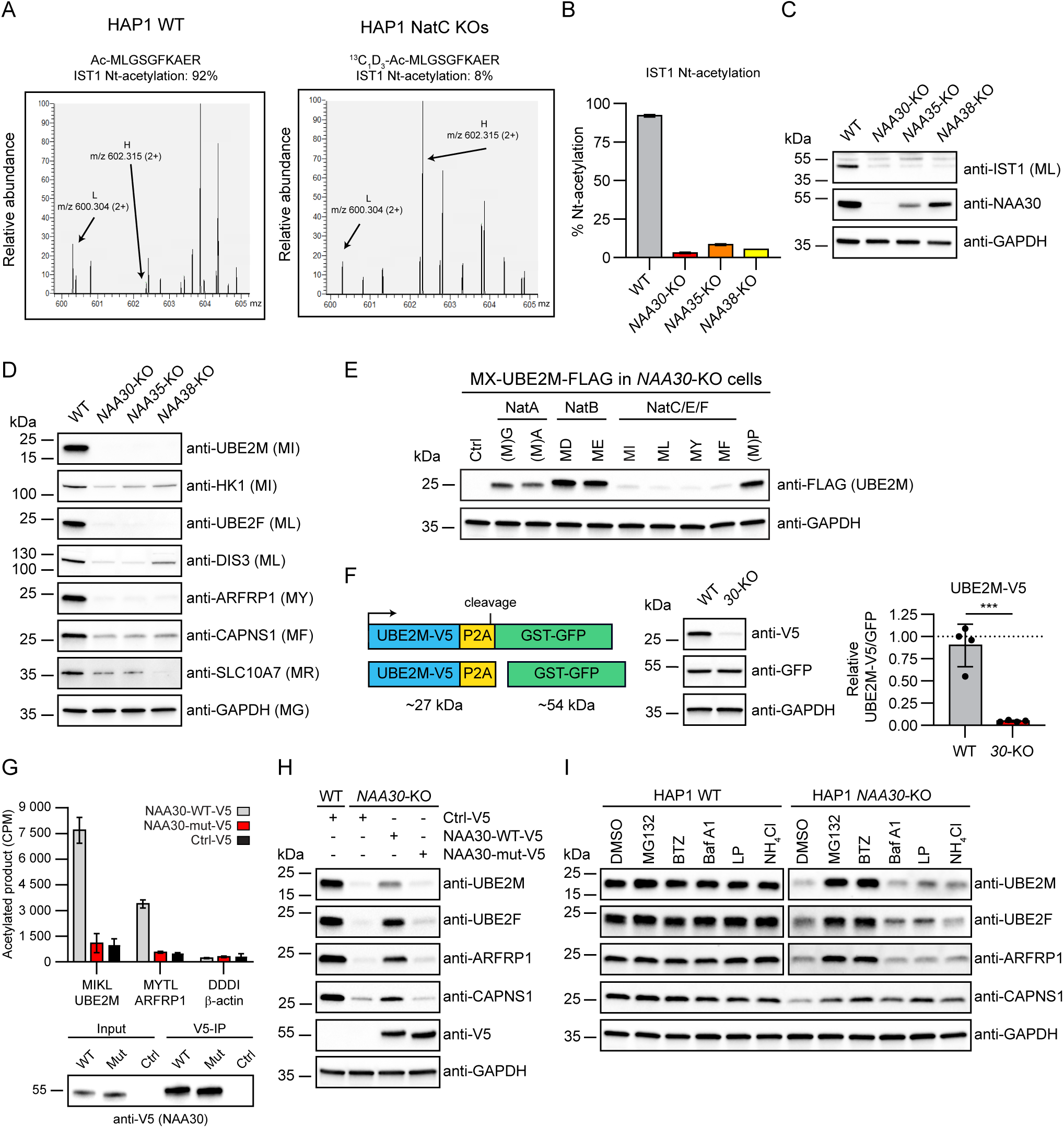
N-terminal acetylation by NatC protects proteins from degradation. **(A)** MS-spectra for the N-terminal peptide of endogenous IST1 (P53990) from HAP1 WT and NatC KO cells following trypsin digestion and strong cation exchange (SCX) enrichment. **(B)** Bar graph showing the degree of Nt-acetylation of IST1 in HAP1 WT and NatC KO cells as determined by proteomics. Data are shown as mean ± SD (n = 4). **(C)** Endogenous IST1 protein levels from indicated HAP1 cells determined by immunoblotting. **(D)** Immunoblot analysis of confirmed and putative NatC substrates using total cell extract from HAP1 WT and NatC KO cells. **(E)** N-terminal variants of UBE2M-FLAG were expressed in HAP1 *NAA30*-KO cells and protein levels were determined by immunoblotting. **(F)** HAP1 WT and *NAA30*-KO cells were transfected with the indicated UBE2M-V5-P2A-GST-GFP reporter construct, and protein levels were determined by immunoblot analysis. UBE2M-V5 levels were normalized to GST-GFP and expressed relative to WT sample. Data are shown as mean ± SD of four independent experiments. ***p < 0.0004; two-tailed unpaired t test. **(G)** NAA30-WT-V5 and NAA30-mut-V5 was immunoprecipitated from HeLa cell extracts and used in Nt-acetylation assays with [^14^C]-acetyl-CoA and synthetic peptides representing the NatC substrates UBE2M (MIKL) and ARFRP1 (MYTL), and the NAA80/NatH substrate β-actin (DDDI). The experiment was performed three independent times with three technical replicates each. Data of one representative setup is shown as mean ± SD. **(H)** NatC regulates the protein level of UBE2M, UBE2F ARFRP1, and CAPNS1. Immunoblot analysis of HAP1 WT and *NAA30*-KO cells transfected with control V5 plasmid, NAA30-V5 or the catalytically dead mutant NAA30-mut-V5. **(I)** HAP1 WT and *NAA30*-KO cells were treated with proteasomal (MG132 and bortezomib (BMZ)) and lysosomal inhibitors (bafilomycin A (BafA), leupeptin (LP) or ammonium chloride (NH4Cl)) for 6 h followed by immunoblot analysis using the indicated antibodies. DMSO served as vehicle control.

Based on these datasets, a series of potentially regulated NatC substrates were assessed by immunoblotting analysis using lysates from WT cells and the different NatC KO cells. All three NatC KO cell lines displayed downregulation of the NEDD8-conjugating enzymes UBE2M and UBE2F, DIS3, HK1, CAPNS1, SLC10A7, and the previously established NatC substrate ARFRP1 (**Figure 2D**) (Behnia et al., 2004; Starheim et al., 2017). These proteins are thus likely substrates of the trimeric NatC complex and depend on NatC for their stability.

### N-terminal sequence and Nt-acetylation status are determinants of protein degradation

To define a potential Nt-sequence dependency of the NatC KO induced protein level regulation, we expressed N-terminal variants of UBE2M-FLAG in HAP1 *NAA30*-KO cells and determined protein levels by immunoblotting. In these cells, the UBE2M WT protein (MI-) and other variants with NatC-type N-termini (ML-, MY-, MF-) were expressed at very low levels, while variants with N-termini representing lack of Nt-acetylation and lack of methionine (MP-, which is processed to P-), or NatA (MG-, MA-) or NatB (MD-, ME-) substrate N-termini, were present at higher levels suggesting that these provide a stabilizing effect (**Figure 2E**). To exclude any effects at the transcriptional or protein synthesis levels, HAP1 WT and *NAA30*-KO cells were transfected with a bicistronic P2A reporter vector encoding UBE2M-V5-P2A-GST-GFP. Normalized UBE2M-V5 protein levels were significantly lower in *NAA30*-KO cells compared to WT cells (**Figure 2F**). The observed effects of NatC KO on several proteins are likely to be driven by the direct lack of NatC-mediated Nt-acetylation. To exclude the involvement of non-catalytic roles of NatC subunits, we generated a catalytically dead mutant of NAA30 based on sequence analysis of evolutionarily conserved amino acid residues crucial for catalysis (**Figure S2A**) (Grunwald et al., 2020). While NAA30-WT was capable of Nt-acetylating both UBE2M and ARFRP1-derived synthetic peptides, as shown by an *in vitro* acetylation assay, the mutant NAA30-mut (E321A) was not, despite being normally expressed (**Figure 2G**). In *NAA30*-KO cells, re-expression of NAA30-WT, but not NAA30-mut was able to partially restore protein levels of UBE2M, UBE2F, ARFRP1 and CAPNS1 (**Figure 2H**). This both confirms that the observed effects are NAA30-specific and that the catalytic activity of NAA30 is essential for the regulation of these proteins. To confirm that the NatC-mediated protein regulation is at the level of protein degradation, HAP1 WT and *NAA30*-KO cells were treated with proteasomal (MG132 and bortezomib) and lysosomal (bafilomycin A, and leupeptin) inhibitors. Immunoblot analyses of cell lysates clearly demonstrated that UBE2M, UBE2F and ARFRP1 were all degraded via the proteasome in *NAA30*-KO cells while CAPNS1 might be degraded both via proteasomal and lysosomal degradation in *NAA30*-KO cells (**Figure 2I**). These data strongly suggest that NatC-mediated Nt-acetylation may directly and positively steer protein stability of its substrates.

### N-recognin UBR4-KCMF1 targets non-Nt-acetylated NatC substrates for degradation

To elucidate how non-Nt-acetylated NatC substrates are targeted for degradation, we perturbed various UBR-box E3 ligases, called N-recognins, which recognizes different types of protein N-termini (Tasaki et al., 2005; Tasaki et al., 2009) and examined the endogenous UBE2M protein levels by immunoblotting. Knockdown of UBR4, and to some extent also knockdown of UBR1 and UBR2, increased the protein levels of UBE2M in *NAA30*-KO cells (**Figure 3A**). A combination of UBR1 and UBR2 knockdown, and in particular a triple knockdown of UBR1, UBR2 and UBR4 strongly raised UBE2M protein levels. This combined effect is consistent with the finding that these UBRs have partially overlapping targets (Tasaki et al., 2009; Timms et al., 2019) The E3 ligase UBR4 was shown to physically interact with the potential E3 KCMF1 and the E2 UBE2A (RAD6) (Hong et al., 2015), suggesting that they act in a ubiquitin ligase complex. Both UBR4, KCMF1 and UBE2A were identified as positive GIs of NatC (**Figure 1D**). To assess whether this UBE2A-KCMF1-UBR4 complex was responsible for targeting non-Nt-acetylated UBE2M for degradation, we separately knocked down KCMF1 as well as UBE2A and its human paralogue UBE2B. Indeed, UBE2M protein levels were restored when depleting human cells for UBR4, KCMF1 or by simultaneous knockdown of UBE2A and UBE2B (**Figure 3B**). In agreement with the lack of Nt-acetylation as a determinant for UBR4-KCMF1-mediated UBE2M degradation, we were not able to observe any changes in UBE2M protein level when knocking down UBR4 or KCMF1 in HAP1 WT cells where NatC Nt-acetylation is intact (**Figure S3A**). Moreover, the knockdown experiments revealed mutually interdependent protein levels of UBR4 and KCMF1. Finally, the protein levels of the top positive GIs were also analyzed, to rule out differential activity in NatC KO cells relative to WT cells, and except from the putative NatC substrate RSPRY1 (MI-) none of these appeared to be markedly shifted in NatC KO cells (**Figure S3B**).

**Figure 3.**
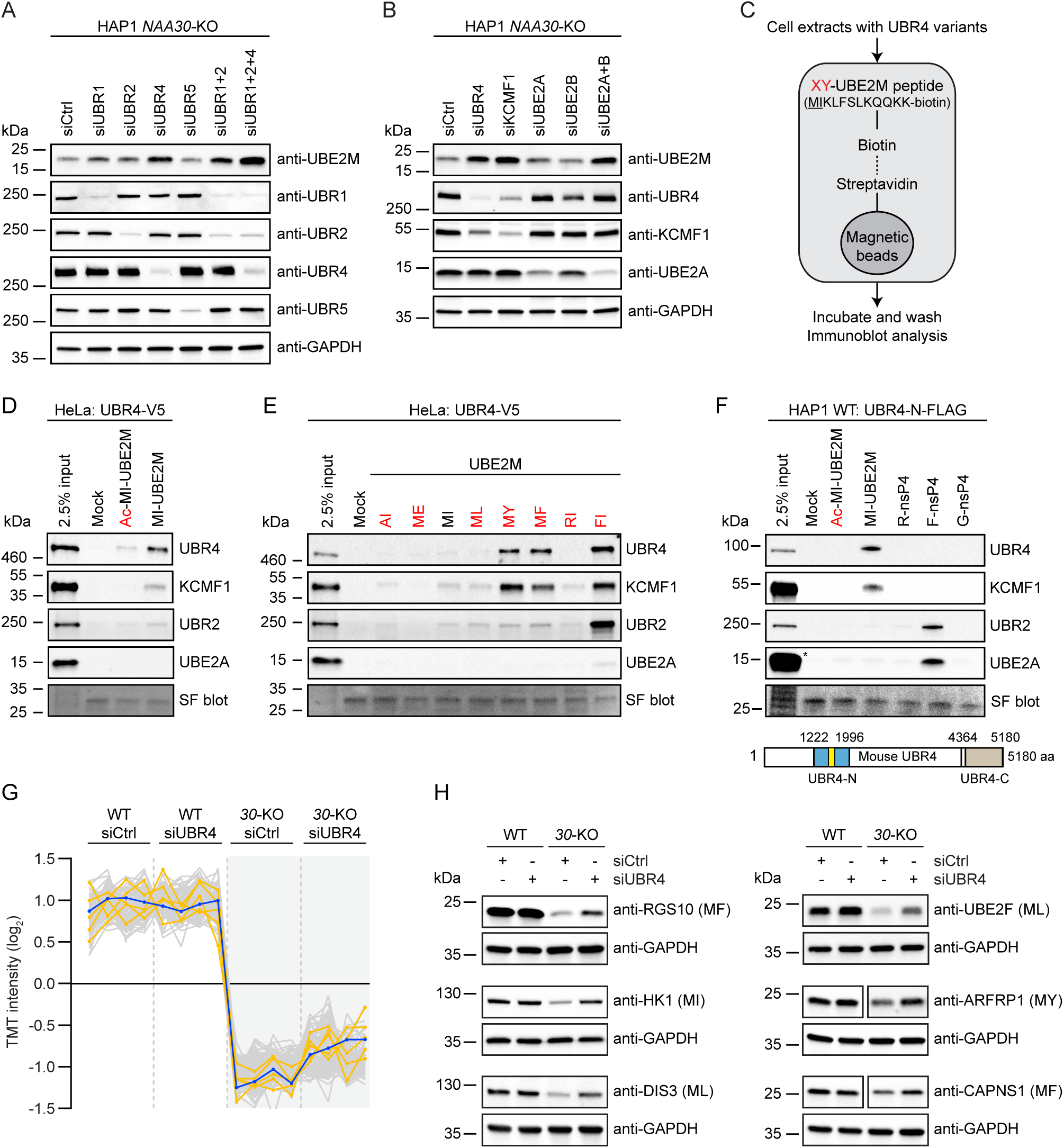
UBR4-KCMF1 targets unacetylated N-terminal methionine followed by a hydrophobic residue. **(A and B)** The protein level of endogenous UBE2M in HAP1 WT and *NAA30*-KO cells transfected with the indicated siRNAs for 72 h was assessed by immunoblotting. **(C)** Schematic representation of protein capture by peptide pulldown. A set of 11-mer peptides derived from the N-terminal sequence of UBE2M were C-terminally labelled with K-biotin (<u>MI</u>KLFSLKQQK(K-biotin)) and conjugated to streptavidin magnetic beads. The first two residues were replaced to represent different N-termini (XY-UBE2M). Biotinylated UBE2M peptides were incubated with cell extracts and the pulled-down proteins were identified by immunoblot analysis. (**D)** *In vitro* peptide pulldown assay of UBR4-V5 expressed in HeLa cells using acetylated and non-Nt-acetylated UBE2M peptide. **(E)** XY-UBE2M peptide pulldown assay with UBE2M peptides bearing different N-terminal amino acids and UBR4-V5 expressed in HeLa cells. **(F)** *In vitro* peptide pulldown assay of UBR4-N-FLAG expressed in HAP1 WT cells using acetylated and non-Nt-acetylated UBE2M peptide and X-nsP4 controls peptides. The UBR4-N construct contains the UBR-box (yellow), which is the substrate recognition domain of the UBR proteins. * indicates saturated UBE2A band. **(G)** HAP1 WT and *NAA30*-KO cells were transfected with siCtrl or siUBR4 for 72 h and protein abundance was determined by TMT-based quantitative proteomics (see **Table S5**). Intensity profile plot showing protein levels of the top 100 proteins with abundance profiles most similar to UBE2M (blue trace) (FDR = 0.01, S0 = 0.1). The intensity profiles of RGS10, ARLB, CAPSN1, DIS3, and HK2 are highlighted in orange. **(H)** UBR4 knockdown stabilizes the protein levels of non-Nt-acetylated RGS10, HK1, DIS3, UBE2F, ARFRP1 and CAPNS1 in *NAA30*-KO cells. HAP1 WT and *NAA30*-KO cells were transfected with siCtrl or siUBR4 for 72 h followed by immunoblotting using the indicated antibodies.

To directly assess the Nt-acetylation dependency of UBR4-KCMF1-mediated protein targeting, we performed a peptide pulldown assay using a 11-mer peptide derived from the N-terminal sequence of UBE2M and HeLa cells transiently transfected with full-length UBR4-V5 (**Figure 3C**). The non-Nt-acetylated UBE2M (MI-UBE2M) pulled out UBR4-V5, KCMF1 and UBR2 from cell lysates to a larger extent than Nt-acetylated UBE2M (Ac-MI-UBE2M) (**Figure 3D**). Also, all the non-Nt-acetylated UBE2M peptides with different NatC-type N-terminal sequences (MI/ML/MY/MF-UBE2M) pulled out UBR4-V5, KCMF1 and UBR2, while UBE2M peptides with a NatA-type (AI-UBE2M) or NatB-type N-terminus (ME-UBE2M) were impaired in this capacity (**Figure 3E**). Control UBE2M peptides bearing the classical Arg/N-degron N-terminal residues Arg (type 1) (RI-UBE2M) or Phe (type 2) (FI-UBE2M) (Tasaki et al., 2005) weakly and strongly captured these cellular E2-E3 components, respectively.

The UBR box is the substrate recognition domain of UBR1 and UBR2, and recognizes type 1 N-degrons (basic), while an additional segment, termed the N-domain, is required for type 2 (bulky hydrophobic) recognition (Tasaki et al., 2009). The UBR box of UBR4 most likely assumes the same fold as UBR1 and UBR2, but lacks a critical coordination residue and may therefore require additional structural elements for substrate recognition (Matta-Camacho et al., 2010). To further define the UBE2M-UBR4 interaction, we tested whether an N-terminal segment of mouse UBR4 containing the UBR box (UBR4-N) had the ability to bind to non-Nt-acetylated UBE2M peptide. We found that UBR4-N transiently expressed in HAP1 WT cells was specifically pulled out together with KCMF1 by non-Nt-acetylated UBE2M, but not by Nt-acetylated UBE2M (**Figure 3F**). In this case, the Phe-starting control peptide (F-nsP4), derived from a Sindbis virus polymerase (Degroot et al., 1991; Tasaki et al., 2005), pulled out UBR2, but not UBR4-N and KCMF1, while Arg- and Gly-starting control peptides (R-nsP4 and G-nsP4) did not pull out any of these components of the N-degron pathway. Furthermore, none of the peptides were able to pull out a C-terminal segment of UBR4 (data not shown) considered to be sufficient for ubiquitin ligation (Hunt et al., 2019; Lin et al., 2013). Taken together, our data suggest that N-termini of NatC-type substrates (MI, ML, MY, MF) are recognized by UBR4, KCMF1 and UBR2 when present in their non-Nt-acetylated state.

### Loss of UBR4 protects unacetylated hydrophobic N-termini from degradation

We have already shown that the stability of several proteins depends on the NatC complex and its catalytic activity (**Figure 2**), and that *UBR4* knockdown can reverse this effect for UBE2M in *NAA30*-KO cells (**Figure 3A**). To assess the potential interplay between NatC and UBR4-mediated protein degradation on a global level, we performed *UBR4* siRNA knockdown in HAP1 WT and in *NAA30*-KO cells followed by TMT-based quantitative proteomics. Several proteins were regulated depending on NatC and/or UBR4 status (**Figure S3C and Table S5**). We identified proteins with abundance profiles similar to UBE2M: displaying reduced protein levels in *NAA30*-KO as compared to WT cells and additionally increased protein levels in siUBR4-treated compared to siCtrl-treated *NAA30*-KO cells (**Figure 3G**) (FDR = 0.01, S0 = 0.1). Regulator of G protein signaling (RGS) proteins play crucial regulatory roles in G-protein-mediated signal transduction, and RGS2, RGS4, RGS5 and RGS16 are known *in vivo* substrates of the Arg/N-degron pathway (Hu et al., 2005; Lee et al., 2005; Park et al., 2015). Interestingly, we found that RGS10 (MF-) has the most similar abundance profile compared to UBE2M, indicating that yet another RGS protein might be targeted by the N-degron pathway. We also identified the NatC substrate ARL8B (Hofmann and Munro, 2006; Starheim et al., 2009), and several other proteins thought to be regulated by NatC, such as CAPNS1, HK1/2, and DIS3. These putative NatC substrates proteins were independently verified by immunoblot analyses to be negatively regulated by *NAA30*-KO and positively regulated by *UBR4*-knockdown (**Figure 3H**). The data suggest that many NatC substrates are shielded from UBR4-targeted degradation via Nt-acetylation.

### The principal molecular role for NatC-mediated Nt-acetylation is blocking protein degradation occurring via the N-degron pathway

Our finding that several components of the Arg/N-degron pathway, UBR4, KCMF1, UBR2, and UBE2A are strong positive GIs of the NatC subunits (**Figures 1D-F**), suggests that removal of this protein degradation system may eliminate NatC KO mediated negative effects on cell viability. Furthermore, our data show that several cellular proteins are regulated, in a Nt-acetylation and sequence dependent manner by NatC and N-recognins (**Figure 2**, **Tables S3-4**). Based on (i) Nt-acetylome analyses, (ii) the substrate specificity of human NatC, and (iii) proteome-wide estimations, up to 20% of the human proteome might be Nt-acetylated by NatC (Aksnes et al., 2016; Ree et al., 2018; Van Damme et al., 2016). This plethora of NatC substrates, many of which might need Nt-acetylation for stability, may imply that multiple cellular functions and pathways are regulated by the NatC complex. Indeed, we found that NatC KO cells displayed abnormalities in organelles and in the secretory pathway: abnormal mitochondrial morphology (**Figure 4A, Figure S4A**) and increased lysosomal content (**Figure 4B**). Organellar abnormalities were reflected in increased cellular granularity for all NatC KO cell lines (**Figure 4C**) and increased cellular dehydrogenase activity (**Figure S4B**). Furthermore, we found that NatC-mediated Nt-acetylation is important for UBE2M and UBE2F mediated cullin neddylation. CUL1, CUL4A and CUL5 were less neddylated in all NatC KO cell lines (**Figure 4D**) (the upper band represents neddylated cullins). In addition, the NatC KO cells had increased levels of the NEDD8 ligase RBX2, but the same effect was not observed for RBX1. Our proteomics analysis suggested that the autophagy receptor p62/SQSTM1 was upregulated in NatC KO cells (**Table S4**), which was corroborated at the single-cell level by immunofluorescence analysis (**Figure 4E**). Finally, we found by immunoblotting that the autophagy regulator BCL2 and the early endosome marker EEA1 were downregulated, while p62 was upregulated, but without an apparent activation of autophagy via LC3B detection (**Figure 4F**). We next wanted to define whether NatC’s ability to shield proteins from N-degron mediated protein degradation represents a general cellular NatC function. We depleted *NAA30*-KO cells for UBR1, UBR2, and UBR4 due to their overlapping effect on NatC substrates (**Figure 3A**). UBR knockdown fully or partially restored all tested cellular NatC KO phenotypes including increased p62, decreased CUL5 neddylation (**Figure 4G**), abnormal mitochondria morphology (**Figure 4H**), elevated presence of lysosomes (**Figure 4I**), and increased cellular granularity (**Figure 4J**). In summary, the combination of genetics data (**Figure 1**), cellular data (**Figures 2**-**3**) biochemical data (**Figure 3**), and phenotype data (**Figure 4**) show that the major molecular role of (NatC-mediated) Nt-acetylation is shielding proteins from the N-degron pathway.

**Figure 4.**
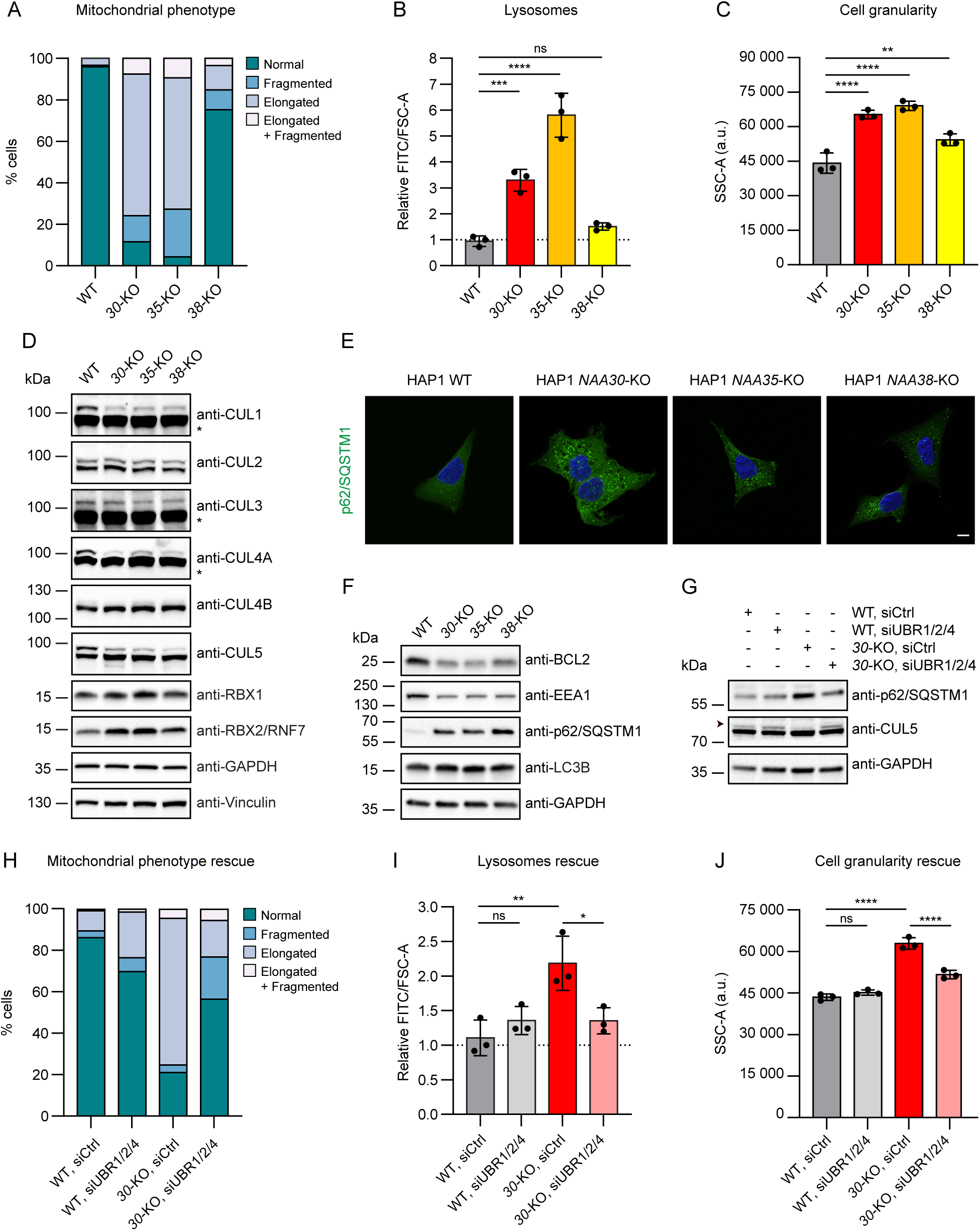
UBR-dependent NatC knockout phenotypes include abnormal mitochondrial morphology, increased lysosomal content, and increased cell granularity. **(A)** NatC KO cells have abnormal mitochondrial morphology. HAP1 cells were stained with the mitochondrial marker anti-COX IV and analyzed by immunofluorescence (IF). Cells were grouped into four bins based on mitochondrial morphology: normal, fragmented, elongated and elongated + fragmented. 100 cells per cell line were counted. **(B)** *NAA30*-KO and *NAA35*-KO cells have increased lysosomal content. Lysosomes were stained with LysoView 488 and analyzed by flow cytometry. Median FITC values were normalized to cell size (FSC-A) and expressed relative to WT sample. Data are shown as mean ± SD of three independent experiments. ***p < 0.001, ****p < 0.0001; one-way ANOVA with Dunnett’s correction. **(C)** NatC KO cells display increased granularity. Median side scatter area (SSC-A) indicating cell granularity or internal complexity was determined by flow cytometry. Data are shown as mean ± SD of three independent experiments. **p < 0.01, ****p < 0.0001; one-way ANOVA with Dunnett’s correction. (**D)** NatC regulates protein neddylation. Immunoblot analysis of several components of the neddylation pathway using total cell extract from HAP1 WT and NatC KO cells. The upper cullin band represents the neddylated form. * indicates saturated bands. **(E)** Increased p62/SQSTM1 levels in NatC KO cells. HAP1 cells were stained with antibody targeting p62 and analyzed by immunofluorescence. Scale bar, 10 μm. **(F)** NatC affects components of the autophagy pathway. Immunoblot analysis of the autophagy regulator BCL2, early endosome marker EEA1, autophagy receptor p62 and autophagy marker LC3B using the indicated HAP1 cell extracts. **(G-J)** HAP1 WT and *NAA30*-KO cells were transfected with siCtrl or siUBR1, siUBR2 and siUBR4 for 72 h. **(G)** Endogenous p62 and CUL5 protein levels were determined by immunoblot analysis. Arrowhead indicates NEDD8-CUL5. **(H)** Mitochondrial morphology was assessed by IF like in (A). **(I)** Lysosome levels were determined by Lysoview staining combined with flow cytometry like in (B) **(J)** Cell granularity (SSC-A) was determined by flow cytometry like in (C). **(I-J)** Data are shown as mean ± SD of three independent experiments. *p < 0.05, **p < 0.01, ****p < 0.0001; one-way ANOVA with Šídák’s correction. ns; not significant.

### Identification of the *Drosophila melanogaster* NatC complex

To further investigate the *in vivo* function of NatC we turned to the fruit fly *Drosophila melanogaster,* which provides an integrative model where genetics and behavioral tests can be combined. First, we performed a bidirectional BLAST search to identify the catalytic subunit of *Drosophila* NatC. We identified the protein sequence encoded by the gene *CG11412* as the most similar to human NAA30 and yeast Naa30 (Mak3), having an identity of 72.7% and 52.3%, respectively (**Figure S5A**). Supporting our analysis, *CG11412* was recently annotated by FlyBase as N(alpha)-acetyltransferase 30 A (Naa30A). A closely related, testis-specific paralog, Naa30B, was also annotated. To confirm that Naa30A is a functional orthologue of human NAA30 and yeast Naa30, we expressed Myc-tagged Naa30A in *Drosophila* embryos, using the UAS/Gal4 system and the nanos-Gal4 driver (Brand and Perrimon, 1993; Rorth, 1998; Van Doren et al., 1998). Immunoprecipitation with an anti-Myc antibody (**Figure S5B**) coupled to mass spectrometry analysis of co-immunoprecipitated proteins, identified the *Drosophila* orthologues of human and yeast NAA35/Mak10 (CG4065) and NAA38/Mak31 (CG31950), as interacting proteins of Naa30A (**Figure S5C, Table S6**). This confirmed that the NatC subunit composition is conserved between yeast, *Drosophila* and humans. In yeast, Golgi targeting of Arl3 requires NatC-mediated Nt-acetylation (Osberg et al., 2016; Setty et al., 2004). Interestingly, and further supporting functional conservation, ectopic expression of *Drosophila* Naa30A in yeast *naa30*-deletion cells rescued the Arl3 localization defect (**Figure S5D).** Altogether, our results suggest that NatC composition and substrate specificity is likely conserved between yeast, *Drosophila* and human cells.

### *Drosophila Naa30A* deletion flies are viable

To study the function of NatC in flies, we generated a genomic *Naa30A* deletion using p-element-mediated imprecise excision. We thus created a *Naa30A* mutant animal where almost all coding sequence of *Naa30A* gene was deleted (**Figure S6A**). This resulted in the deletion of the first 302 amino acids of Naa30A, with an additional frameshift mutation within the remaining coding sequence and a stop codon in the newly generated position 20. Surprisingly, deletion of *Naa30A* did not significantly impair adult viability (**Figure S6B**). However, and suggesting underlying phenotypes, *Naa30A* deletion flies had a striking “held out” wing phenotype, where adult animals keep their wings with a 90° angle from the axis of the body axis. This phenotype is typical of animals with muscular defects (Zaffran et al., 1997). We concluded that although NatC is likely to have hundreds of substrates, *Naa30A* is not essential for adult viability and most of *Drosophila* development. This is possibly due to partial redundancy with other related paralogue proteins, like Naa30B and CG31730, or even with NatE and NatF, which have previously been suggested to share overlapping substrate specificity (Van Damme et al., 2015; Van Damme et al., 2011b).

### *Drosophila* NatC is required for normal longevity and to prevent age-dependent motility loss

To further investigate the function of Naa30A we performed longevity assays (see STAR methods). We found a significant longevity reduction of the *Naa30A* deletion males (**Figure 5A**) (replica 1: median survival of control is 44 days after pupae eclosion vs 32 days of *Naa30A* deletion; replica 2: median survival of control is 42 days after pupae eclosion vs 25 days of *Naa30A* deletion). Importantly, the longevity defects of Naa30A deletion males could be fully rescued with a *Naa30A* genomic construct (genomic rescue) proving specificity (replica 1: median survival is 51 days after pupae eclosion; replica 2 median survival is 49 days) (**Figure 5A**).

**Figure 5.**
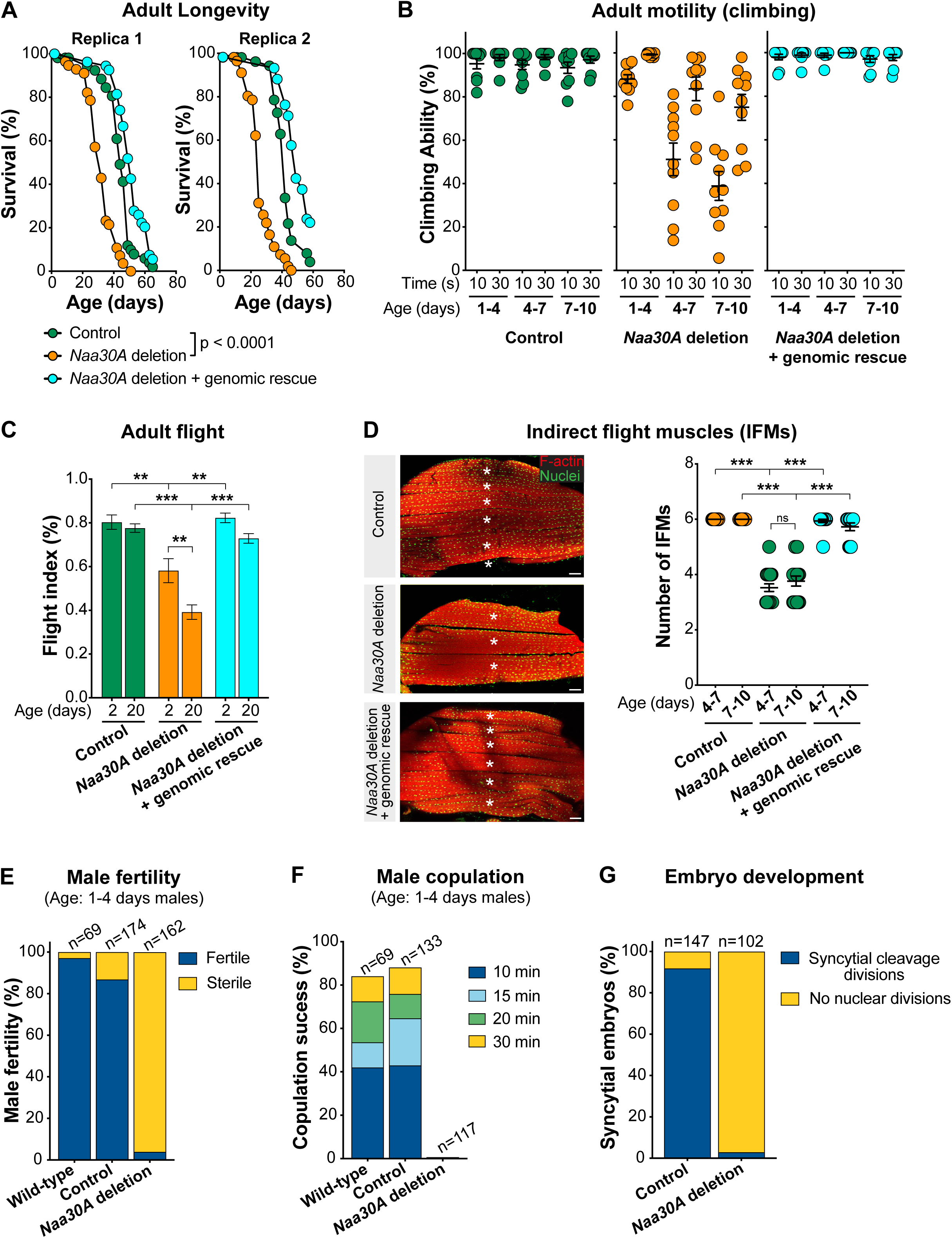
*Drosophila* Naa30A is required for normal longevity, fertility, and age-dependent motility. **(A)** Loss of Naa30A significantly reduces adult longevity. Survival curves of control males, *Naa30A* deletion males, and *Naa30A* deletion males carrying a *Naa30A* genomic rescue construct (two biological replicas). p<0.0001, log-rank (Mantel-Cox) test, n>50. **(B)** *Naa30A* deletion males show an accelerated age-dependent loss of motility. Climbing ability was used as a measure of mobility and represents the percentage of males that were able to climb 8 cm after 10 and 30 s. Control males, *Naa30A* deletion males, and *Naa30A* deletion males carrying a *Naa30A* genomic rescue construct at the ages of 1-4 days, 4-7 days, and 7-10 days after pupae eclosion were used. Results were obtained with males collected from two independent biological replicas. Each dot represents the average climbing ability of a group of 10 males after 10 repetitions (technical replicates), mean ± SEM is also indicated. The ability of very young *Naa30A* deletion males (1-4 days) to reach the 8 cm mark after 10 s is similar to control males. (p=0.8497; one-way ANOVA with Bonferroni’s correction), while the ability of slightly older *Naa30A* deletion males (4-7 or 7-10 days) to reach this mark after 10 s is significantly reduced (p<0.0001; one-way ANOVA with Bonferroni’s correction). **(C)** *Naa30A* deletion males show reduced flight ability. Flight ability of control males, *Naa30A* deletion males, and *Naa30A* deletion males carrying a *Naa30A* genomic rescue construct 2 or 20 days after pupae eclosion. Results are mean ± SEM of at least 41 males. **p<0.001, \*\*\**p*<0.0001; one-way ANOVA with Bonferroni’s correction. **(D)** *Naa30A* deletion males show developmental defects of flight muscles. <u>Left panels</u>: Sagittal sections of adult indirect flight muscles (IFMs) stained for F-actin (Phalloidin, red) and nuclei (Mef2, green). The number of IFMs decreases in *Naa30A* deletion and this phenotype was rescued by a genomic construct containing *Naa30A*. Scale bar is 50 μm. * marks individual dorsal longitudinal IFM. <u>Right panel</u>: Quantification of the number of IFMs in control, *Naa30A* deletion, and *Naa30A* deletion carrying a *Naa30A* genomic rescue construct at 4-7 and 7-10 days after pupae eclosion. Each dot represents the number of IFM per hemithorax. Results are mean ± SEM of least 10 thoraxes per condition. \*\*\**p*<0.0001; one-way ANOVA with Bonferroni’s correction. (**A-D**) Genotypes of the males used: control males (y^1^/Y), *Naa30A* deletion males (y^1^, Naa30A^Δ74^/Y) and *Naa30A* deletion males carrying a *Naa30A* genomic rescue construct (y^1^, Naa30A^Δ74^/Y;; gNaa30A-myc/+). **(E-G)** *Naa30A* deletion males show reduced fertility and significant copulation defects. **(E)** Male fertility measured by crossing single wild-type males, control males and *Naa30A* deletion males (all 1-4 days after pupae eclosion) with 5 wild-type virgins. Percentage of fertile males was assessed by the presence of progeny (egg hatching and larvae). Results were obtained with males collected from 2 to 4 independent crosses; n represents the total number of males tested. **(F)** Copulation success was measured by the percentage of wild-type, control and *Naa30A* deletion males that were able to initiate copulation with 5-7 days old wild-type adult female virgins (OR) within 10, 15, 20 or 30 min. All males were 1-4 days old after pupae eclosion. Results are the mean of 2 independent crosses and n represents the total number of males tested. **(G)** Percentage of developing embryos (syncytial nuclear divisions or later stages of development) from wild-type (OR) females crossed with 1-4 days (after pupae eclosion) control males or *Naa30A* deletion males. Results are the sum of two independent experiments and n represents the total number of embryos scored. (**E**-**G**) Genotypes of the males used: wild-type males (Oregon R; OR), control males (y^1^/Dp(1;Y)y^+^) and *Naa30A* deletion males (y^1^, Naa30A^Δ74^/ Dp(1;Y)y^+^), and wild-type virgins (OR).

Beyond a significant reduction of longevity, we also noticed that *Naa30A* deletion males were apparently less active than control flies. To investigate this, we performed a climbing behavior assay. *Drosophila* negative geotaxis locomotor behavior is well established and has been used to study, for example, aging and distinct degenerative disorders (Feany and Bender, 2000; Gargano et al., 2005; Jahn et al., 2011). While the ability of very young *Naa30A* deletion males (1-4 days; after pupae eclosion) to climb 8 cm in 10 seconds was similar to control males (**Figure 5B**), the climbing ability of slightly older *Naa30A* deletion males (4-7 or 7-10 days) was significantly reduced (**Figure 5B and Movie 1**). The climbing defects of *Naa30A* deletion males were fully rescued by a *Naa30A* genomic construct (**Figure 5B**). Similar results could also be observed for female flies (data not shown). Altogether this suggests that NatC is required to prevent age-dependent loss of motility.

Consistent with the “held out” adult wing phenotype of *Naa30A* deletion males, we also observed an age-dependent reduction of adult flight ability (**Figure 5C**). The flight ability defects could also be fully rescued with the *Naa30A* genomic rescue (**Figure 5C**). To investigate if the age-dependent loss of motility was associated with muscle defects, we decided to analyze the *Drosophila* indirect flight muscle (IFM). Consistently, we found that *Naa30A* deletion males show a significant reduction in the number of dorsal longitudinal IFM (**Figure 5D**). Reassuringly, this phenotype was fully rescued by the *Naa30A* genomic rescue (**Figure 5D**). The reduced number of adult dorsal longitudinal IFMs suggests defects in the splitting of the three sets of larval muscle fibers during metamorphosis, an event required to form the six dorsal longitudinal IFM fibers per hemisegment in adults (Gunage et al., 2017). Taken together, these data suggest that the observed age-dependent motility phenotypes are partially due to underlying developmental defects of the muscles, which increasingly impaired motility with age.

### *Drosophila* NatC is required for male copulation and fertility

Given the underlying muscle defects induced by *Naa30A* deletion, we investigated other motility phenotypes beyond climbing and flight ability. We observed that *Naa30A* deletion males were sterile, even when matted without male competition with highly receptive aged virgin females (**Figure 5E**). The use of aged virgin females, in the absence of male competition (e.g., single male crosses), mitigates the relevance of potential courtship defects for male sterility. To examine if copulation was normal, we measured copulation success of *Naa30A* deletion males and found that they did not copulate even in the presence of receptive aged virgin females (**Figure 5F**). Consistent with the copulation defects, the eggs laid by females mated with *Naa30A* deletion males did not initiate development (**Figure 5G**). We therefore concluded that *Naa30A* deletion males are sterile due to an inability to copulate. Such defects are likely to be due to musculature abnormalities of the male genitalia and terminalia involved in copulation.

### *Drosophila* NatC is seemingly not rate-limiting for mitochondrial function

NatC perturbation is associated with significant mitochondrial network defects in human cells (**Figures 4A, S4A**) (Van Damme et al., 2016). Since such defects could easily explain the motility and longevity phenotypes of *Drosophila Naa30A* deletion males, we decided to investigate if mitochondria were normal in the IFM muscles. We failed to detect obvious morphological defects of the mitochondrial network within muscles from *Naa30A* deletion males (**Figure S7A**). However, we detected a small reduction in the total levels of ATP in the absence of Naa30A (**Figure S7B**). Altogether, this suggests that although mitochondrial defects can potentially be a contributor factor to the observed *Naa30A* deletion phenotypes, they are not likely to be the main cause.

### *Drosophila* NatC is required to maintain muscle proteostasis

Age-dependent loss of muscle function has been associated with proteostasis defects and the accumulation of protein aggregates in *Drosophila* and humans (Ayyadevara et al., 2016; Demontis and Perrimon, 2010). As expected, we observed by, using an antibody specific for mono- and polyubiquitinylated conjugates, an age-dependent increase in aggregate-like structures in control muscles (**Figures 6A-C**). Remarkably, analysis of young and old *Naa30A* deletion males showed a dramatic increase in the accumulation of polyubiquitinylated aggregates (**Figures 6A-C**), clearly suggesting an important role of Naa30A in maintaining muscle proteostasis. However, since the total amount of protein aggregates were already high in young flies’ muscles, and this level did not significantly increase with age (**Figure 6C**), we reasoned that they were not likely to be directly related with the age-dependent motility defects of *Naa30A* deletion males. An alternative explanation would be that although young muscles have high levels of aggregates, older muscles are significantly more susceptible to such aggregates.

**Figure 6.**
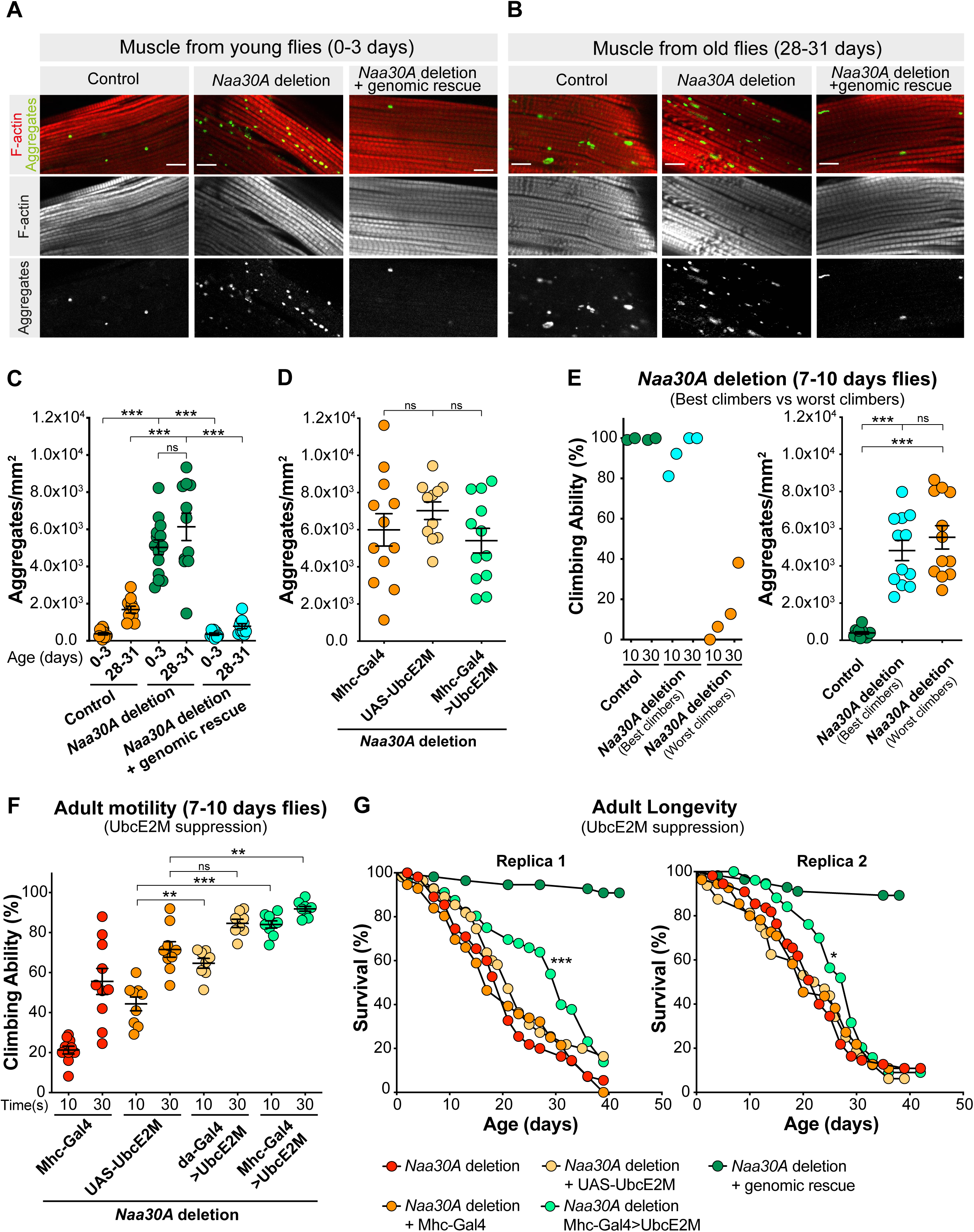
Overexpression of UbcE2M in the muscles suppresses the longevity and motility defects of *Drosophila Naa30A* deletion. **(A-B)** *Naa30A* deletion males show an accelerated accumulation of aggregate-like polyubiquitinated protein structures in muscles. Indirect flight muscles from **(A)** young (0-3 days after pupae eclosion) and **(B)** old (28-31 days after pupae eclosion) control males, *Naa30A* deletion males, and *Naa30A* deletion males carrying a *Naa30A* genomic rescue construct were immunostained with phalloidin-TRITC to visualize muscle actin (red) and anti-polyubiquitin (green). Scale bar is 10 μm. **(C)** Quantification of the aggregate-like polyubiquitinated protein structures (aggregate/mm^2^) as shown in (A and B). **(D)** Overexpression of UbcE2M does not suppress the accumulation of aggregate-like structures in *Drosophila Naa30A* deletion males. Quantification of the aggregate-like polyubiquitinated protein structures (aggregate/mm^2^) in *Naa30A* deletion males carrying the muscle-specific Gal4 driver Mhc-Gal4, the UAS-UbcE2M-HA construct, or overexpressing UAS-UbcE2M-HA in muscles. **(C and D)** Results were obtained with males collected from two independent crosses and each dot represents the aggregate density in a muscle fiber of a different male. Results are mean ± SEM is also shown. “ns” denotes not significant, \*\*\**p*<0.0001; one-way ANOVA with Bonferroni’s correction. **(E)** No correlation between the motility defects and the density of aggregate-like structures in *Drosophila Naa30A* deletion males. <u>Left panel</u>: climbing ability of 11 randomly selected control males, 9 *Naa30A* deletion males with good climbing ability (best climbers) and 11 *Naa30A* deletion males with poor climbing ability (worst climbers). Climbing ability represents the percentage of males that were able to climb 8 cm after 10 and 30 sec. Results are mean ± SEM with 10 repetitions (technical replicates). <u>Right panel:</u> quantification of the aggregate-like polyubiquitinated protein structures (aggregate/mm^2^) present in indirect flight muscles from 11 randomly selected control males, 9 *Naa30A* deletion males with good climbing ability (best climbers), and 11 *Naa30A* deletion males with poor climbing ability (worst climbers) previously used in the climbing assay (left panel). Results are from two different biological replicas and mean ± SEM of aggregate density is indicated. “ns” denotes not significant, \*\*\**p*<0.0001; one-way ANOVA with Bonferroni’s correction. The age of all males was 7-10 days after pupae eclosion. The best and worst climbers were selected from a group of 200 *Naa30A* deletion males. **(F)** Overexpression of UbcE2M in the muscles suppresses the motility defects of *Drosophila Naa30A* deletion. Climbing ability as a measure of mobility of *Naa30A* deletion males carrying the muscle-specific Mhc-Gal4 driver, carrying the UAS-UbcE2M-HA construct, overexpressing UAS-UbcE2M-HA ubiquitously, or overexpressing UAS-UbcE2M-HA specifically in the muscles. The age of all males was 7-10 days after pupae eclosion. Results were obtained with males collected from two independent crosses. Each dot represents the average climbing ability of a group of 10 males assessed with 10 repetitions (technical replicates). Mean ± SEM is indicated. **p<0.001 and ***p<0.0001; one-way ANOVA with Bonferroni’s correction. **(G)** Overexpression of UbcE2M in the muscles partially suppresses the adult longevity defects of *Drosophila* Naa30A deletion. Two survival curves (biological replicates) of *Naa30A* deletion males carrying the muscle-specific Mhc-Gal4 driver, carrying the UAS-UbcE2M-HA construct, overexpressing UAS-UbcE2M-HA in muscles, or carrying a *Naa30A* genomic rescue construct. Survival of *Naa30A* deletion males was significantly improved by UbcE2M overexpression in muscles in both biological replicates. *p<0.05, ***p<0.001, log-rank (Mantel-Cox) test, n>50).

To better understand this conundrum, we decided to compare the best and worse climbers within similarly aged *Naa30A* deletion males. For this purpose, we collected 200 *Naa30A* deletion males with 7-10 days of age after pupae eclosion. From these, we evaluated their climbing ability and selected the 9 males with best climbing ability (best climbers) and the 11 males with worst climbing ability (worst climbers). If aggregate accumulation is directly related to the observed age-dependent motility defects, then the best climbers should have less aggregates when compared to the worst climbers of a similar age and identical genetic background. We failed to detect any significant difference in the amount of protein aggregates within the muscles of the best and worst climbers of *Naa30A* deletion males (**Figure 6E**). This suggests that although aggregate accumulation is suggestive of muscle proteostasis defects, they are however not causal for the motility defects of *Naa30A* deletion.

### Overexpression of UbcE2M in the muscles suppresses the longevity and motility defects of *Drosophila Naa30A* deletion

Since the accumulation of protein aggregates was not likely causal for the motility defects induced by *Naa30A* deletion, and since we showed that several human NatC substrates are protected from degradation by Nt-acetylation (**Figure 2**), we hypothesized that these phenotypes instead resulted from the ability of NatC to protect a subset of proteins from protein degradation. Among the most relevant NatC substrates found in this study are the NEDD8-conjugating enzymes UBE2M and UBE2F, which control neddylation of several Cullins (Huang et al., 2009). Human UBE2M and UBE2F have partially overlapping roles (Enchev et al., 2015), while in *Drosophila* there is apparently only one homologue, UbcE2M, fulfilling the functions. Interestingly, the N-terminal sequence of *Drosophila* UbcE2M is highly similar to the human UBE2M (**Figure S2B**), which suggests that it is very likely to be a NatC substrate.

To investigate if *Drosophila* UbcE2M is related to *Naa30A* deletion phenotypes, we overexpressed UbcE2M, using the UAS/Gal4 system (Brand and Perrimon, 1993; Rorth, 1998) and the da-Gal4 driver (Wodarz et al., 1995), for ubiquitous expression, and Mhc-Gal4 driver (Schuster et al., 1996) for specific expression in the skeletal muscle. Supporting the role of *Naa30A* in protein degradation protection and the central role of UbcE2M in *Naa30A* deletion phenotypes, we observed that ubiquitous and muscle-specific overexpression of UbcE2M was sufficient to fully rescue the age-dependent climbing defects of *Naa30A* deletion (**Figure 6F**). Interestingly, protein aggregate accumulation in the muscles was not reduced by UbcE2M overexpression (**Figure 6D**), which supported our conclusion that these aggregates are not causal for the motility defects of *Naa30A* deletion males. Remarkably, the muscle-specific overexpression of UbcE2M also partially rescued the longevity defects of *Naa30A* deletion (**Figure 6G**), which highlights the importance of *Naa30A*-dependent muscle protein degradation protection and muscle proteostasis for longevity and age-dependent motility.

## DISCUSSION

Modifications have major impacts on the N-terminal landscape of proteins, with some being mutually exclusive while others play in concert to influence key biological functions. Nt-acetylation catalyzed by the NATs is one of the most common protein modifications in eukaryotes, whose general role remains enigmatic. Mutations in the NATs can cause rare disorders, resulting in developmental delay, intellectual disability, and congenital heart defects (Cheng et al., 2018; Morrison et al., 2021; Muffels et al., 2021; Rope et al., 2011; Ward et al., 2021). The molecular framework and consequently the mechanistic basis for how the NATs manifest in cellular and organismal phenotypes remains however largely unknown. One of the major current challenges is to identify which critical proteins NATs work on and mechanistically define why acetylation is important for the proper functioning of these proteins.

Here, we have performed unbiased genome-wide CRISPR KO screens to uncover genetic vulnerabilities related to Nt-acetylation. Our data revealed a strong interaction between the NatC complex and components of the Arg/N-degron pathway, highlighting a cellular system where Nt-acetylation protects against protein degradation. This study thus sheds new light on the intricate interplay between protein Nt-acetylation and the N-degron pathways, where the acetyl group can both promote or prevent selective protein degradation. For instance, Nt-acetylation may conditionally target proteins for degradation by the Ac/N-degron pathway, thereby playing a key role in protein quality control by removing improperly folded proteins and ensuring subunit stoichiometry (Hwang et al., 2010; Park et al., 2015; Shemorry et al., 2013). At the same time, non-Nt-acetylated proteins, including those starting with methionine, may be degraded via the Arg/N-degron pathway in yeast (Kim et al., 2014). A comprehensive mapping of yeast N-degrons indicated that the hydrophobic character is the predominant feature of N-degrons, not Nt-acetylation (Kats et al., 2018). A direct and general impact on protein stability was also not observed when comprehensively assessing NatA and NatB deletion yeast strains instead distinct cellular functions for the two major NATs were found towards gene regulation/genome integrity and protein folding, respectively (Friedrich et al., 2021). The complexity is further stressed by yeast NatA deletion impacting proteostasis via several different routes involving the Hsp90 chaperone and its client proteins (Oh et al., 2017), increased proteasomal activity via Rpn4 regulation, as well as increased activity of several ubiquitin ligases (Kats et al., 2022). In human cells, a fraction of non-Nt-acetylated NatA substrates can be targeted by IAP E3 ubiquitin ligases for proteasomal degradation (Mueller et al., 2021). Finally, NatA was shown to affect global protein turnover in *Arabidopsis*, via currently unknown N-recognin(s) (Linster et al., 2022). In the current study, we not only show that unacetylated methionine-hydrophobic-starting N-termini are physically recognized by the Arg/N-degron pathway but find that the cellular phenotypes induced by NatC KO can be fully reversed by removal of specific N-recognins, revealing a principal role for Nt-acetylation in shielding proteins from degradation in human cells.

One of the strongest positive GIs identified in our *NAA30* and the *NAA35* screens was the N-recognin UBR4 (Tasaki et al., 2009). Unlike other N-recognins, UBR4 appears to lack a classical ubiquitylation domain (Tasaki et al., 2005), suggesting that its modus operandi might deviate from that of canonical N-recognins, and possibly requires other components of the N-degron pathway. Intriguingly, UBR4 was shown to form a complex with the potential E3 ligase KCMF1 (Heo et al., 2021; Jang, 2004) and the E2 ubiquitin-conjugating enzyme UBE2A (Hong et al., 2015), and both have a strong positive GI with *NAA30* and *NAA35.* The UBE2A-KCMF1-UBR4 complex may target substrates for proteasomal and/or lysosomal-mediated degradation (Hong et al., 2015; Hunt et al., 2021; Kim et al., 2018; Tasaki et al., 2013). In this study, we showed that non-Nt-acetylated NatC substrates are targeted for degradation by UBR4-KCMF1.

The NatC-UBR sensitive phenotypes are most likely steered by several NatC substrates impacting vesicle trafficking, mitochondria, endosomes and lysosomes (**Figure 7**). Here, we studied selected confirmed and putative NatC substrates. Calpain small subunit 1 (CAPNS1, MF-) regulates the activity of calpain 1 and calpain 2 (CAPN1/2), two non-lysosomal cysteine proteases (Demarchi and Schneider, 2007). We found that both CAPNS1 and CAPN1 (MS-) were downregulated in NatC KO cells (**Table S4**), indicating a role for NatC-mediated acetylation in subunit quality control (Shemorry et al., 2013) where loss of CAPNS1 results in decreased CAPN1 protein levels. ARFRP1 (MY-) is a small GTPase regulating *trans*-Golgi trafficking and it is essential for glucose and lipid metabolism (Hesse et al., 2012; Hommel et al., 2010). Both human ARFRP1 and the yeast orthologue Arl3 have negative GI with the NatC genes, supporting an evolutionarily conserved role for NatC in the ARL3-ARL1-SYS1 axis (Behnia et al., 2004; Osberg et al., 2016; Setty et al., 2004). The two NEDD8-conjugating enzymes UBE2M (MI-) and UBE2F (ML-) have distinct substrate specificities. UBE2M interacts with the RING E3 ligase RBX1 while UBE2F pairs with RBX2 to regulate neddylation of CUL1-4 and CUL5, respectively (Huang et al., 2009). Cullin neddylation is mediated through a dual E3 mechanism involving a co-E3 ligase called DCN1, which accelerates UBE2M/UBE2F binding (Kim et al., 2008; Kurz et al., 2008; Scott et al., 2010). Nt-acetylation of UBE2M and UBE2F serves as an avidity enhancer by allowing burial of the N-terminus into a hydrophobic pocket of DCN1 (Monda et al., 2013; Scott et al., 2017; Scott et al., 2011). The key role of Nt-acetylation of UBE2M and UBE2F is stressed by the fact that the longevity and motility defects of the *Drosophila Naa30A* deletion males could partially be complemented by overexpression of UbcE2M, the only *Drosophila* UBE2M homolog (**Figure 5**). Considering that UBR knockdown increased UBE2M/UBE2F protein levels and restored neddylation of CUL5 in *NAA30*-KO cells (**Figure 4**), the degradation of UBE2M/UBE2F may be a crucial molecular event following lack of its Nt-acetylation, not merely the impaired UBE2M-DCN1 complex. These two events are likely connected: non-Nt-acetylated UBE2M will be more dissociated from its partner DCN1 thereby exposing its free N-terminus to the UBRs which then target UBE2M for degradation. Still, the finding that UBR knockdown restores cullin neddylation in *NAA30*-KO cells (**Figure 4**) suggests that UBE2M/UBE2F still form active neddylation complexes even in the absence of Nt-acetylation. Interestingly, UBE2M and UBE2F can crosstalk with each other, where UBE2M can promote ubiquitination and degradation of UBE2F under both normal and stressed conditions (Zhou et al., 2018). Nt-acetylation is an irreversible modification, and its permanency suggests a role in proteostasis rather than signal transduction. In yeast, global Nt-acetylation levels are generally unaffected by prolonged starvation (Varland et al., 2018a), but fluctuations in cellular acetyl-CoA levels may affect Nt-acetylation and apoptotic fate in human cancer cells (Yi et al., 2011). Further research is necessary to fully understand the effects NatC and Nt-acetylation have on protein neddylation under normal and stressed conditions.

**Figure 7.**
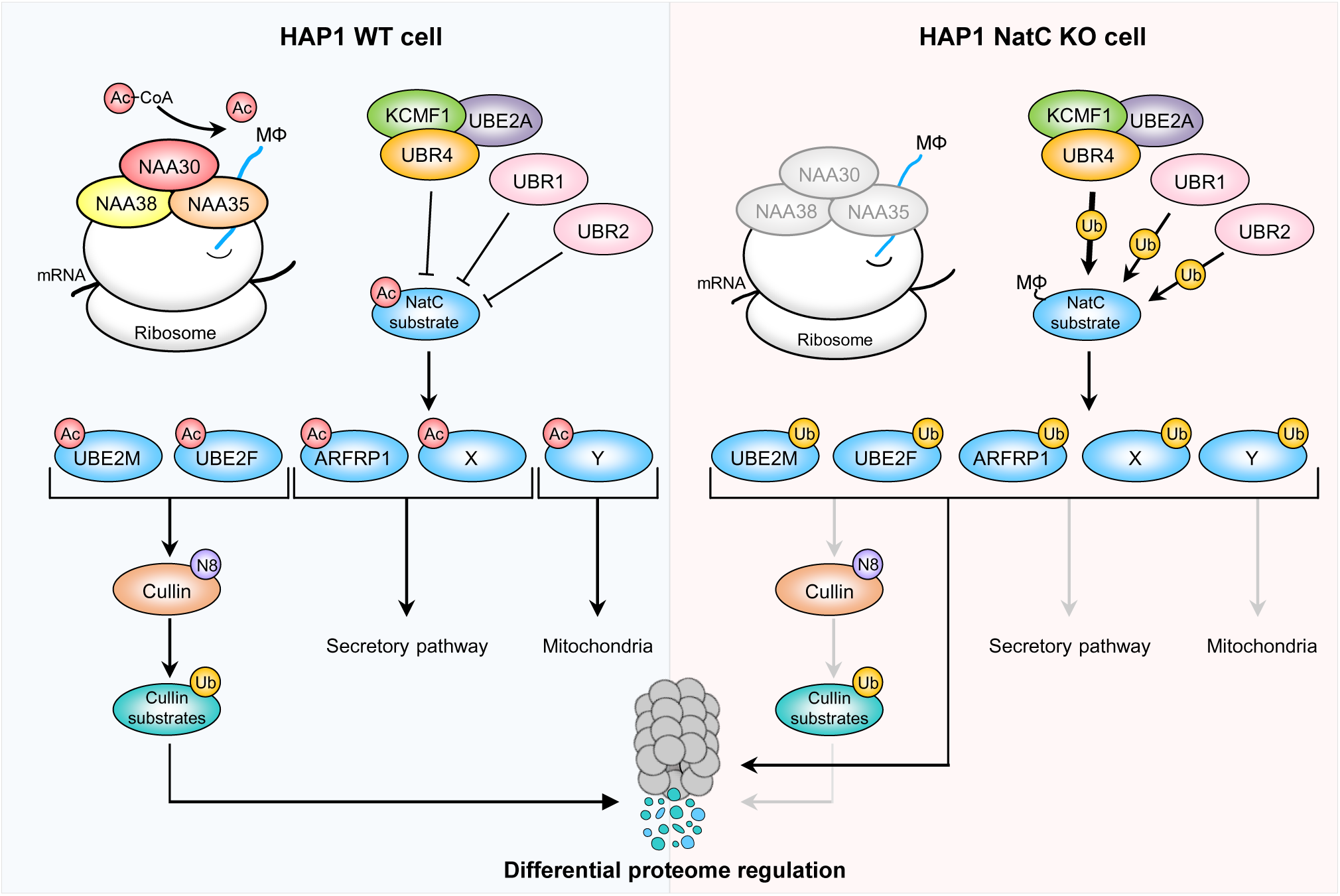
N-terminal acetylation by NatC shields proteins from degradation by preventing N-recognin UBR4-KCMF1 targeting. (**Left**) The NatC complex co-translationally acetylates proteins harboring a hydrophobic residue in the second position (MΦ-). Following acetylation, the NEDD8 E2 ligases Ac-UBE2M and Ac-UBE2F promote cullin neddylation, resulting in proteasomal degradation of targeted substrates, Ac-ARFRP1 is targeted to the Golgi where it plays a role in the secretory pathway, while the hypothetical proteins Ac-X and Ac-Y are thought to affect the secretory pathway and mitochondria, respectively. (**Right**) Loss of NatC leads to proteasomal and, in some cases, lysosomal degradation of non-Nt-acetylated substrates primarily via the Arg/N-recognin UBR4-KCMF1 and to some extent via UBR1 and UBR2. Targeted degradation of unacetylated non-Nt-acetylated NatC substrates leads to decreased cullin neddylation, increased mitochondrial elongation and fragmentation, and is thought to affect intracellular trafficking.

Decline of human and *Drosophila* muscle function precedes most age-related changes and has been associated with defective proteostasis and the progressive accumulation of protein aggregates (Demontis and Perrimon, 2010; Demontis et al., 2013; Nair, 2005; Zheng et al., 2005). The contribution of proteostatic defects to the loss of muscular function remains however mechanistically poorly understood. Here, we showed that *Drosophila* NatC is required for longevity, motility, and normal development of muscles, as deletion of *Naa30A* is associated with decreased lifespan, lower climbing ability, male copulation defects and a reduced number of adult IFM fibers (**Figure 5**). Supporting our hypothesis that one of the critical organismal functions of *Drosophila* NatC is the regulation of protein stability, *Naa30A* deletion leads to a dramatic increase in the accumulation of protein aggregates in the muscles, whereas muscle-specific overexpression of UbcE2M suppressed the decreased longevity and motility defects induced by this deletion (**Figure 6**). Interestingly, and strongly suggesting that NatC’s effect on the cellular repertoire of the UBR4-KCMF1 complex is highly conserved among metazoans, *Drosophila* Ubr4/Poe also regulates muscle function and lifespan, with reduced adult flying and climbing activity (Hunt et al., 2021; Hunt et al., 2019).

Loss of Ubr4 induces muscle hypertrophy, through decreased ubiquitination and degradation of a core set of target proteins (Hunt et al., 2021; Hunt et al., 2019). Consistently, muscle-specific knockdown of Ubr4 increased the number of poly-ubiquitinated protein aggregates, as Ubr4 directly associates with and increases the proteolytic activity of the proteasome. It was therefore suggested that these muscle defects most likely arose from reduced proteasome-mediated proteolysis and not via the autophagy-lysosome system (Hunt et al., 2021). However, the relationship between protein aggregates accumulation and loss of muscular function remains unclear, as most available evidence is correlative without a clear link to causality. The fact that we failed to detect a clear correlation between the amount of protein aggregates and the degree of climbing fitness among *Naa30A* deletion flies with similar age suggests no phenotypic causality. Furthermore, and significantly, overexpression of UbcE2M in the muscles was sufficient to rescue adult motility defects, without reducing protein aggregate accumulation in the muscles. This clearly demonstrates that although NatC regulation of the UBR4-KCMF1 substrates is critical to muscular function, the functional relevance of protein aggregate accumulation, at least in this context, is less clear. Altogether, these results suggest that *Drosophila* Naa30A deletion phenotypes can be explained by increased degradation of a subset of UBR4 target proteins due to a lack of NatC-mediated acetylation.

In conclusion, using an unbiased and global genetic screen, we uncovered a general function for NatC-mediated Nt-acetylation in protecting proteins from degradation in human cells. Our molecular investigations defined the ubiquitin ligases UBR4-KCMF1 and partially UBR1/UBR2 as responsible for degrading a major class of human proteins when lacking Nt-acetylation. The role of NatC-mediated protection of enzymes involved in cullin neddylation is evident both in human cells and in *Drosophila*. The impact of these pathways on longevity and motility in aged individuals underscores the vital role of protein Nt-acetylation.

### Limitations of the study

This study defines several new human NatC substrates, a role of Nt-acetylation in shielding proteins from N-recognins, and a direct involvement of specific N-degron pathway components in degrading NatC substrates when unacetylated. We could expand these investigations by testing more NatC substrates towards a broader panel of N-recognins (Sherpa et al., 2022). The dependency of *Drosophila* NatC deletion phenotypes on these UBRs could be investigated by muscle-specific depletion of different UBRs in NatC deletion *Drosophila* strains. Although the NatC enzyme appears to be fully conserved in eukaryotes in terms of subunits and substrate specificity, our study could be expanded by proteomic analyses to define *in vivo Drosophila* substrates by comparing the degree of Nt-acetylation between WT and *Naa30A* deletion flies.

## ACKNOWLEDGEMENTS

We thank members of the Arnesen, Martinho, Boone, and Andrews labs for helpful discussions. Lena Thurnes, Kjellfrid Haukenes, Nina Glomnes, Olga Sizova, and Andrea Habsid are gratefully acknowledged for technical assistance. We also thank Dr. Jaakko Saraste and Dr. Marc Niere for help and advice. Furthermore, we thank Dr. Scott J. Dixon for cell lines, Dr. Eileen Furlong for antibody, as well as Dr. Yong Tae Kwon, Dr. Fabio Demontis, and Dr. Liam C. Hunt for kindly providing plasmids. Next-generations sequencing was performed at the Centre for Applied Genomics, The Hospital for Sick Children, Toronto, Canada. Proteomics work of human cells lines was performed at the VIB Proteomics Core, located at the VIB-UGent Center for Medical Biotechnology, Belgium, and was supported by EPIC-XS, project number 823839, funded by the Horizon 2020 programme of the European Union. Proteomics work of *Drosophila* purified proteins was performed at the Mass Spectrometry Laboratory, Institute of Biochemistry and Biophysics, Polish Academy of Sciences, Warsaw, Poland. Sequencing was performed at Haukeland University Hospital. Flow cytometry analysis and cell sorting were performed at the Flow Cytometry Core Facility, Department of Clinical Science, University of Bergen, Norway, with advice from Dr. Brith Bergum. Confocal imaging was performed at the Molecular Imaging Center (MIC), Department of Biomedicine, University of Bergen.

This work was supported by grants from the Research Council of Norway (RCN) (FRIPRO Mobility Grant 261981 to S.V. which was co-funded by the European Union’s Seventh Framework Programme under Marie Curie grant agreement no 608695), the Norwegian Health Authorities of Western Norway (F-12540 to T.A.), the Norwegian Cancer Society (171752-PR-2009-0222 to T.A), the European Research Council (ERC) under the European Union Horizon 2020 Research and Innovation Program (Grant 772039 to T.A), the Research Foundation - Flanders (FWO) (G008018N and G002721N to K.G.), the French National Centre for Scientific Research (CNRS) (ATIP-Avenir grant to H.B), AFM-Telethon (Trampoline grant 23108 to H.B.), the Medical Research Council (MRC) (MC_UU_00028/6 to A.J.W), the Canadian Institutes of Health Research (CIHR) (FDN-143264 and FDN-143265 to C.B. and B.J.A.; PJT-180285 to C.B, PJT-463531 to J.M.), the National Institutes of Health (NIH) (R01HG005853 to B.J.A., C.B., and C.L.M. R01HG005084 to C.L.M.), and the Portuguese national funding through Fundação para a Ciência e a Tecnologia (FCT) (DL 57/2016/CP1361/CT0019 to R.D.S; PTDC/BIA-BID/28441/2017 and PTDC/BIA-BID/1606/2020 to R.G.M). The Algarve Biomedical Center Research Institute Microscopy Unit was partially supported by the Portuguese national funding (PPBI-POCI-01-0145-FEDER-022122). This work was developed with the support of the research infrastructure Congento (LISBOA-01-0145-FEDER-022170).

The funding bodies had no roles in study design, data collection, data analysis, data interpretation, or manuscript writing.

## Author contributions

Conceptualization: S.V., R.D.S., C.B., R.G.M., and T.A. Investigation: S.V., R.D.S., I.K., A.F., A.B., H.B., B.K., A.D., C.O., K.C., A.H.Y.T., S.A., L.J.J., and L.N. Data analysis: S.V., R.D.S., I.K., A.B., M.B., H.B., M.C., X.Z., M.U., A.J.W., R.G.M., and T.A. Supervision: S.V., R.D.S., B.J.A., J.M., C.L.M., K.G., C.B., R.G.M., and T.A. Funding acquisition: S.V., R.D.S., H.B., A.J.W., B.J.A., J.M., C.L.M., K.G., C.B., R.G.M., and T.A. Writing original draft: S.V., R.D.S., R.G.M., and T.A with help from all authors. All authors read and approved the final manuscript.

## Declaration of interests

J.M. is a shareholder and advisor of Century Therapeutics and Aelian Biotechnology.

## METHODS

### RESOURCE AVAILABILITY

#### Lead contact

Further information and requests for resources and reagents should be directed to and will be fulfilled by the lead contact, Thomas Arnesen

#### Material availability

Materials generated in this study are available upon reasonable request to the authors.

### EXPERIMENTAL MODEL AND SUBJECT DETAIL

#### Cell lines and culturing

HAP1 WT cells (clone C631; sex: male; RRID:CVCL Y019) and the HAP1 gene-KO cell lines *NAA30*-KO (HZGHC006637c010) and *NAA35*-KO (HZGHC006636c011) were obtained from Horizon Genomics GmbH (Vienna, Austria). HAP1 *NAA38*-KO (KO2 in original ref) and an unmodified HAP1 control cell line that underwent CRISPR treatment (herein referred to as WT2) were a generous gift from Dr. Scott Dixon, Stanford University (Cao et al., 2019). Gene disruption in the KO cell lines were confirmed by Sanger sequencing of PCR products of the edited genomic region. HAP1 cells were maintained in IMDM (Gibco) supplemented with 10% FBS and 1% penicillin-streptomycin (both from Sigma-Aldrich). HeLa cells were obtained from ATCC (clone CCL-2; sex: female; RRID:CVCL 0030) and maintained in high glucose DMEM supplemented with 10% FBS, 4 mM L-glutamine and 1% penicillin-streptomycin (all from Sigma-Aldrich). Cells were detached using Trypsin (Gibco) and maintained at 37°C in a humidified atmosphere with 5% CO2. Cells were routinely tested for mycoplasma using the MycoAlert Detection Kit (Lonza) or by NucBlue DAPI staining (Invitrogen). For CRISPR screening HAP1 cells with lowest possible passage number were used. For all other experiments, HAP1 cell lines were passaged and sorted until diploid status was confirmed by an Accuri C6 flow cytometer (BD) using propidium iodide staining before they were used in experiments (Beigl et al., 2020).

#### Fly work and crosses

Flies were raised under standard procedures. Source of all strains used in this study are listed in Key Resources Table. All males used in this study resulted from different *Drosophila* crosses. Control males resulted from crosses between y^1^ females with OR males or with FM0/DP (1;Y) y^+^ males. *Naa30A* deletion males resulted from crosses between y^1^, *Naa30A*D^74^/FM0 females or w, y^1^, *Naa30A*^Δ74^/FM0 females with OR males, or between y^1^, *Naa30A*^Δ74^/FM0 females with FM0/DP (1;Y) y^+^ males. *Naa30A* deletion + genomic rescue resulted from crosses between y^1^, *Naa30A*^Δ74^/FM0 females or w, y^1^, *Naa30A*^Δ74^/FM0 females with w^1118^;; gNaa30A-myc males. *Naa30A* deletion with Mhc-Gal4 males resulted from crosses between w, y^1^, *Naa30A*^Δ74^/FM0 females with w;; Mhc-Gal4 males. *Naa30A* deletion with UAS-UcbE2M males resulted from crosses between w, y^1^, *Naa30A*^Δ74^/FM0 females with w;; UAS-UcbE2M3xHA males. *Naa30A* deletion males ubiquitously expressing UcbE2M resulted from crosses between w, y^1^, *Naa30A^Δ74^*/FM0;; UAS-UbcE2M/TM3,sb females with w;; Da-Gal4 males. *Naa30A* deletion males expressing UcbE2M in muscles resulted from crosses between w, y^1^, *Naa30A^Δ74^*/FM0;; UAS-UbcE2M/TM3,sb females with w;; Mhc-Gal4 males.

#### Generation of mutants and transgenic flies

*Drosophila Naa30A* deletion was generated by imprecise excision of a P element, inserted in the 5’ UTR of the *Drosophila Naa30A* gene (CG11412). Briefly, the y1 P(EPgy2)Naa30AEY10202 w67c23 (BL16976) was crossed with PΔ2-399B, Sb/TM3, Ser transposase line. F1 male flies were balanced by crossing them with female virgins Df(1)pn38/FM0. F1 female flies, resulting from this second cross, were chosen as candidate deletions (white eye) and single-female crosses with males Df(1)pn38/FM0 were performed for stock balancing. Deletion sizes were determined by PCR and sequencing, using a forward primer 1 kb upstream (CAAGGAAAGTGGAGGAAGTGC) and a reverse primer 1.5 kb downstream (GGTATGTATCCCTCGCCAATG) of the 5’ end of CG11412. Nearly 100 lines were tested and the y1, *Naa30A^Δ74^*, w was selected. Removal of the w recessive marker to create the stock y1, *Naa30A^Δ74^*, was performed by recombination with the wild-type X chromosome from the Oregon-R (OR) strain.

For generation of *Drosophila* strains carrying a genomic fragment with the Naa30A gene, a fragment containing the genomic sequence of *Naa30A* franked by 1 kb pairs upstream and downstream of the 5’UTR and 3’UTR was synthesized by Genescript (Piscataway, NJ, USA). This fragment was sequenced and cloned into pCaSpeR2 in the PstI and EcoRI restriction sites to create the pCaSpeR2-gNaa30. Microinjection of pCaSpeR2-gNaa30 and selection of transfected strains was performed by BestGene (Chino Hills, CA, USA).

##### Saccharomyces cerevisiae strains

The *S. cerevisiae* haploid strain BY4741 Arl3-GFP was obtained from Invitrogen. Deletion of *NAA30/MAK3/YPR051W* was performed by homologous recombination as previously described and verified by PCR (Osberg et al., 2016). Arl3-GFP *naa30Δ* strain was transformed with pBEVY-U-HA-DmNAA30, and both the Arl3-GFP WT and *naa30Δ* strains were transformed with empty pBEVY-U (Miller et al., 1998) to enable the same growth conditions. Yeast strains were grown in SD-Ura medium (Sunrise Science Products) at 30°C. For microscopic imaging, the cells were diluted from overnight cultures and grown to exponential growth phase OD600 0.8-1.2. Cell preparation and live imaging were performed as previously described (Aksnes et al., 2013; Osberg et al., 2016). Images were taken using a Leica DMI6000 B widefield microscope equipped with a Leica DC500 camera and a 100 ×1.4 NA oil objective in addition to a 2 × magnification lens.

#### METHOD DETAILS

##### Pooled genome-wide CRISPR dropout screens

Pooled CRISPR-Cas9 dropout screens in HAP1 cell lines were performed essentially as described (Aregger et al., 2020; Hart et al., 2017). Briefly, 90 million cells were transduced with the TKOv3 lentiviral library (70,948 guides (4 guides/gene) targeting 18,053 protein-coding genes, lentiCRISPRv2 backbone) at an MOI of ~0.30 ensuring >200-fold library representation after selection. 24 h after infection, the transduced cells were selected for viral integration with medium containing 1 μg/mL puromycin (Sigma) for 48 h. The cells were collected, and the pooled cells were split into three replicates containing 15 million cells each, passaged every third day and maintained at 200-fold library coverage. 20-30 million cells were collected for genomic DNA extraction after selection (T0 reference) and at every passage until day 18 post-selection (T18) or ~15 population doublings (for *NAA30*-KO screen T14 and for *NAA38*-KO screen T15). Genomic DNA was extracted from the cell pellets using the Wizard Genomics DNA Purification Kit (Promega) and the concentration was determined using the Qubit dsDNA Broad Range Assay kit (Invitrogen).

Sequencing libraries were prepared from 52.5 μg gDNA using two-step PCR strategy with NEBNext Ultra II Q5 Polymerase (NEB). First the gRNA regions were enriched from the genome, then the gRNAs were amplified with primers harboring Illumina TruSeq adapters with i5 and i7 indices (Aregger et al., 2019; Hart et al., 2017). Barcoded libraries were gel-purified and subsequently sequenced on an Illumina HiSeq2500 (RRID:SCR_016383) using dual-index single-read sequencing with standard sequencing primers with HiSeq SBS Kit v4 reagents. The first 21 cycles of sequencing were dark cycles, or base additions without imaging. The actual 37-bp index read containing the gRNAs began after the dark cycles. Sequence data were demultiplexed and gRNA flanking sequences were trimmed. All primer sequences can be found in **Table S7**. Scoring of log2-fold change (LFC) and quantitative genetic interactions (qGI), reproducibility analysis of *NAA35*-KO screens and biological pathway enrichment analysis were performed as in (Aregger et al., 2020). The Pearson correlation coefficients of the qGI scores measured in two replicated screens was adjusted to the similarity of a *NAA35*-KO screen to a panel of HAP1-KO screens. The resulting Within vs Between replicate Correlation (WBC) score provides a confidence of reproducibility that can be interpreted as a z-score (Billmann et al., 2022). The HAP1 *NAA35*-KO screens were performed three separate times with three replicates each, whereas the *NAA30* and *NAA35* screens were performed once with three replicates each.

##### HAP1 cell harvesting for proteomics analysis

HAP1 WT, *NAA30*-KO, *NAA35*-KO, and *NAA38*-KO cells were grown in 10-cm dishes to 70-80% confluency. Cells were washed twice in ice-cold DPBS buffer (Gibco), detached by scraping in ice-cold DBPS with 1 × cOmplete protease inhibitor (Roche), and collected by centrifugation 16,000 × *g*, 15 s, 4°C (twice). Cell pellets were flash frozen in liquid nitrogen and stored at −80°C until further processing. For each cell line, four samples per proteome study were used. The N-terminal acetylation status was determined by positional proteomics using strong cation exchange (SCX) enrichment, while protein abundance was determined by label-free quantitative (LFQ) shotgun proteomics. Detailed information is given below.

##### N-terminal acetylation proteomics

As protein N-terminal acetylation (either *in vivo* or *in* vitro) neutralizes the N-terminal positive charge, the charge of N-terminal peptides is altered compared to that of internal peptides. This altered biophysical property can be used to enrich for N-terminal peptides by using low pH strong cation exchange (SCX) chromatography. SCX enrichment of N-terminal peptides from HAP1 control and NatC KO cells was performed essentially as previously described (Van Damme et al., 2011a). Cell pellets were thawed on ice, resuspended in 1.5 mL of ice-cold CHAPS lysis buffer (50 mM sodium phosphate pH 7.5, 100 mM NaCl, 0.8% (wt/vol) CHAPS in water, and 1 × cOmplete EDTA-free protease inhibitor cocktail (Roche)) and incubated on ice for 30 min. Samples were cleared by centrifugation at 16.000 × *g* for 15 min at 4°C. Protein concentrations were determined using Bradford reagent (Bio-Rad) and 1 mg of proteins was collected. To this, guanidinium chloride (GuHCl) was added to a final concentration of 2 M. Subsequently, proteins were reduced and alkylated for 15 min at 37°C in the dark and at a pH of 7.9 with iodoacetamide (30 mM f.c.) and TCEP-HCl (15 mM f.c.). Samples were subsequently desalted on Illustra NAP-10 columns (GE Healthcare, Cat # 17085402) and eluted in 50 mM sodium phosphate pH 8 containing 1.33 M GuHCl, after which the volume was reduced from 1.5 mL to 1 mL by vacuum drying (to reach a final concentration of 2 M GuHCl). To enable the assignment of *in vivo* Nt-acetylation events (Ac), all primary protein amines were blocked at the protein level making use of a N-hydroxysuccinimide ester of acetic acid encoded with stable heavy isotopes (i.e. an NHS ester of ^13^C1D3-acetate (Staes et al., 2011)). In this way, it is possible to distinguish between *in vivo* (Ac) and *in vitro* (AcDC) acetylated N-termini and to calculate the degree of *in vivo* Nt-acetylation as described below. Samples were acetylated with 10 mM (final concentration) of AcDC for 1 h at 30°C. The NHS ester was added once more, and incubation proceeded for an additional hour at 30°C. The excess of acetylation reagent was quenched with 40 mM glycine (2-fold molar excess) and incubated for 10 min at room temperature (RT) followed by the addition of 100 mM hydroxylamine and an incubation for an additional 10 min at RT. Samples were then desalted on a Illustra NAP-10 column (GE Healthcare) and eluted in 10 mM ammonium bicarbonate buffer (pH 7.6). Before trypsin digestion, samples were boiled for 5 min and cooled for 10 min on ice. Trypsin (Promega, Cat # V5111) was added in a trypsin/protein ration of 1/50 (w/w) and incubated ON at 37°C. Samples were then vacuum-dried.

Dried peptides were re-dissolved in 212 µL of pyro-glu buffer (16 mM NaCl, 0.5 mM EDTA, 3 mM cysteamine and 50 μM aprotinin (Roche)). Purified pGAPase was activated by adding 1 μL of 800 mM NaCl, 1 μL of 50 mM EDTA (pH 8.0) and 11 μL of 50 mM cysteamine, and was incubated for 10 min at 37°C. Activated pGAPase and Q-cyclase (both parts of the Tagzyme kit, Qiagen, Cat # 34342) were added to the peptides and incubated for 1 h at 37°C.

Samples were diluted to 1 mL with SCX buffer A and pH was adjusted to 2.98-3.02. Peptide concentrations were measured on Lunatic microfluidic device (Unchained Labs) and 700 μg of peptide material was loaded onto the dual mode SCX cartridge according to the following protocol: a) wet with acetonitrile (1 mL); b) wash with Milli-Q water (1 mL); c) equilibrate with SCX buffer A (2 mL); d) load sample (1 mL); e) wash with SCX buffer A (1 mL); f) elute with SCX buffer B (5 mL) and 1 mL of SCX buffer B containing 5 mM NaCl, and this 6 mL was collected; g) adjust the pH of the elution to about 6.0. Peptides were then dried and re-dissolved in 1 mL in loading buffer (0.1% TFA and 2% ACN). Methionines were oxidized by 0.06% H2O2 (final concentration) for 30 min at 30°C, followed by SampliQ SPE C18 desalting and clean-up (Agilent Technologies, Cat # 5982-1135). Peptides were eluted in elution buffer (60% acetonitrile (ACN), 0.1% TFA), vacuum-dried and stored at −20°C. SCX buffer A: mix buffer A1 (60 mg NaH2PO4 + 50 ml H2O) with buffer A2 (60 mg H3PO4 + 50 mL H2O) to a pH of 3.00. SCX buffer B: mix buffer B1 (57.6 mg NaH2PO4 + 15 mL H20 + 35 mL acetonitrile) with buffer B2 (57.6 mg H3PO4 + 15 mL H2O + 35 mL acetonitrile) to a pH of 3.00.

Purified peptides were re-dissolved in 22 μL loading solvent A (0.1% TFA in water/ACN (98:2, v/v)) and the peptide concentration was measured on the Lunatic microfluidic device. 2 μg of peptides were injected for LC-MS/MS analysis on an Ultimate 3000 RSLCnano system in-line connected to an Orbitrap Fusion Lumos mass spectrometer (Thermo). Trapping was performed at 10 μL/min for 4 min in loading solvent A on a 20 mm trapping column (made in-house, 100 μm internal diameter (I.D.), 5 μm beads, ReproSil-Pur Basic-C18-HD (Dr. Maisch, Germany). The peptides were separated on a 200 cm µPAC column (C18-endcapped functionality, 300 μm wide channels, 5 μm porous-shell pillars, inter pillar distance of 2.5 μm and a depth of 20 μm; PharmaFluidics, Belgium). The column temperature was kept constant at 50°C. Peptides were eluted by a linear gradient reaching 55% MS solvent B (0.1% formic acid (FA) in water/acetonitrile (2:8, v/v)) after 115 min, 99% MS solvent B at 120 min, followed by a 10-min wash at 99% MS solvent B and re-equilibration with MS solvent A (0.1% FA in water). The first 15 min the flow rate was set to 750 nL/min after which it was kept constant at 300 nL/min. The mass spectrometer was operated in data-dependent top speed mode with a cycle time of 3 s, automatically switching between MS and MS/MS acquisition. Full-scan MS spectra (300-1500 m/z) were acquired at a resolution of 120,000 in the Orbitrap analyzer after accumulation to an AGC value of 200,000 with a maximum injection time of 250 ms. The most intense ions above a threshold value of 5,000 and a charge state ranging from 2 to 7, subjected to a dynamic exclusion of 60 s, were isolated in the quadrupole for fragmentation in the ion routing multipole at a normalized collision energy of 34% after accumulation of precursor ions at a target value of 12,000 with a maximum of 40 ms with an isolation width of 1.2 Th. The fragments were analyzed in the Ion Trap Analyzer at normal scan rate.

Mascot generic files were created using the Mascot Distiller software (version 2.7.1.0, Matrix Science). Peak lists were searched with the Mascot search engine using the Mascot Daemon interface (version 2.6.2, Matrix Science) against the SwissProt database restricted to human proteins (*Homo sapiens* database release version of January 2020). Spectra were searched twice, one search to quantify the N-terminal acetylated peptides and another search to identify all the peptides including the ones that are not N-terminally acetylated. Both peptide searches were set with semi-ArgC/P enzyme (/P indicates arginine can also be cleaved when followed by a proline), allowing 1 missed cleavage; heavy acetylation (^13^C1D3) of lysine residues, carbamidomethylation of cysteine residues and oxidation of methionine were set as fixed modifications. Mass tolerance of the precursor ions was set to 10 ppm (with Mascot’s C13 option set to 1) and of fragment ions to 0.5 Da. The instrument was set to ESI-TRAP. In the first search, to allow identification and quantification of N-terminal acetylated peptides, a quantitation method with two different components was made, defining light and heavy acetylation of peptide N-termini as respectively light and heavy exclusive modification groups. Only peptides that were ranked first and scored above the threshold score set at 99% confidence were withheld. In a second search, peptides without N-terminal acetylation were identified using a similar search parameter set, including the cyclization of N-terminal glutamine residues to pyroglutamate (pyro-Glu) residues as a variable modification. Identified peptides were quantified using Mascot Distiller Toolbox (version 2.7.1.0, Matrix Science) in the precursor mode. For processing of all MS data, the ms_lims software platform was used (Helsens et al., 2010). Peptides shorter than 8 amino acid residues were removed for downstream analysis. To further check the N-terminal data, all quantified acetylated peptides were extracted from ms_lims. After extraction, peptides were filtered on confident spectrum (“TRUE”), tryptic cleavage and start position (one or two) to retrieve the true N-terminal peptides. The degree of *in vivo* acetylation was calculated separately using the light/heavy (Ac/AcDC) ratio reported by the Mascot Distiller Toolbox and by the equation Ac% = ratio (Ac/AcDC) / (ratio (Ac/AcDC) + 1). In those cases where N-terminal peptides were identified and quantified multiple times, the median and standard deviation of the acetylation degrees of the individual peptides was calculated. Processed data are presented in **Table S3**.

##### Label-free quantitative shotgun proteomics

Cell pellets were dissolved in 0.5 mL urea lysis buffer (8 M urea pH 8.0, 20 mM HEPES). The cells were then sonicated 3 × 15 s (with 1 min incubations on ice) and centrifuged at 16,000 × *g* for 15 min at RT. The protein concentrations of the samples were determined by the Bradford assay (Bio-Rad). Proteome samples were prepared in quadruplicate for each condition. From each replicate, equal protein amounts (1 mg) were used for further analysis. Proteins were reduced with 5 mM DTT f.c. for 30 min at 55°C and alkylated with 10 mM f.c. iodoacetamide at RT in the dark for 15 min. The samples were diluted with 20 mM HEPES at pH 8.0 to achieve a urea concentration of 4 M and then the proteins were digested with 1/100 (w/w) endoLysC (Lysyl Endopeptidase, Wako, Cat # 121-05063) for 1.5 h at 37°C. All samples were further diluted with 20 mM HEPES at pH 8.0 to a final urea concentration of 2 M, and the proteins were digested with 1/100 (w/w) trypsin (Promega, Cat # V5111) overnight at 37°C. Digested samples were acidified with TFA (to a final concentration of 1% TFA), which allows precipitation of undigested proteins. Samples were incubated on ice for 15 min, centrifuged at 16,000 × *g* for 15 min at RT, and the supernatant was transferred to a new tube. The peptides (half of the sample was used, the other half was stored) were then purified on a SampliQ SPE C18 cartridge (Agilent Technologies, Cat # 5982-1111.). The eluted peptides were dried and stored at −20°C before MS analysis.

MS/MS analysis was performed similar as described for the enrichment of N-terminal peptides (acetylation proteomics) with some minor changes outlined below. Peptides were separated on an Ultimate 3000 RSLCnano system in-line connected to an Orbitrap Fusion Lumos mass spectrometer (Thermo Scientific) with a linear gradient reaching 55% MS solvent B after 145 min and 99% MS solvent B at 150 min, followed by a 10-min wash at 99% MS solvent B and re-equilibration with MS solvent A. All other settings were kept as described above.

The generated MS/MS spectra were processed with MaxQuant (version 2.0.1.0) (Cox and Mann, 2008) using the Andromeda search engine with default search settings, including a false discovery rate set at 1% on both the peptide and protein level. Spectra were searched against the sequences of the human proteins in the Swiss-Prot database (release May 2021). The enzyme specificity was set at trypsin/P, allowing for two missed cleavages. Variable modifications were set to oxidation of methionine residues and N-terminal protein acetylation. Carbamidomethylation of cysteine residues was put as fixed modification. In the settings of advanced identifications, matching between runs was enabled (with standard settings). All other settings were kept as standard. Proteins were quantified by the MaxLFQ algorithm integrated in the MaxQuant software. Minimum of two ratio counts and both unique and razor peptides were considered for protein quantification. The iBAQ function was also used. Further data analysis was performed with the Perseus software (version 1.6.15.0) (Tyanova et al., 2016) after loading the protein groups file from MaxQuant. Proteins only identified by site, reverse database hits and potential contaminants were removed, and replicate samples were grouped.

Proteins with less than three valid values in at least one group were removed, and missing values were imputed from a normal distribution around the detection limit. Then, a multiple sample t-tests was performed (FDR = 0.01 and S0 = 0) after which the z-scores of the significant proteins of each sample were calculated and further analyzed by hierarchical clustering (Euclidean with standard Perseus settings). Pairwise comparisons were also performed by a t-test (FDR = 0.01 and S0 = 0.1) between the different KO cell lines and control samples. Processed data is presented in **Table S4**.

##### Plasmids

The plasmid pCMV6-UBE2M-Myc-DDK (DDK is the same as FLAG®) was obtained from OriGene Technologies (Rockville, Maryland, USA, # RC208946). pcDNA3.1-NAA30-V5 was previously described and was constructed using cDNA from a human cell line (Starheim et al., 2009). For both plasmids, point mutations were introduced using the Q5 Site-Directed Mutagenesis Kit (New England Biolabs) according to the manufacturer’s instructions. Mutations were confirmed by Sanger sequencing. The bicistronic pUC-UBE2M-V5-P2A-GST-GFP plasmid was custom-made by VectorBuilder (Chicago, Illinois, USA) and expresses the indicated UBE2M P2A construct from a *CMV* promoter. pcDNA6.2-hUBR4-V5-Lumio plasmid was a kind gift from Dr. Yong Tae Kwon, Seoul National University College of Medicine, and contains a 15.9-kb human *UBR4* open reading frame (Tasaki et al., 2009). The plasmids pCDH-EF1-UBR4N-FLAG-T2A-copGFP and pCDH-EF1-UBR4C-FLAG-T2A-copGFP were a kind gift from Dr. Fabio Demontis and Dr. Liam C. Hunt, St. Jude Children’s Research Hospital (Hunt et al., 2019). These plasmid expresses FLAG-tagged truncated variations of mouse UBR4, where UBR4-N corresponds to a N-terminal region which includes the UBR box (p.1222-1996) while UBR4-C corresponds to the C-terminal UBR4 domain (p.4364-5180). *Drosophila Naa30A* was amplified with the introduction of a N-terminal HA-tag from the cDNA clone LD45352 (DGRC Stock 3896; RRID:DGRC_3896) and then subcloned into pBEVY-U (Miller et al., 1998) downstream of the *ADH1* promoter using the XmaI and EcoRI sites, yielding plasmid pBEVY-U-HA-DmNAA30. pBEVY-U-HA-hNAA30 was previously described and expresses human NAA30 from the *ADH1* promoter (Osberg et al., 2016). All primers can be found in **Table S7**.

##### Immunoblotting

Cells were washed with cold PBS (Gibco), scraped on ice, and collected at 16,000 x *g* for 15 s at 4°C. Cell pellets were lysed in IPH lysis buffer (50 mM Tris-HCl pH 8.0, 150 mM NaCl, 5 mM EDTA, 0.5% NP-40) supplemented with 1 × cOmplete EDTA-free protease inhibitor cocktail (Roche) for 30 min on ice. Cell lysates were cleared by centrifugation at 16,000 × *g* for 5 min at 4°C and protein concentration was determined using the Pierce BCA protein assay kit (Thermo Scientific). Cell lysates were mixed with Laemmli sample buffer (Alfa Aesar) and denatured at 95°C for 5 min. Next, 20-30 μg of total protein was resolved on 8-16% TGX stain-free gels and transferred onto nitrocellulose membrane using the Trans-Blot Turbo RTA Transfer kit and the Trans-Blot Turbo Transfer system set to 7 min transfer protocol (all from Bio-Rad). Stain-free gel and blots were imaged using the Gel Doc EZ imaging system (Bio-Rad) when applicable. For the UBR4 blots, proteins were typically resolved on 7.5% TGX stain-free gels and transferred onto nitrocellulose membrane (GE Healthcare, Amersham Protran) using wet-transfer at 100 V for 60 min. The blots were blocked for 1 h at RT in 5% non-fat dry-milk (AppliChem) resuspended in 1 × TBS-T (20 mM Tris pH 7.5, 150 mM NaCl, 0.05% (v/v) Tween-20). Subsequently, the blots were incubated with primary antibody diluted in 1% blocking buffer overnight at 4°C followed by washing and 2 h incubation with HRP-conjugated secondary antibody diluted in 3% blocking buffer. Protein bands were visualized using SuperSignal West Pico chemiluminescent substrate (Thermo Scientific) and the chemiluminescent signals were captured with a ChemiDoc XRS+ imaging system coupled with Image Lab Software version 6.0.1 (both from Bio-Rad). Densitrometry analysis were performed in Image Lab.

The following antibodies were used: anti-IST1 (GeneTex, GTX101972, 1:1000), anti-NAA30 (Sigma-Aldrich, HPA057824, 1:1000), anti-UBE2M (Abcam, ab109507, 1:1000), anti-HK1 (Thermo Scientific, MA5-14789, 1:2000), anti-UBE2F (Abcam, ab185234, 1:1000), anti-DIS3 (Thermo Scientific, PA5-78427, 1:1000), anti-ARFRP1 (Sigma-Aldrich, HPA04702, 1:1000), anti-CAPNS1 (Thermo Scientific, PA5-82266, 1:1000), anti-SLC10A7 (Sigma-Aldrich, SAB2102163, 1:1000), anti-RSPRY1 (Thermo Scientific, PA5-32048, 1:1000), anti-FLAG (Sigma, clone M2, F3165, 1:3000), anti-V5 (Invitrogen, R960CUS, 1:5000-20,000), anti-GFP (Roche, clones 7.1 and 13, 11814460001, 1:5000), anti-UBR1 (Bethyl Laboratories, A302-988A-M, 1:1000), anti-UBR2 (Abcam, ab217069, 1:2000), anti-UBR4 (Abcam, ab86738, 1:1000), anti-UBR5 (Cell Signaling, 65344, 1:1000), anti-KCMF1 (Sigma-Aldrich, HPA030383, 1:1000), anti-UBE2A/UBE2B (Abcam, ab31917, 1:2000), anti-RGS10 (Abcam, ab154172, 1:1000), anti-CUL1 (Cell Signaling, 4995, 1:1000), anti-CUL2 (Abcam, ab166917, 1:1000), anti-CUL3 (Cell Signaling, 2759, 1:1000), anti-CUL4A (Cell Signaling, 2699, 1:1000), anti-CUL4B (Proteintech, 12916-1-AP, 1:1000), anti-CUL5 (Abcam, ab264284, 1:1000), anti-RBX1 (Abcam, ab221548, 1:1000), anti-RBX2/RNF7 (Abcam, ab181986, 1:1000), anti-BCL2 (Proteintech, 12789-1-AP, 1:500), anti-EEA1 (Santa Cruz, sc-33585, 1:1000), anti-p62/SQSTM1 (Santa Cruz, sc-28359, 1:300-1:1000), anti-LC3B (Thermo Scientific, PA1-46286, 1:1000), anti-vinculin (Abcam, ab129009, 1:10000), anti-GAPDH (Santa Cruz, sc-47724, 1:10000), HRP-linked-sheep-anti-Mouse (Cytiva, NA931, 1:3000-1:20000), and HRP-linked-donkey-anti-Rabbit (Cytiva, NA934, 1:1000-1:10000).

##### N-terminal variants of UBE2M-FLAG

To investigate the stability of N-terminal variants of UBE2M, HAP1 *NAA30*-KO cells were transiently transfected with a set of Met-X-UBE2M-FLAG constructs where the second residue was glycine, alanine, (NatA), aspartic acid, glutamic acid, (NatB), isoleucine (native), leucine, tyrosine, phenylalanine (NatC/E/F substrate), or proline (stabilizing control). HAP1 *NAA30*-KO cells were seeded at a density of 2.5 × 10^6^ cells per 10-cm dish and incubated overnight. The medium was replaced, and the cells were transfected for 24 h with a mixture of 6 μg plasmid DNA, 24 μL X-tremeGENE 9 DNA transfection reagent (Roche) and 500 μL Opti-MEM media (Gibco), according to the manufacturer’s instructions. Protein levels were determined by immunoblotting using anti-FLAG clone M2 (Sigma, F3165, 1:3000).

##### UBE2M reporter assay

To exclude any effects at the transcriptional or protein synthesis levels, we transfected HAP1 WT and *NAA30*-KO cells with a bicistronic P2A reporter vector allowing expression of both UBE2M-V5 and GST-GFP from a *CMV* promoter. HAP1 cells were seeded the day before and incubated overnight. HAP1 WT and *NAA30*-KO cells were transfected with 1.25 μg and 6.0 μg pUC-UBE2M-V5-P2A-GST-GFP, respectively. WT cells were co-transfected with an empty plasmid of similar size to obtain equal amounts of DNA in the transfection mixture. Transfections were performed using X-tremeGENE 9 DNA Transfection Reagent (Roche) (3:1 ratio of reagent to DNA) according to the manufacturer’s instructions. After 24 h, transfected cells were harvested and processed for immunoblotting. The transfections were performed four independent times. The immunoblot were probed three consecutive times with anti-V5 (1:5000), anti-GFP (1:5000), and finally anti-GAPDH (1:10000). Densitometry analysis was performed using Image Lab 6.0.1. Background-adjusted signal intensities of UBE2M-V5 were normalized to GST-GFP and expressed relative to WT. Data are shown as mean ± SD of four biologically independent experiments. Significance was determined using two-tailed unpaired t test.

##### Immunoprecipitation

HeLa cells were seeded at a density of 1.5 × 10^6^ cell per 10-cm dish and incubated overnight. The medium was replaced, and the cells were transfected with a mixture of 4 μg pcDNA3.1-NAA30-WT-V5, pcDNA3.1-NAA30-E321A-V5 or pcDNA3.1-LacZ-V5 (negative control), 12 μL X-tremeGENE 9 DNA Transfection Reagent (Roche) and 500 μL Opti-MEM (Gibco) according to the manufacturer’s instructions. After 24 h, the transfected cells were washed twice and scraped in cold PBS (Gibco) and collected by centrifugation at 1,200 × *g* for 5 min at 4°C. Cell pellets were lysed in 20 μl/mg IPH lysis buffer (50 mM Tris-HCl pH 8.0, 150 mM NaCl, 5 mM EDTA, 0.5% NP-40) with 1 × cOmplete EDTA-free protease inhibitor cocktail (Roche) on ice for 30 min. Cell debris was removed by centrifugation at 16,000 × *g* for 5 min at 4°C and the supernatants were transferred to low-protein binding tube. ~40 μL cell lysate was saved for immunoblot analysis. For immunoprecipitation (IP), the cell lysates were incubated with 2 μg anti-V5 antibody per dish (Invitrogen, Cat # R960CUS) for 3 h at 4°C on a rotating wheel. Then, 20 μL of pre-washed Dynabeads Protein G per dish (Invitrogen) was added to each IP sample, which were incubated overnight at 4°C to retrieve the immunocomplexes. The beads were washed twice in IPH lysis buffer, once in acetylation buffer (50 mM HEPES pH 8.5, 100 mM NaCl, 1 mM EDTA) and resuspended in 40 μL acetylation buffer per dish. The IP samples were used in N-terminal acetylation assays (see below) and immunoblotting. One transfected 10-cm dish were typically used per peptide in the acetylation assay. The immunoblots were probed with anti-V5 (Invitrogen, #R960CUS, 1:20,000 dilution).

##### *In vitro* N-terminal acetylation assay

The acetyltransferase activity of NAA30-WT-V5 and NAA30-E321A-V5 were determined using an *in vitro* N-terminal acetylation assay based on radiolabeled acetyl-CoA as previously described with minor modifications (Drazic and Arnesen, 2017). Briefly, 10 μL immunoprecipitated enzyme were mixed with 300 μM synthetic peptide and 50 μM [^14^C]-acetyl-CoA (PerkinElmer) in acetylation buffer (50 mM HEPES pH 8.5, 100 mM NaCl, 1 mM EDTA) to a final volume of 25 μL. The reactions were incubated for 1 h at 37°C using a Thermoshaker at 1400 rpm. After incubation, the beads were isolated using a magnetic rack and the reactions was quenched by spotting 20 μL of the reaction mixture onto P81 phosphocellulose paper (Millipore, 10 mm × 10 mm). The filters were washed 3 × 5 min with 10 mM HEPES (pH 7.4), air dried on paper, and immersed in 5 mL Ultima Gold F scintillation mixture (PerkinElmer). Radioactivity was detected by a Tri-Carb 2900TR liquid scintillation analyzer (PerkinElmer). The measured counts per minute (CPM) signal for each reaction, representing the product formation, was normalized to the amount of immunoprecipitated NAA30-V5 quantified from immunoblots. The experiment was performed three independent times with three technical replicates each. Reaction mixtures with immunoprecipitated V5-control plasmid were used as negative controls to assess background signal. Data are shown as mean ± SD from one representative experiment.

The peptides were custom-made by Innovagen AB (Lund, Sweden) or BioGenes (Berlin, Germany) to a purity of ≥95% and were dissolved in Ultrapure distilled water (Invitrogen). The 24-mer peptides contain seven unique amino acids at their N-terminus followed by the same 17 C-terminal residues (RWGRPVGRRRRPVRVYP). The C-terminal portion is derived from adrenocorticotropic hormone (ACTH) peptide sequence, but the lysines have been replaced with arginines to prevent possible lysine acetylation from interfering with the activity measurements. The following synthetic oligopeptides were used: UBE2M UniProtKB: P61081 ([NH2] <u>MIKLFSL</u> RWGRPVGRRRRPVRVYP [COOH]), ARFRP1 UniProtKB: Q13795 ([NH2] <u>MYTLLSG</u> RWGRPVGRRRRPVRVYP [COOH]), UNC50 UniProtKB: Q53HI1 ([NH2] <u>MLPSTSV</u> RWGRPVGRRRRPVRVYP [COOH]) and β-actin UniProtKB: P60709 ([NH2] <u>DDDIAAL</u> RWGRPVGRRRRPVRVYP [COOH]).

##### NAA30 rescue assay

HAP1 cells were seeded out at a density of 2.5 × 10^6^ cells per 10-cm dish and incubated overnight. HAP1 WT cells were transfected with 5 μg empty V5-vector, and HAP1 *NAA30*-KO cells were transfected with 5 μg of empty V5-vector, 4 μg pcDNA3.1-NAA30-WT-V5 + 1 μg of empty V5-vector, or 5 μg pcDNA3.1-NAA30-E321A-V5. Transfections were performed using X-tremeGENE 9 DNA Transfection Reagent (Roche) (3:1 ratio of reagent to DNA) according to the manufacturer’s instructions. After 24 h, transfected cells were harvested and processed for immunoblotting.

##### Inhibition of proteasomal and lysosomal protein degradation

HAP1 WT and *NAA30*-KO cells were seeded at a density of 2.5 × 10^6^ cells per 10-cm dish. After 24 h, the cells were treated with inhibitors of proteasomal (MG132 and bortezomib) and lysosomal (bafilomycin A, leupeptin and ammonium chloride) protein degradation for 6 h at 37°C. Cells were washed twice and scraped in ice-cold PBS (Gibco), and the resulting cell pellets were stored at −80°C until immunoblot processing. The final drugs concentrations were as follows: 10 μM MG132, 10 nM bortezomib (both from Calbiochem), 200 nM bafilomycin A, (MedChem Express), 100 μg/mL leupeptin, and 10 mM ammonium chloride (both from Sigma-Aldrich). DMSO (Sigma-Aldrich) was used as vehicle control.

##### siRNA transfection

HAP1 WT and *NAA30*-KO cells were seeded at a density of 200,000 cells per 6-cm dish for proteomic analysis, 3-500,000 cells per 10-cm dish for immunoblotting, and 60,000 cells/well in a 6-well plate for flow analysis. After 24 h the medium was replaced and the cells were transfected with 20 nM of ON-TARGETplus siRNA SMART pools using DharmaFECT1 (all from Horizon Discovery) according to the manufacturer’s instructions. A non-targeting siRNA pool (D-001810-10-05) was used as control treatment. The cells were harvested 72 h post-transfection. For lysosomal staining, the transfected cells were incubated with fresh medium containing 1 × LysoView 488 for 1 h at 37°C and then processed for flow analysis.

##### XY-UBE2M peptide pulldown assay

HeLa cells were transfected with pcDNA6.2-hUBR4-V5-Lumio (7.5 × 10^5^ cells/10-cm, 4 μg plasmid DNA, 48 h) and HAP1 WT cells were transfected with pCDH-EF1-UBR4N-FLAG-T2A-copGFP or pCDH-EF1-UBR4C-FLAG-T2A-copGFP (2.5 × 10^6^ cell/10-cm, 6 μg plasmid DNA, 24 h) using X-tremeGENE 9 DNA Transfection Reagent (Roche) (3:1 ratio of reagent to DNA) according to the manufacturer’s instructions. The cells were washed and scraped in cold PBS, collected by centrifugation at 1000 × *g* for 5 min, and stored at −80°C until further processing.

For the peptide binding assays we used a set of biotinylated 11-mer peptides derived from the N-terminal sequence of UBE2M (<u>MI</u>KLFSLKQQK(K-biotin)) bearing NatA type (AI-), NatB type (ME-), NatC/E/F type (MI/ML/MY/MF-), Arg/N-degron type 1 (RI-), or Arg/N-degron type 2 (FI-) N-terminal residues. Alternatively, we used a set of 11-mer Sindbid nsP4-derived peptides (<u>X</u>-IFSTIEGRTY(K-biotin)) where the N-terminal X residues was Arg, Phe or Gly (Cha-Molstad et al., 2015). The peptides were cross-linked to streptavidin-conjugated magnetic beads (Thermo Scientific) using 20 μg peptide to 20 μl beads diluted in ~500 μL 1 × TBST 0.01% (25 mM Tris pH 7.5, 150 mM NaCl, and 0.01% v/v Tween 20). The peptide-bead mixtures were incubated on a rotating wheel (20 rpm) for 2-3 h at 4°C, and subsequently washed twice in TBST 0.01%. and once in binding buffer (20 mM, HEPES pH 7.9, 200 mM KCl, 0.05% (v/v) Tween 20, 10% glycerol).

The cell pellets were resuspended in cold hypotonic buffer (10 mM HEPES pH 7.9, 10 mM KCl, and 1.5 mM MgCl2) supplemented with cOmplete protease inhibitor (Roche) and incubated on ice for 30 min. The cell suspensions were then subjected to five freeze-thaw cycles in liquid nitrogen followed by centrifugation at 16,000 × *g* for 5 min at 4°C. Protein concentration was determined using the Pierce BCA protein assay kit (Thermo Scientific). The cell lysates were diluted 1:10 in binding buffer (20 mM, HEPES pH 7.9, 200 mM KCl, 0.05% (v/v) Tween 20, 10% glycerol) with protease inhibitor. Next, 300-500 μg of total protein was transferred to the peptide-conjugated beads and the mixtures were gently rotated overnight at 4°C. The beads were washed three times with binding buffer, resuspended in 50 μL 1 × SDS sample buffer and incubated at 95°C for 5 min, followed by SDS-PAGE and immunoblot analysis. The peptides were custom-made by Innovagen AB, Lund, Sweden to a purity of >95%, and was dissolved in UltraPure distilled water (Invitrogen). All peptides can be found in **Table S7**.

##### Quantification of siUBR4 samples with MS

HAP1 WT and *NAA30*-KO cells were transfected with 20 nM of ON-TARGETplus UBR4 siRNA SMART pool (Cat # L-014021-01-0005) or non-targeting control pool (Cat # D-001810-10-05) using DharmaFECT1 (all from Horizon Discovery) and harvested 72 h post-infection. The cells were washed twice in ice-cold DPBS buffer (Gibco), detached by scraping in cold DBPS with 1 × cOmplete protease inhibitor (Roche) and collected by centrifugation 16,000 × *g* for 15 s at 4°C. The centrifugation step was repeated once to remove as much DPBS as possible. The resulting cell pellets were flash frozen in liquid nitrogen and stored at −80°C until further processing. There were a total four samples with four replicates each (16 tubes in total).

Samples were prepared using the S-Trap protocol according to the manufacturer’s instructions (ProtiFi, Cat # C02-mini-40). Cells were lysed in 100 μL S-trap lysis buffer (5% SDS in 50 mM TEAB, pH 8.5) on ice. The cells were then sonicated 3 ×15 s (with 1 min incubations on ice) each and centrifuged at 16 000 × *g* for 10 min at 4°C. The supernatant was collected, and the protein concentration was determined using the Pierce BCA protein assay kit (Thermo Scientific). From each sample, 100 μg of protein material was retrieved and the volume was adjusted with S-trap lysis buffer to 46 µL. Subsequently, proteins were reduced by addition of DTT (15 mM f.c.) and incubation for 15 min at 55°C, followed by alkylation with iodoacetamide (30 mM f.c.) for 15 min in the dark. Next, 5 μL of 12% phosphoric acid (to a f.c. of 1.1%) was added. The pH of the samples was checked if this was <1. 350 µL of protein binding/wash buffer (90% methanol, 100 mM TEAB, pH 7.55) was added to the samples before loading them onto the S-Trap column. Columns were centrifuged for 30 s at 4,000 × *g* at RT and the flow-through was discarded. The column was washed three times by addition of 400 µL of binding/wash buffer and centrifuged for 30 s at 4,000 × g. The samples were centrifuged a final time for 1 min at 4,000 × *g* to completely remove the remaining binding/wash buffer. To each column, 125 μL digestion buffer (trypsin in 50 mM TEAB in a 1/100 (w/w)) was added and incubated ON at 37°C. Peptides were eluted in three steps: first, 80 μL elution buffer 1 (50 mM TEAB in ddH2O) was added and samples were centrifuged for 1 min at 4,000 × *g*. Followed by an addition of 80 μL of elution buffer 2 (0.1% FA in ddH2O) and a second centrifugation step. Finally, 80 μL of elution buffer 3 (50% ACN, 0.1% FA in ddH2O) was added and samples were centrifuged a third time. Elution fractions were pooled and vacuum-dried.

Samples were re-dissolved in 100 µL 100 mM TEAB with 10% ACN pH 8.5 and the peptide concentration was measured on Lunatic microfluidic device (Unchained Labs). For each sample, 50 µg of peptide material was retrieved, and the volume was adjusted to 100 µL. TMTpro 16-plex labels (Thermo Fisher Scientific, Cat # A44521) were thawed at RT, suspended in 20 µL ACN and vortexed vigorously. Each sample was labelled by adding 10 µL label and incubating for 1 h at RT (600 rpm) (see **Table S5** for an overview of sample labelling).

The reaction was quenched with 1:20 v:v 5% hydroxylamine (NH2OH) (f.c. 0.25%) for 15 min at RT (600 rpm) and the 16 fractions were ultimately combined. 100 µg of peptide material was retrieved and vacuum dried. The labeled peptides were re-dissolved in 100 µL 0.1% TFA and 1 µL 100% TFA was added to lower the pH before sample clean-up. Samples were desalted using OMIX C18 tips (Agilent Technologies, Cat # A57003100) according to the manufacturer’s instructions and vacuum dried. Peptides were re-dissolved in 100 µL solvent A (0.1% TFA in water/ACN (98:2, v/v)) and injected for fractionation by RP-HPLC (Agilent series 1200) connected to a Probot fractionator (LC Packings). Peptides were first loaded in solvent A on a 4 cm pre-column (made in-house, 250 µm internal diameter (ID), 5 µm C18 beads, Dr. Maisch) for 10 min at 25 µL/min and then separated on a 15 cm analytical column (made in-house, 250 µm ID, 3 µm C18 beads, Dr Maisch). Elution was done using a linear gradient from 100% RP-HPLC solvent A (10 mM ammonium acetate (pH 5.5) in water/ACN (98:2, v/v)) to 100% RP-HPLC solvent B (70% ACN, 10 mM ammonium acetate (pH 5.5)) in 100 min at a constant flow rate of 3 µL/min. Fractions were collected every min between 20 and 96 min and pooled every 12 min to generate a total of 12 samples for LC-MS/MS analysis. All 12 fractions were dried under vacuum in HPLC inserts and stored at −20°C until further use.

MS/MS analysis was performed similar as described for the enrichment of N-terminal peptides (acetylation proteomics) with some minor changes outlined below. Peptides were separated on an Ultimate 3000 RSLCnano system in-line connected to an Orbitrap Fusion Lumos mass spectrometer (Thermo Scientific) with a linear gradient reaching 55% MS solvent B after 85 min, 99% MS solvent B at 90 min, followed by a 10-min wash at 99% MS solvent B and re-equilibration with MS solvent A. Full-scan MS spectra (300-1500 m/z) were acquired at a resolution of 120,000 in the Orbitrap analyzer after accumulation to an AGC value of 400,000 with a maximum injection time of 50 ms. The most intense ions above a threshold value of 50,000 and a charge state ranging from 2 to 7, subjected to a dynamic exclusion of 60 s, were isolated in the quadrupole for fragmentation in the ion routing multipole at a normalized collision energy of 37% after accumulation of precursor ions at a target value of 100,000 with a maximum of 86 ms with an isolation width of 0.9 Th. The fragments were analyzed in the in the Orbitrap with a resolution of 50,000.

The generated MS/MS spectra were processed with MaxQuant (version 1.6.17.0) using the Andromeda search engine with default search settings, including a false discovery rate set at 1% on both the peptide and protein level. Spectra were searched against the sequences of the human proteins in the Swiss-Prot database (release January 2021). The enzyme specificity was set at trypsin/P, allowing for two missed cleavages. Variable modifications were set to oxidation of methionine residues and N-terminal protein acetylation. Carbamidomethylation of cysteine residues was put as fixed modification. To cope with the spectra that suffer from co-fragmentation, the precursor ion fraction (PIF) option was set to 75%. MS2-based quantification using TMTpro16plex labels was chosen as quantification method and a minimum ratio count of 2 peptides (both unique and razor) was required for quantification. Further data analysis was performed with the Perseus software (version 1.6.15.0) after loading the protein groups file from MaxQuant (load reporter ion intensities corrected per label). With Perseus, reverse proteins, proteins that are only identified by site and contaminants were removed. Samples were normalized by subtraction of the median per sample. Proteins with less than three valid values in at least one group were removed and missing values were imputed from a normal distribution around the detection limit. Then, a multiple sample t-test (false discovery rate (FDR) = 0.01) was performed to detect enrichments in the different samples. Also, t-tests (FDR = 0.01 and S0 = 0.1) were performed between the different cell types. Processed data is presented in **Table S5**.

##### WST-1 assay

Cell viability was determined using a colorimetric assay, based on the cleavage of WST-1 (Roche) to formazan dye by cellular mitochondrial dehydrogenases in viable cells. HAP1 cells were seeded in 96-well plates with a density of 5,000 cells/well. After 24 h the cells were incubated with fresh medium supplemented with 10% (v/v) WST-1 reagent for 3 h at 37°C. The absorbance of formazan dye was measured at 450 nm using a Tecan microplate reader. Samples with culture medium plus WST-1 and without cells were used as background control and was subtracted from the sample absorbance. Background subtracted values were expressed relative to WT. Data are shown as mean ± SD of three independent experiments with four technical replicates each. Significance was determined using one-way ANOVA with Dunnett’s correction.

##### Flow cytometry analysis

Cell granularity, size, and lysosomal content was determined by flow cytometry analysis. HAP1 cells were seeded in 6-well plates with a density of 150,000 cells/well or 60,000 cells/well for siRNA treatment and incubated for 24 h or 96 h, respectively. For lysosomal analysis, cells were stained with LysoView 488 (Biotium) for 1 h at 37°C according to the manufacturer’s protocol. Cells were detached using TrypLE Express (Gibco), washed with PBS, and resuspended in FACS buffer (PBS with 5% FBS). Cells were quantified on a LSRFortessa flow cytometer equipped with FACSDiva software version 9.0.1 (both from BD Life Sciences). Samples were run with a fluidics speed set to slow, giving an event rate of 200-400 events/sec. About 50,000 events were collected after gating for live cells and singlets using side scatter (SSC) vs forward scatter (FSC) and FSC-A vs FSC-H dot plots, respectively. Lysosomal content was assessed by detecting fluorescence at 530/30 nm (median FITC). Granularity and cell size was determined by the light scatter properties of the cells, using median SSC-A and median FSC-A. The flow cytometry results were analyzed using FlowJo version 10.8.1 (BD Life Sciences). Data are shown as mean ± SD of three independent experiments. Significance was determined using one-way ANOVA with Dunnett’s correction (NatC KO) or Šidák correction (siRNA).

##### Holographic live cell imaging

HAP1 cells were seeded in a 24-well plate (16,000 cells/well) in 2.5 mL filtrated IMDM medium and incubated for 20-30 min at RT to ensure cell attachment. Cells were monitored for 48 h in a HoloMonitor M4 live cell imaging system (Phase Holographic Imaging PHI AB, Lund, Sweden) acquiring holographic images every 10 min at 20× magnification. Each cell line was imaged from three different wells with one randomly chosen field of view per well (n = 3). Images were analyzed using HoloMonitor App Suite version 3.5.0.214. Single cells were automatically identified using auto minimum error with background threshold set to 120-140 and minimum object size set to 15-25. For analysis, images were manually examined to ensure correct cell number, size and morphology was identified, and out-of-focus images were removed. Cell proliferation analysis is based on cell count from images taken every 4^th^ h in the timeframe 0-42 h. Growth curve show mean ± SEM of three independent experiments. Cell proliferation was quantified by comparing relative increase in cell numbers at 42 h. Data are shown as mean ± SEM of three independent experiments pooled together and are expressed relative to WT (n=9). Significance was determined using one-way ANOVA with Dunnett’s correction.

##### Immunofluorescence

HAP1 cells were seeded on 12-mm glass coverslips (Paul Marienfeld GmbH) in 24-well plate (20,000-35,000 cells/well) and incubated overnight. Cells were fixed with 4% (w/v) paraformaldehyde in PBS buffer supplemented with 4% (w/v) sucrose for 15 min, washed with PBS, permeabilized with 0.1% Triton X-100 for 10 min, washed in PBS, and blocked with blocking solution (8% BSA and 2% goat serum in PBS) for 1 h. Cells were incubated with rabbit anti-COX IV (Cell Signaling, 4850, 1:200) or mouse anti-p62/SQSTM1 (Santa Cruz, sc-28359, 1:200) diluted in blocking solution for 1 h at RT in the dark. Subsequently, cells were washed with PBS and incubated with Alexa Fluor 594 goat anti-rabbit (Thermo Fisher Scientific, A-11012, 1:100) or Alexa Fluor 488 goat anti-mouse (Jackson ImmunoResearch, 115-547-003, 1:100) overnight at 4°C in the dark. Cells were washed four times in PBS for a total of 1 h followed by a final wash in water before the coverslips were mounted on a drop of ProLong antifade mountant with NucBlue stain (Invitrogen). Cells stained for COX IV were examined with a Zeiss Axiovert 200M widefield fluorescence microscope equipped with a AxioCam HRm camera and a Plan-Neofluar 100×/1.30 Ph3 oil-immersion objective (Carl Zeiss, Germany). Images were processed using ImageJ/Fiji version 2.1.0/1.53c (Schindelin et al., 2012). Around 200 cells per cell line were examined for mitochondrial morphology and categorized as either normal, fragmented, elongated or elongated + fragmented. Cells stained for p62/SQSTM1 were examined using a Andor Dragonfly 500 confocal microscopy (Oxford instruments) equipped with an iXon 888 Life EMCCD camera and a 100×/1.49 oil-immersion objective. Images were processed using Imaris version 9.7.2 (Oxford instruments).

HAP1 cells intended for siRNA treatment were seeded on ibiTreat μ-slide 4-well (ibidi GmbH) and incubated overnight. Then, cells were transfected with UBR1, UBR2 and UBR4 siRNA (siUBRs) or non-targeting control siRNA (siCtrl) as described earlier. Due to long incubation time from seeding to fixation (72 h post-transfection, 96 h post-seeding) cells were singularized by trypsinization (Gibco) 48 h post-transfection to increase spreading of the cells. Cells were prepared for immunofluorescent imaging as described earlier, using primary antibody rabbit anti-COX IV (Cell Signaling, 4850, 1:200) and Alexa 594-conjugated secondary antibody (Thermo Fisher Scientific, A-11012, 1:100) together with Rhodamine Phalloidin (Invitrogen, R415, 1:50). Samples were mounted by applying 4-5 drops of ibidi mounting medium with DAPI (ibidi GmbH) to each well. Cells were examined using a Andor Dragonfly 500 confocal microscopy (Oxford instruments) equipped with an iXon 888 Life EMCCD camera and a 100×/1.49 oil-immersion objective. Images were processed using IMARIS version 9.7.2. Cells were examined for mitochondrial morphology as earlier. Data are shown as mean ± SD of three independent experiments. Significance was determined using one-way ANOVA with Šídák’s correction. Around 200 cells per sample were examined.

##### *Drosophila* immunoprecipitation

For protein extracts, embryos with 0-4 hours expressing Naa30A-myc (CG11412) were dechorionated with 50% of bleach. Protein extraction was performed through homogenization of embryos in NB buffer (150 mM NaCl, 50 mM Tris–HCl pH 7.5, 2 mM EDTA, 0.1% NP-40, 1 mM DTT, 10 mM NaF, and EDTA-free protease inhibitor cocktail, Roche, Germany), and centrifugation at 20,000 × *g* for 3 min. Supernatant was collected and centrifuged twice. Total protein concentration was determined using Bio-Rad protein assay (Bio-Rad, Hercules, CA, USA) that is based on the Bradford dye-binding method.

For co-immunoprecipitation, proteins extracts (1.5 mg) of embryos expressing CG11412-myc were incubated with 1 μg of c-Myc antibody (Santa Cruz, sc-40, clone 9E10) for 1 h at 4°C. Subsequently, 0.9 mg of Dynabeads Protein G (Invitrogen) was added and incubated for 1 h at 4°C. After washing the beads 3 times with NB buffer, protein elution was performed with 100 μL of 100 mM Glycine pH 3.0 for 1 min and stopped with 10 μL of 1 M Tris Base pH 10.8. Proteins of the eluate were precipitated with 5 volumes of acetone at −20°C.

To determine the efficiency of Naa30A-myc immunoprecipitation, total protein extracts and eluted proteins were boiled for 5 min in SDS-PAGE sample buffer. Samples were centrifuged and loaded on a 10% SDS-gel for electrophoresis and transferred to a nitrocellulose membrane. Immunoblot analysis was performed according to standard protocols using the anti-myc antibody (Covance, PRB-150C-200, 1:2000).

##### Proteomic analysis of *Drosophila* proteins

Mass Spectrometry analysis was performed at the Mass Spectrometry Laboratory, Institute of Biochemistry and Biophysics, Polish Academy of Sciences, Warsaw, Poland. Briefly, peptides mixtures were analyzed by LC-MS-MS/MS (liquid chromatography coupled to tandem mass spectrometry) using Nano-Acquity (Waters, Milford, MA, USA) LC system and Orbitrap Velos mass spectrometer (Thermo Electron Corp., San Jose, CA, USA). Prior to analysis, proteins were subjected to standard “in-solution digestion” procedure, during which proteins were reduced with 100 mM DTT (for 30 min at 56°C), alkylated with 0.5 M iodoacetamide for 45 min in the dark at room temperature, and digested overnight with trypsin (Promega, Cat # V5111). The peptide mixture was applied to a RP-18 pre-column (nanoACQUITY Symmetry C18, Waters, 186003514) using water containing 0.1% TFA as mobile phase, then transferred to nano-HPLC RP-18 column (nanoACQUITY BEH C18, Waters, 186003545) using an acetonitrile gradient (0%-35% ACN in 180 min) in the presence of 0.05% formic acid with a flow rate of 250 nL/min. The column outlet was directly coupled to the ion source of the spectrometer, operating in the regime of data dependent MS to MS/MS switch. A blank run ensuring no cross contamination from previous samples preceded each analysis.

Raw data were processed by Mascot Distiller followed by Mascot Search (Matrix Science) against the Flybase database (FB2018_05). Search parameters for precursor and product ions mass tolerance were 15 ppm and 0.4 Da, respectively, enzyme specificity: trypsin, missed cleavage sites allowed: 0, fixed modification of cysteine by carbamidomethylation, and variable modification of methionine oxidation. Peptides with Mascot Score exceeding the threshold value corresponding to <5% False Positive Rate, calculated by Mascot procedure, and with the Mascot score above 30 were considered to be positively identified. Human orthologs were determined using DSRC Integrative Ortholog Prediction Tool (DIOPT) (Hu et al., 2011). Only scores above two were considered such as the best matches when there was more than one match per input.

##### *Drosophila* immunostaining and image analysis

Adult indirect flight muscles were prepared and stained as described in (Hunt and Demontis, 2013). Briefly, dissected hemithoraxes were fixed for 30 min with 4% formaldehyde in PBST (PBS with 0.2% Triton X-100). Samples were then rinsed 4 times with PBST and blocked for 30 min at room temperature with 5% BSA in PBST. Primary antibodies used were mouse anti-mono- and polyubiquitinylated conjugates (Enzo Life Sciences, BML-PW8810, Clone FK2, 1:250) and rabbit anti-Mef2 (Gift from Dr. Eileen Furlong, 1:200) and were diluted in 5% BSA in PBST. Samples were then rinsed 4 times with PBST and incubated with secondary antibody and/or stains diluted in 5% BSA in PBST for 3 h at room temperature. The secondary antibodies used for visualization of ubiquitin conjugates and Mef2 were anti-mouse Alexa Fluor 488 (Thermo Fischer, A-11029, 1:1000) and the anti-Rabbit Alexa Fluor 555 (Thermo Fisher, A-21428, 1:200), respectively. For F-actin visualization, Alexa-conjugated Phalloidin (Thermo Fisher, A12379, 1:200) or TRITC-conjugated Phalloidin (Sigma-Aldrich, P1951, 1:200) was used. Samples were then rinsed 4 times and mounted in Vectashield antifade mounting medium (Vector Labs) and imaged using a Leica SP8 confocal microscope or a Zeiss LSM710 confocal microscope. Protein aggregates were counted using Fiji (Schindelin et al., 2012) cell counter plugin and the number of aggregates were represented as the number of aggregates per area of muscle.

##### Longevity assay

A total of at least 50 males per genotype were used in this assay. Males were kept at a density of 6 to 8 flies per vial distributed by 8 tubes kept in an incubator at 25°C. Flies were transferred to fresh vials every 2^nd^ day, and viability was recorded at the time of vial transfer.

##### Climbing assay

The negative geotaxis assays were used to identify adult climbing defects. Flies were collected 0-3 days following eclosion and separated into batches of 10 in vials containing standard food media and were allowed to recover from the CO2 anesthesia for at least 24 hours. For the climbing assay, the flies from one vial were transferred to a test tube with 22 mm diameter with a line drawn at 8 cm from the bottom. Flies were tapped to the bottom to induce an innate climbing response. The number of flies that climbed 8 cm in 10 s and in 30 s was recorded. For each batch of 10 flies, the assay was repeated 10 times, allowing for 1 min rest period between each trial. Two independent biological replicas were tested per genotype with approximately 5 batches of 10 males for each replica.

##### Classifying fly climbing ability

To select *Naa30A* deletion males with good climbing ability (best climbers) and poor climbing ability (worst climbers), 200 *Naa30A* deletion males with 0-3 days after pupae eclosion were collected, separated into batches of 20 flies in food vials and transferred to fresh vials every 2 days until reaching the age of 7-10 days. The flies from each vial were transferred to test tube used for the climbing assay. Flies were tapped to the bottom to induce an innate climbing response, and the 9 males with best climbing ability (best climbing) and the 11 males with worst climbing ability were selected.

##### Copulation assay

The day before the assay, a single male (0-3 days) and a single virgin wild-type (OR) female (4-6 days) were introduced into a different food vial where they recovered from the CO2 anesthesia. After 24 hours the male and female were pooled and the number of males that were able to initiate copula within the first 10 min, 15 min, 20 min, or 30 min were recorded. Two independent biological replicas were tested per genotype. To avoid interference of the yellow mutation in the copulation success all males were carrying the yellow gene in the Y chromosome.

##### Male fertility

For measuring male fertility, a single male with 1-4 days were placed with five 5-7 days wild-type (Oregon-R (OR)) females into vials containing standard food media. Flies were allowed to mate for 2 days and then transferred to a fresh vial and allowed to mate for another 2 days. A male was classified as fertile when emerging larvae were observed in any of those tubes after an additional 2-day incubation. Two independent biological replicas were tested per genotype.

##### Flight assay

Flight assay was performed as previously described in (Greene et al., 2003). Briefly, an acetate sheet vertically divided into five different regions was coated with vacuum grease and inserted into a 1 L graduated cylinder and 2- or 20-days old males (after pupae eclosion) were dispensed into the apparatus by gently tapping vials containing 20 males into a funnel placed on top of the graduated cylinder Flies became stuck to the sheet where they landed. The sheet was removed, and the number of flies was counted in each of the five regions. The flight index was calculated as the weighted average of the region into which the flies landed. At least 100 flies of each genotype were tested.

##### Assessment of ATP levels

ATP levels were determined as described in (Lovero et al., 2018). Briefly, for each genotype groups of 5 males were collected in Eppendorf tubes and then homogenized in PBS using a potter homogenizer. An aliquot was incubated on ice with 12% perchloric acid for 1 min and then neutralized with a solution of 3 M K2CO3 and 2 M Tris. After centrifugation at 13,000 rpm for 1 min, ATP content was measured at 25°C, using the ATP Bioluminescence Assay kit (Sigma-Aldrich), according to manufacturer’s instruction. The luminescence was calculated as an average of three technical replicates and normalized to the total protein levels quantified using the Bradford’s method (Bio-Rad). Results are from the two or three biological replicas and are expressed as a percentage of the normalized luminescence of the control.

##### Mitochondrial morphology in *Drosophila*

Mitochondria morphology of the flight muscles were analyzed. To this purpose thoraxes from control and *Naa30A* deletion males expressing the mitochondrial marker mitoGFP by da-GAL4 driver were dissected and mitochondrial morphology imaged on a Zeiss LSM710 confocal microscope.

#### QUANTIFICATION AND STATISTICAL ANALYSIS

Image analysis of immunostaining of human cells and *Drosophila* muscles was performed using Imaris version 9.7.2 and Fiji software (Schindelin et al., 2012), respectively. Statistical analysis and graphical representations were performed using GraphPad Prism 9 software. Multiple comparisons were performed using one-way ANOVA. All results are obtained from at least two biological replicas. Sample size details are included in the respective figure, figure legend and methods.

### SUPPLEMENTAL INFORMATION

Supplementary references include the following: (Aksnes et al., 2013; Aregger et al., 2019; Beigl et al., 2020; Cha-Molstad et al., 2015; Cox and Mann, 2008; Drazic and Arnesen, 2017; Greene et al., 2003; Hu et al., 2011; Huang et al., 2005; Hunt and Demontis, 2013; Lovero et al., 2018; Miller et al., 1998; Perez-Riverol et al., 2019; Schindelin et al., 2012; Staes et al., 2011; Tyanova et al., 2016; Wodarz et al., 1995).

**Figure S1. Fitness effects of independent *NAA35*-KO replicate screens are highly reproducible.**

**(A)** Schematic representation of the NatC KO cell lines. HAP1 *NAA30*-KO, *NAA35*-KO and *NAA38*-KO cell lines were generated using CRISPR-Cas9 mediated genome editing. *NAA30*-KO cells have a 1 bp insertion in exon 2 (c.355_356insA) of the *NAA30* gene (Gene ID: 122830; CCDS32088). *NAA30*-KO cells could produce a putative protein product of 143 aa (p.(Asp119Glufs*26)) based on UniProt ID: Q147X3-1, which would be missing the catalytic GNAT domain (blue) containing the acetyl-CoA binding motif. *NAA35*-KO cells have a 1 bp deletion in exon 6 (c.370del) of the *NAA35* gene (Gene ID: 60560; CCDS 6673) and could generate a putative protein product of 140 aa (p.(Ser124Hisfs*18)) based on UniProt ID: Q5VZE5-1. *NAA38*-KO cells have a 32 bp deletion in the exon 1/intron border (NC_000017.11:g.7857000_7857031del) of the *NAA38* gene (Gene ID 83316; CCDS 11122). *NAA38*-KO cells could produce a putative protein product of 136 aa (p.(Phe132Cysfs*6) based on UniProt ID: Q9BRA0-2. Schematic is not scaled to size. Deletion (red), insertion (orange), GNAT domain (blue), new C-terminal sequence after frameshift mutation (grey). WT: wild-type; KO: knockout; bp: base pair; aa: amino acids. **(B)** Fitness effects were highly correlated across *NAA35* replicate screens. The reproducibility of the fitness effects (log2-fold change, LFC) of three independent *NAA35* screens was determined by calculating the Pearson correlation coefficient (*r*) between all possible pairwise combinations. **(C-D)** HAP1 *NAA38*-KO cells have increased cell proliferation. HAP1 WT and NatC KO cells were monitored in a HoloMonitor M4 imaging system for 42 h with image acquisition every 10 min. Single cell data in each experiment was obtained from one randomly selected field of view from three individual wells (n = 3). **(C)** Cell proliferation curves showing fold increase in cell number relative to 0 h (T0). Data are shown as mean ± SEM of three independent experiments pooled together (n = 3). **(D)** Cell proliferation was quantified by comparing relative increase in cell numbers between 0 and 42 h. Data are shown as mean ± SEM of three independent experiments expressed relative to WT. Mean cell proliferation WT = 105%, *NAA30*-KO = 82%, *NAA35*-KO = 98 %, and *NAA38*-KO = 149%. n = 9. **p < 0.01; one-way ANOVA with Dunnett’s correction. ns; not significant.

**Figure S2. Multiple sequence alignment of NAA30 and UBE2M/UbcE2M/UBC12.**

Multiple sequence alignment of **(A)** NAA30 and **(B)** UBE2M/UbcE2M/UBC12 orthologs from *Homo sapiens* (Hs), *Drosophila melanogaster* (Dm) and *Saccharomyces cerevisiae* (Sc). Secondary structures were predicted from yeast NatC structure (PDB: 6YGA) (Grunwald et al., 2020) and human UBE2M (PDB: 1Y8X) (Huang et al., 2005). Completely and highly conserved amino acids are shown in red and yellow boxes, respectively. Sequence alignment was performed using Clustal Omega, and secondary structures were visualized using ESPript 3.0 The acetyl-CoA binding motif (Q/RxxGxG/A) and residues Glu29, Glu118, and Tyr130, which are crucial for *in vitro* ScNAA30 activity, are indicated

**Figure S3. Knockdown of UBR4 partly restores UBE2M protein levels.**

**(A)** Immunoblot analysis of endogenous UBE2M, UBR4 and KCMF1 protein levels in HAP1 WT and *NAA30*-KO cells transfected with the indicated siRNAs for 72 h. **(B)** Immunoblot analysis of proteins representing positive GIs of NatC using total cell extract from HAP1 WT and NatC KO cells. **(C)** Heat map representation of proteins that were identified as significantly differentially regulated by UBR4 knockdown in at least 3 of 4 measurements and that passed an ANOVA multiple sample test (FDR = 0.01 and S0 = 0).

**Figure S4. NatC is required for normal mitochondrial phenotype, dehydrogenase activity and cell size.**

**(A)** Loss of NatC leads to elongated and fragmented mitochondria. HAP1 WT and NatC KO cells were stained with the mitochondrial marker anti-COX IV (red) and the nuclear marker NucBlue (blue) and analyzed by immunofluorescence. (Upper panel) Representative micrographs showing the different mitochondrial phenotypes observed in HAP1 *NAA30*-KO cells. (Lower panel) Same micrographs with zoom-in-frames of mitochondria. Cells were grouped into four bins based on mitochondrial morphology: normal, fragmented, elongated, and elongated + fragmented, see Figure 4A. At least 200 cells per cell line were examined. Scale bar, 10 μm. **(B)** NatC KO cells have increased cellular dehydrogenase activity. Cell viability of HAP1 cells was measured by WST-1 staining 24 h post-seeding. The WST-1 assay relies on cellular mitochondrial dehydrogenase activity. Data are shown as mean ± SD of four independent experiments with four technical replicates expressed relative to WT. **p < 0.01, ***p < 0.001; one-way ANOVA with Dunnett’s correction. **(C)** NatC KO cells show increased cell size. Median forward scatter area (FSC-A) indicating cell size were determined by flow cytometry. Data are shown as mean ± SD of three independent experiments. *p < 0.05, ***p < 0.001; one-way ANOVA with Dunnett’s correction. (*NAA30-*KO, p = 0.1171). **(D-G)** UBR knockdown in *NAA30*-KO cells rescues the mitochondrial phenotype. HAP1 WT and *NAA30*-KO cells were transfected with siCtrl or siUBR1, siUBR2 and siUBR4 (siUBRs), immunostained with anti-COX IV, and analyzed by immunofluorescence. Cells were grouped into four bins based on mitochondrial morphology: normal **(D)**, fragmented **(E)**, elongated **(F)**, and elongated + fragmented **(G)**. Data are shown as mean ± SD of three independent experiments. **p < 0.01, ****p < 0.0001; one-way ANOVA with Šídák’s correction. See combined data in Figure 4H. ns; not significant.

**Figure S5. Naa30A/CG11412 is the catalytic subunit of *Drosophila* NatC.**

**(A)** Schematic representation of *Drosophila melanogaster* Naa30A (also known as CG11412), and its orthologs, *Homo sapiens* NAA30 and *Saccharomyces cerevisiae* Mak3. Naa30A has a GNAT domain with an identity of 72.7% and 52.3% compared to human NAA30 and yeast Mak3, respectively. **(B)** Immunoprecipitation of Naa30A-Myc from *Drosophila* embryos total protein extracts. Immunoprecipitation using an anti-Myc antibody and embryos with 0 to 4 hours after egg laying. Embryos were collected from females expressing UAS-Naa30A-myc under the control of the nanos-Gal4 driver. The control embryos with 0 to 4 hours after egg laying were collected from wild-type (Oregon R; OR) females. Two different proteins corresponding to Naa30A-Myc could be detected by immunoblotting using anti-Myc antibody (arrows). **(C)** *Drosophila* Naa30A interacts with distinct orthologs of known auxiliary subunits of yeast and human NatC complexes. LC-MS/MS analysis of immunoprecipitated *Drosophila* Naa30A complexes from embryos expressing Naa30A-Myc and control wild-type embryos (no Myc-tagged Naa30A). (−), (+) and (++) correspond to 0, 1–9, and >10 unique peptide sequences, respectively. The complete list of proteins co-immunoprecipitated with Myc-tagged Naa30A is shown in **Table S6**. **(D)** *Drosophila* Naa30A rescues Arl3-GFP localization phenotype in yeast Naa30/Mak3 deletion strain. Arl3-GFP localization in wild-type yeast cells, *naa30Δ* yeast cells and *naa30Δ* yeast cells expressing *Drosophila* Naa30A. Scale bar, 2 μm.

**Figure S6. *Drosophila* Naa30A is not required for adult viability.**

**(A)** <u>Top:</u> Schematic representation of the wild-type *Drosophila Naa30A* gene and the two annotated unique transcripts, Naa30A-RA and Naa30A-RD. <u>Bottom</u>: Schematic representation of the *Naa30A* deletion of the first 302 amino acids (almost all coding sequence), with an additional frameshift mutation within the remaining coding sequence and a stop codon in the newly generated position 20. **(B)** Images of control (y^1^/Y), and *Naa30A* deletion males (y^1^, Naa30A^Δ74^/Y) showing absence of any obvious developmental defects.

**Figure S7. *Drosophila* Naa30A is not obviously required for mitochondrial function.**

**(A)** Mitochondrial network in indirect flight muscles from young (1-4 days after pupae eclosion) control males (y^1^/Y) or *Naa30A* deletion males (y^1^, Naa30A^Δ74^/Y) labelled with UAS-mito-GFP expressed under control of the ubiquitous daughterless-Gal4 (da-GAL4) driver. Results are shown for indirect flight muscles of three different thoraxes. Scale bar is 5 μm. **(B)** Total ATP levels in young (2 days after pupae eclosion) and old (10 days after pupae eclosion) control males (y^1^/Y), *Naa30A* deletion males (y^1^, Naa30A^Δ74^/Y) and *Naa30A* deletion males carrying a *Naa30A* genomic rescue construct (y^1^, Naa30A^Δ74^/Y;; gNaa30A-myc/+). Results are mean ± SEM of two (young flies) or three (old flies) independent experiments. *p<0.01, **p<0.001,***p<0.0001; one-way ANOVA with Dunnett’s correction.

**Table S1. Genome-wide CRISPR knockout screens in HAP1 cells.**

Quantitative Genetic Interaction (qGI) scores for genome-wide CRISPR knockout screens in HAP WT, *NAA3*5-KO, *NAA30*-KO, and *NAA38*-KO cells using TKOv3 gRNA library.

**Table S2. Biological pathway (BP) enrichment analysis of negative and positive genetic interactions of *NAA35*, *NAA30*, and *NAA38*.**

**Table S3. N-terminal acetylome analysis of HAP1 WT, *NAA30*-KO, *NAA35*-KO and *NAA38*-KO cells.**

Protein extracts were digested using trypsin followed by enrichment of *in vivo* acetylated N-terminal peptides using strong cation exchange (SCX) chromatography.

**Table S4. Label-free quantitative (LFQ) shotgun proteomic analysis of HAP1 WT, *NAA30*-KO, *NAA35*-KO, and *NAA38*-KO cells.**

The impact of NatC subunits NAA30, NAA35, and NAA38 on the human proteome (protein abundance) was determined by LFQ shotgun proteomics analysis by co-digesting protein extracts with Lys-C and trypsin.

**Table S5. Knockdown of UBR4 rescues *NAA30*-KO induced changes in protein abundance.**

HAP1 WT and *NAA30*-KO cells were transfected with siUBR4 or siCtrl for 72 h and protein abundance was determined by TMT-based quantitative proteomics.

**Table S6. List of proteins co-immunoprecipitated with Myc-tagged *Drosophila* Naa30A from embryonic protein extracts.**

Anti-Myc immunoprecipitation of Naa30A complexes using protein extracts from *Drosophila* embryos expressing Naa30A-Myc or from wild-type embryos (not expressing the Myc-tagged Naa30A; negative control (−)). Immunoprecipitated proteins were identified by LC-MS and data were searched against the Flybase database (FB2018_05). Two replica experiments were performed for each condition. Proteins identified in the negative control (−) were discarded as being due to non-specific interactions with the anti-Myc antibody and/or the magnetic beads used.

**Table S7. List of primers and peptides used in this study.**

**Movie S1. *Drosophila Naa30A* deletion males show significantly slower motility (climbing).**

Climbing of control males (y^1^/Y), *Naa30A* deletion males (y^1^, Naa30A^Δ74^/Y) and *Naa30A* deletion males carrying a *Naa30A* genomic rescue construct (y^1^, Naa30^Δ74^/Y;; gNaa30-myc/+).

## Notes

### Competing Interest Statement

Jason Moffat is a shareholder and advisor of Century Therapeutics and Aelian Biotechnology.

## REFERENCES

Aksnes, H., Drazic, A., Marie, M., and Arnesen, T. (2016). First Things First: Vital Protein Marks by N-Terminal Acetyltransferases. Trends Biochem Sci 41, 746–760.

Aksnes, H., Osberg, C., and Arnesen, T. (2013). N-terminal acetylation by NatC is not a general determinant for substrate subcellular localization in Saccharomyces cerevisiae. Plos One 8, e61012.

Aksnes, H., Ree, R., and Arnesen, T. (2019). Co-translational, Post-translational, and Non-catalytic Roles of N-Terminal Acetyltransferases. Mol Cell 73, 1097–1114.

Aksnes, H., Van Damme, P., Goris, M., Starheim, K.K., Marie, M., Stove, S.I., Hoel, C., Kalvik, T.V., Hole, K., Glomnes, N., et al. (2015). An Organellar N alpha-Acetyltransferase, Naa60, Acetylates Cytosolic N Termini of Transmembrane Proteins and Maintains Golgi Integrity. Cell Reports 10, 1362–1374.

Aregger, M., Chandrashekhar, M., Tong, A.H.Y., Chan, K., and Moffat, J. (2019). Pooled Lentiviral CRISPR-Cas9 Screens for Functional Genomics in Mammalian Cells. Methods Mol Biol 1869, 169–188.

Aregger, M., Lawson, K.A., Billmann, M., Costanzo, M., Tong, A.H.Y., Chan, K., Rahman, M., Brown, K.R., Ross, C., Usaj, M., et al. (2020). Systematic mapping of genetic interactions for de novo fatty acid synthesis identifies C12orf49 as a regulator of lipid metabolism. Nat Metab 2, 499–513.

Arnesen, T., Van Damme, P., Polevoda, B., Helsens, K., Evjenth, R., Colaert, N., Varhaug, J.E., Vandekerckhove, J., Lillehaug, J.R., Sherman, F., et al. (2009). Proteomics analyses reveal the evolutionary conservation and divergence of N-terminal acetyltransferases from yeast and humans. Proc Natl Acad Sci U S A 106, 8157–8162.

Ayyadevara, S., Balasubramaniam, M., Suri, P., Mackintosh, S.G., Tackett, A.J., Sullivan, D.H., Shmookler Reis, R.J., and Dennis, R.A. (2016). Proteins that accumulate with age in human skeletal-muscle aggregates contribute to declines in muscle mass and function in Caenorhabditis elegans. Aging (Albany NY) 8, 3486–3497.

Bachmair, A., Finley, D., and Varshavsky, A. (1986). Invivo Half-Life of a Protein Is a Function of Its Amino-Terminal Residue. Science 234, 179–186.

Behnia, R., Panic, B., Whyte, J.R.C., and Munro, S. (2004). Targeting of the arf-like GTPase Arl3p to the Golgi requires N-terminal acetylation and the membrane protein Sys1p. Nature Cell Biology 6, 405-+.

Beigl, T.B., Kjosas, I., Seljeseth, E., Glomnes, N., and Aksnes, H. (2020). Efficient and crucial quality control of HAP1 cell ploidy status. Biol Open 9.

Billmann, M., Ward, H.N., Aregger, M., Costanzo, M., Andrews, B.J., Boone, C., Moffat, J., and Myers, C.L. (2022). Reproducibility metrics for CRISPR screens. bioRxiv, 2022.2002.2019.480892.

Blomen, V.A., Majek, P., Jae, L.T., Bigenzahn, J.W., Nieuwenhuis, J., Staring, J., Sacco, R., van Diemen, F.R., Olk, N., Stukalov, A., et al. (2015). Gene essentiality and synthetic lethality in haploid human cells. Science 350, 1092–1096.

Brand, A.H., and Perrimon, N. (1993). Targeted gene expression as a means of altering cell fates and generating dominant phenotypes. Development 118, 401–415.

Cao, J.Y., Poddar, A., Magtanong, L., Lumb, J.H., Mileur, T.R., Reid, M.A., Dovey, C.M., Wang, J., Locasale, J.W., Stone, E., et al. (2019). A Genome-wide Haploid Genetic Screen Identifies Regulators of Glutathione Abundance and Ferroptosis Sensitivity. Cell Rep 26, 1544–1556 e1548.

Cha-Molstad, H., Sung, K.S., Hwang, J., Kim, K.A., Yu, J.E., Yoo, Y.D., Jang, J.M., Han, D.H., Molstad, M., Kim, J.G., et al. (2015). Amino-terminal arginylation targets endoplasmic reticulum chaperone BiP for autophagy through p62 binding. Nat Cell Biol 17, 917–929.

Chen, S.J., Wu, X., Wadas, B., Oh, J.H., and Varshavsky, A. (2017). An N-end rule pathway that recognizes proline and destroys gluconeogenic enzymes. Science 355.

Cheng, H., Dharmadhikari, A.V., Varland, S., Ma, N., Domingo, D., Kleyner, R., Rope, A.F., Yoon, M., Stray-Pedersen, A., Posey, J.E., et al. (2018). Truncating Variants in NAA15 Are Associated with Variable Levels of Intellectual Disability, Autism Spectrum Disorder, and Congenital Anomalies. Am J Hum Genet 102, 985–994.

Costanzo, M., Baryshnikova, A., Bellay, J., Kim, Y., Spear, E.D., Sevier, C.S., Ding, H., Koh, J.L., Toufighi, K., Mostafavi, S., et al. (2010). The genetic landscape of a cell. Science 327, 425–431.

Costanzo, M., Kuzmin, E., van Leeuwen, J., Mair, B., Moffat, J., Boone, C., and Andrews, B. (2019). Global Genetic Networks and the Genotype-to-Phenotype Relationship. Cell 177, 85–100.

Costanzo, M., VanderSluis, B., Koch, E.N., Baryshnikova, A., Pons, C., Tan, G.H., Wang, W., Usaj, M., Hanchard, J., Lee, S.D., et al. (2016). A global genetic interaction network maps a wiring diagram of cellular function. Science 353.

Cox, J., and Mann, M. (2008). MaxQuant enables high peptide identification rates, individualized p.p.b.-range mass accuracies and proteome-wide protein quantification. Nat Biotechnol 26, 1367–1372.

Degroot, R.J., Rumenapf, T., Kuhn, R.J., Strauss, E.G., and Strauss, J.H. (1991). Sindbis Virus-Rna Polymerase Is Degraded by the N-End Rule Pathway. P Natl Acad Sci USA 88, 8967–8971.

Demarchi, F., and Schneider, C. (2007). The calpain system as a modulator of stress/damage response. Cell Cycle 6, 136–138.

Demontis, F., and Perrimon, N. (2010). FOXO/4E-BP signaling in Drosophila muscles regulates organism-wide proteostasis during aging. Cell 143, 813–825.

Demontis, F., Piccirillo, R., Goldberg, A.L., and Perrimon, N. (2013). Mechanisms of skeletal muscle aging: insights from Drosophila and mammalian models. Dis Model Mech 6, 1339–1352.

Deng, S.B., Gottlieb, L., Pan, B.Y., Supplee, J., Wei, X.P., Petersson, J., and Marmorstein, R. (2021). Molecular mechanism of N-terminal acetylation by the ternary NatC complex. Structure 29, 1094-+.

Dong, C., Chen, S.J., Melnykov, A., Weirich, S., Sun, K., Jeltsch, A., Varshavsky, A., and Min, J. (2020). Recognition of nonproline N-terminal residues by the Pro/N-degron pathway. Proc Natl Acad Sci U S A 117, 14158–14167.

Drazic, A., Aksnes, H., Marie, M., Boczkowska, M., Varland, S., Timmerman, E., Foyn, H., Glomnes, N., Rebowski, G., Impens, F., et al. (2018). NAA80 is actin’s N-terminal acetyltransferase and regulates cytoskeleton assembly and cell motility. Proc Natl Acad Sci U S A 115, 4399–4404.

Drazic, A., and Arnesen, T. (2017). [(14)C]-Acetyl-Coenzyme A-Based In Vitro N-Terminal Acetylation Assay. Methods Mol Biol 1574, 1–8.

Enchev, R.I., Schulman, B.A., and Peter, M. (2015). Protein neddylation: beyond cullin-RING ligases. Nat Rev Mol Cell Biol 16, 30–44.

Feany, M.B., and Bender, W.W. (2000). A Drosophila model of Parkinson’s disease. Nature 404, 394–398.

Friedrich, U.A., Zedan, M., Hessling, B., Fenzl, K., Gillet, L., Barry, J., Knop, M., Kramer, G., and Bukau, B. (2021). N(alpha)-terminal acetylation of proteins by NatA and NatB serves distinct physiological roles in Saccharomyces cerevisiae. Cell Rep 34, 108711.

Gargano, J.W., Martin, I., Bhandari, P., and Grotewiel, M.S. (2005). Rapid iterative negative geotaxis (RING): a new method for assessing age-related locomotor decline in Drosophila. Exp Gerontol 40, 386–395.

Gillingham, A.K., and Munro, S. (2016). Finding the Golgi: Golgin Coiled-Coil Proteins Show the Way. Trends Cell Biol 26, 399–408.

Greene, J.C., Whitworth, A.J., Kuo, I., Andrews, L.A., Feany, M.B., and Pallanck, L.J. (2003). Mitochondrial pathology and apoptotic muscle degeneration in Drosophila parkin mutants. Proc Natl Acad Sci U S A 100, 4078–4083.

Grunwald, S., Hopf, L.V.M., Bock-Bierbaum, T., Lally, C.C.M., Spahn, C.M.T., and Daumke, O. (2020). Divergent architecture of the heterotrimeric NatC complex explains N-terminal acetylation of cognate substrates. Nature Communications 11.

Gunage, R.D., Dhanyasi, N., Reichert, H., and VijayRaghavan, K. (2017). Drosophila adult muscle development and regeneration. Semin Cell Dev Biol 72, 56–66.

Hart, T., Chandrashekhar, M., Aregger, M., Steinhart, Z., Brown, K.R., MacLeod, G., Mis, M., Zimmermann, M., Fradet-Turcotte, A., Sun, S., et al. (2015). High-Resolution CRISPR Screens Reveal Fitness Genes and Genotype-Specific Cancer Liabilities. Cell 163, 1515–1526.

Hart, T., Tong, A.H.Y., Chan, K., Van Leeuwen, J., Seetharaman, A., Aregger, M., Chandrashekhar, M., Hustedt, N., Seth, S., Noonan, A., et al. (2017). Evaluation and Design of Genome-Wide CRISPR/SpCas9 Knockout Screens. G3 (Bethesda) 7, 2719–2727.

Helsens, K., Colaert, N., Barsnes, H., Muth, T., Flikka, K., Staes, A., Timmerman, E., Wortelkamp, S., Sickmann, A., Vandekerckhove, J., et al. (2010). ms_lims, a simple yet powerful open source laboratory information management system for MS-driven proteomics. Proteomics 10, 1261–1264.

Heo, A.J., Kim, S.B., Ji, C.H., Han, D., Lee, S.J., Lee, S.H., Lee, M.J., Lee, J.S., Ciechanover, A., Kim, B.Y., et al. (2021). The N-terminal cysteine is a dual sensor of oxygen and oxidative stress. P Natl Acad Sci USA 118.

Hesse, D., Jaschke, A., Kanzleiter, T., Witte, N., Augustin, R., Hommel, A., Puschel, G.P., Petzke, K.J., Joost, H.G., Schupp, M., et al. (2012). GTPase ARFRP1 Is Essential for Normal Hepatic Glycogen Storage and Insulin-Like Growth Factor 1 Secretion. Molecular and Cellular Biology 32, 4363–4374.

Hofmann, I., and Munro, S. (2006). An N-terminally acetylated Arf-like GTPase is localised to lysosomes and affects their motility. J Cell Sci 119, 1494–1503.

Homma, Y., Hiragi, S., and Fukuda, M. (2021). Rab family of small GTPases: an updated view on their regulation and functions. Febs J 288, 36–55.

Hommel, A., Hesse, D., Volker, W., Jaschke, A., Moser, M., Engel, T., Bluher, M., Zahn, C., Chadt, A., Ruschke, K., et al. (2010). The ARF-Like GTPase ARFRP1 Is Essential for Lipid Droplet Growth and Is Involved in the Regulation of Lipolysis. Molecular and Cellular Biology 30, 1231–1242.

Hong, J.H., Kaustov, L., Coyaud, E., Srikumar, T., Wan, J., Arrowsmith, C., and Raught, B. (2015). KCMF1 (potassium channel modulatory factor 1) Links RAD6 to UBR4 (ubiquitin N-recognin domain-containing E3 ligase 4) and lysosome-mediated degradation. Mol Cell Proteomics 14, 674–685.

Hu, R.G., Sheng, J., Qi, X., Xu, Z.M., Takahashi, T.T., and Varshavsky, A. (2005). The N-end rule pathway as a nitric oxide sensor controlling the levels of multiple regulators. Nature 437, 981–986.

Hu, Y.H., Flockhart, I., Vinayagam, A., Bergwitz, C., Berger, B., Perrimon, N., and Mohr, S.E. (2011). An integrative approach to ortholog prediction for disease-focused and other functional studies. Bmc Bioinformatics 12.

Huang, D.T., Ayrault, O., Hunt, H.W., Taherbhoy, A.M., Duda, D.M., Scott, D.C., Borg, L.A., Neale, G., Murray, P.J., Roussel, M.F., et al. (2009). E2-RING expansion of the NEDD8 cascade confers specificity to cullin modification. Mol Cell 33, 483–495.

Huang, D.T., Paydar, A., Zhuang, M., Waddell, M.B., Holton, J.M., and Schulman, B.A. (2005). Structural basis for recruitment of Ubc12 by an E2 binding domain in NEDD8’s E1. Mol Cell 17, 341–350.

Hunt, L.C., and Demontis, F. (2013). Whole-mount immunostaining of Drosophila skeletal muscle. Nat Protoc 8, 2496–2501.

Hunt, L.C., Schadeberg, B., Stover, J., Haugen, B., Pagala, V., Wang, Y.D., Puglise, J., Barton, E.R., Peng, J., and Demontis, F. (2021). Antagonistic control of myofiber size and muscle protein quality control by the ubiquitin ligase UBR4 during aging. Nat Commun 12, 1418.

Hunt, L.C., Stover, J., Haugen, B., Shaw, T.I., Li, Y., Pagala, V.R., Finkelstein, D., Barton, E.R., Fan, Y., Labelle, M., et al. (2019). A Key Role for the Ubiquitin Ligase UBR4 in Myofiber Hypertrophy in Drosophila and Mice. Cell Rep 28, 1268–1281 e1266.

Hwang, C.S., Shemorry, A., and Varshavsky, A. (2010). N-terminal acetylation of cellular proteins creates specific degradation signals. Science 327, 973–977.

Jahn, T.R., Kohlhoff, K.J., Scott, M., Tartaglia, G.G., Lomas, D.A., Dobson, C.M., Vendruscolo, M., and Crowther, D.C. (2011). Detection of early locomotor abnormalities in a Drosophila model of Alzheimer’s disease. J Neurosci Methods 197, 186–189.

Jang, J.H. (2004). FIGC, a novel FGF-induced ubiquitin-protein ligase in gastric cancers. Febs Lett 578, 21–25.

Kats, I., Khmelinskii, A., Kschonsak, M., Huber, F., Kniess, R.A., Bartosik, A., and Knop, M. (2018). Mapping Degradation Signals and Pathways in a Eukaryotic N-terminome. Molecular Cell 70, 488-+.

Kats, I., Reinbold, C., Kschonsak, M., Khmelinskii, A., Armbruster, L., Ruppert, T., and Knop, M. (2022). Up-regulation of ubiquitin-proteasome activity upon loss of NatA-dependent N-terminal acetylation. Life Sci Alliance 5.

Kim, A.Y., Bommelje, C.C., Lee, B.E., Yonekawa, Y., Choi, L., Morris, L.G., Huang, G.C., Kaufman, A., Ryan, R.J.H., Hao, B., et al. (2008). SCCRO (DCUN1D1) Is an Essential Component of the E3 Complex for Neddylation. Journal of Biological Chemistry 283, 33211–33220.

Kim, H.K., Kim, R.R., Oh, J.H., Cho, H., Varshavsky, A., and Hwang, C.S. (2014). The N-Terminal Methionine of Cellular Proteins as a Degradation Signal. Cell 156, 158–169.

Kim, S.T., Lee, Y.J., Tasaki, T., Mun, S.R., Hwang, J., Kang, M.J., Ganipisetti, S., Yi, E.C., Kim, B.Y., and Kwon, Y.T. (2018). The N-recognin UBR4 of the N-end rule pathway is targeted to and required for the biogenesis of the early endosome. J Cell Sci 131.

Kurz, T., Chou, Y.C., WillemS, A.R., Meyer-Schaller, N., Hecht, M.L., Tyers, M., Peter’, M., and Sicheri, F. (2008). Dcn1 functions as a scaffold-type E3 ligase for cullin neddylation. Molecular Cell 29, 23–35.

Lee, M.J., Tasaki, T., Moroi, K., An, J.Y., Kimura, S., Davydov, I.V., and Kwon, Y.T. (2005). RGS4 and RGS5 are in vivo substrates of the N-end rule pathway. Proc Natl Acad Sci U S A 102, 15030–15035.

Lin, R., Tao, R., Gao, X., Li, T., Zhou, X., Guan, K.L., Xiong, Y., and Lei, Q.Y. (2013). Acetylation stabilizes ATP-citrate lyase to promote lipid biosynthesis and tumor growth. Mol Cell 51, 506–518.

Linster, E., Forero Ruiz, F.L., Miklankova, P., Ruppert, T., Mueller, J., Armbruster, L., Gong, X., Serino, G., Mann, M., Hell, R., et al. (2022). Cotranslational N-degron masking by acetylation promotes proteome stability in plants. Nat Commun 13, 810.

Lovero, D., Giordano, L., Marsano, R.M., Sanchez-Martinez, A., Boukhatmi, H., Drechsler, M., Oliva, M., Whitworth, A.J., Porcelli, D., and Caggese, C. (2018). Characterization of Drosophila ATPsynC mutants as a new model of mitochondrial ATP synthase disorders. Plos One 13, e0201811.

Matta-Camacho, E., Kozlov, G., Li, F.F., and Gehring, K. (2010). Structural basis of substrate recognition and specificity in the N-end rule pathway. Nature Structural & Molecular Biology 17, 1182-+.

Miller, C.A., 3rd, Martinat, M.A., and Hyman, L.E. (1998). Assessment of aryl hydrocarbon receptor complex interactions using pBEVY plasmids: expressionvectors with bi-directional promoters for use in Saccharomyces cerevisiae. Nucleic Acids Res 26, 3577–3583.

Monda, J.K., Scott, D.C., Miller, D.J., Lydeard, J., King, D., Harper, J.W., Bennett, E.J., and Schulman, B.A. (2013). Structural Conservation of Distinctive N-terminal Acetylation-Dependent Interactions across a Family of Mammalian NEDD8 Ligation Enzymes. Structure 21, 42–53.

Morrison, J., Altuwaijri, N.K., Bronstad, K., Aksnes, H., Alsaif, H.S., Evans, A., Hashem, M., Wheeler, P.G., Webb, B.D., Alkuraya, F.S., et al. (2021). Missense NAA20 variants impairing the NatB protein N-terminal acetyltransferase cause autosomal recessive developmental delay, intellectual disability, and microcephaly. Genet Med 23, 2213–2218.

Mueller, F., Friese, A., Pathe, C., da Silva, R.C., Rodriguez, K.B., Musacchio, A., and Bange, T. (2021). Overlap of NatA and IAP substrates implicates N-terminal acetylation in protein stabilization. Sci Adv 7.

Muffels, I.J.J., Wiame, E., Fuchs, S.A., Massink, M.P.G., Rehmann, H., Musch, J.L.I., Van Haaften, G., Vertommen, D., van Schaftingen, E., and van Hasselt, P.M. (2021). NAA80 bi-allelic missense variants result in high-frequency hearing loss, muscle weakness and developmental delay. Brain Commun 3, fcab256.

Mughal, A.A., Grieg, Z., Skjellegrind, H., Fayzullin, A., Lamkhannat, M., Joel, M., Ahmed, M.S., Murrell, W., Vik-Mo, E.O., Langmoen, I.A., et al. (2015). Knockdown of NAT12/NAA30 reduces tumorigenic features of glioblastoma-initiating cells. Mol Cancer 14.

Nair, K.S. (2005). Aging muscle. Am J Clin Nutr 81, 953–963.

Oh, J.H., Hyun, J.Y., and Varshavsky, A. (2017). Control of Hsp90 chaperone and its clients by N-terminal acetylation and the N-end rule pathway. P Natl Acad Sci USA 114, E4370–E4379.

Osberg, C., Aksnes, H., Ninzima, S., Marie, M., and Arnesen, T. (2016). Microscopy-based Saccharomyces cerevisiae complementation model reveals functional conservation and redundancy of N-terminal acetyltransferases. Sci Rep 6, 31627.

Park, S.E., Kim, J.M., Seok, O.H., Cho, H., Wadas, B., Kim, S.Y., Varshavsky, A., and Hwang, C.S. (2015). Control of mammalian G protein signaling by N-terminal acetylation and the N-end rule pathway. Science 347, 1249–1252.

Perez-Riverol, Y., Csordas, A., Bai, J., Bernal-Llinares, M., Hewapathirana, S., Kundu, D.J., Inuganti, A., Griss, J., Mayer, G., Eisenacher, M., et al. (2019). The PRIDE database and related tools and resources in 2019: improving support for quantification data. Nucleic Acids Res 47, D442–D450.

Pesaresi, P., Gardner, N.A., Masiero, S., Dietzmann, A., Eichacker, L., Wickner, R., Salamini, F., and Leister, D. (2003). Cytoplasmic N-terminal protein acetylation is required for efficient photosynthesis in Arabidopsis. Plant Cell 15, 1817–1832.

Polevoda, B., and Sherman, F. (2001). NatC N-alpha-terminal acetyltransferase of yeast contains three subunits, Mak3p, Mak10p, and Mak31p. Journal of Biological Chemistry 276, 20154–20159.

Ree, R., Varland, S., and Arnesen, T. (2018). Spotlight on protein N-terminal acetylation. Exp Mol Med 50, 1–13.

Rope, A.F., Wang, K., Evjenth, R., Xing, J., Johnston, J.J., Swensen, J.J., Johnson, W.E., Moore, B., Huff, C.D., Bird, L.M., et al. (2011). Using VAAST to identify an X-linked disorder resulting in lethality in male infants due to N-terminal acetyltransferase deficiency. Am J Hum Genet 89, 28–43.

Rorth, P. (1998). Gal4 in the Drosophila female germline. Mech Dev 78, 113–118.

Schindelin, J., Arganda-Carreras, I., Frise, E., Kaynig, V., Longair, M., Pietzsch, T., Preibisch, S., Rueden, C., Saalfeld, S., Schmid, B., et al. (2012). Fiji: an open-source platform for biological-image analysis. Nat Methods 9, 676–682.

Schuster, C.M., Davis, G.W., Fetter, R.D., and Goodman, C.S. (1996). Genetic dissection of structural and functional components of synaptic plasticity. I. Fasciclin II controls synaptic stabilization and growth. Neuron 17, 641–654.

Scott, D.C., Hammill, J.T., Min, J., Rhee, D.Y., Connelly, M., Sviderskiy, V.O., Bhasin, D., Chen, Y.Z., Ong, S.S., Chai, S.C., et al. (2017). Blocking an N-terminal acetylation-dependent protein interaction inhibits an E3 ligase. Nature Chemical Biology 13, 850-+.

Scott, D.C., Monda, J.K., Bennett, E.J., Harper, J.W., and Schulman, B.A. (2011). N-Terminal Acetylation Acts as an Avidity Enhancer Within an Interconnected Multiprotein Complex. Science 334, 674–678.

Scott, D.C., Monda, J.K., Grace, C.R., Duda, D.M., Kriwacki, R.W., Kurz, T., and Schulman, B.A. (2010). A dual E3 mechanism for Rub1 ligation to Cdc53. Mol Cell 39, 784–796.

Setty, S.R., Strochlic, T.I., Tong, A.H., Boone, C., and Burd, C.G. (2004). Golgi targeting of ARF-like GTPase Arl3p requires its Nalpha-acetylation and the integral membrane protein Sys1p. Nat Cell Biol 6, 414–419.

Shemorry, A., Hwang, C.S., and Varshavsky, A. (2013). Control of Protein Quality and Stoichiometries by N-Terminal Acetylation and the N-End Rule Pathway. Molecular Cell 50, 540–551.

Sherpa, D., Chrustowicz, J., and Schulman, B.A. (2022). How the ends signal the end: Regulation by E3 ubiquitin ligases recognizing protein termini. Mol Cell 82, 1424–1438.

Staes, A., Impens, F., Van Damme, P., Ruttens, B., Goethals, M., Demol, H., Timmerman, E., Vandekerckhove, J., and Gevaert, K. (2011). Selecting protein N-terminal peptides by combined fractional diagonal chromatography. Nat Protoc 6, 1130–1141.

Starheim, K.K., Gromyko, D., Evjenth, R., Ryningen, A., Varhaug, J.E., Lillehaug, J.R., and Arnesen, T. (2009). Knockdown of human N alpha-terminal acetyltransferase complex C leads to p53-dependent apoptosis and aberrant human Arl8b localization. Mol Cell Biol 29, 3569–3581.

Starheim, K.K., Kalvik, T.V., Bjorkoy, G., and Arnesen, T. (2017). Depletion of the human N-terminal acetyltransferase hNaa30 disrupts Golgi integrity and ARFRP1 localization. Bioscience Rep 37.

Tasaki, T., Kim, S.T., Zakrzewska, A., Lee, B.E., Kang, M.J., Yoo, Y.D., Cha-Molstad, H.J., Hwang, J., Soung, N.K., Sung, K.S., et al. (2013). UBR box N-recognin-4 (UBR4), an N-recognin of the N-end rule pathway, and its role in yolk sac vascular development and autophagy. P Natl Acad Sci USA 110, 3800–3805.

Tasaki, T., Mulder, L.C.F., Iwamatsu, A., Lee, M.J., Davydov, I.V., Varshavsky, A., Muesing, M., and Kwon, Y.T. (2005). A family of mammalian E3 ubiquitin ligases that contain the UBR box motif and recognize N-degrons. Molecular and Cellular Biology 25, 7120–7136.

Tasaki, T., Zakrzewska, A., Dudgeon, D.D., Jiang, Y.H., Lazo, J.S., and Kwon, Y.T. (2009). The Substrate Recognition Domains of the N-end Rule Pathway. Journal of Biological Chemistry 284, 1884–1895.

Tercero, J.C., Dinman, J.D., and Wickner, R.B. (1993). Yeast MAK3 N-acetyltransferase recognizes the N-terminal four amino acids of the major coat protein (gag) of the L-A double-stranded RNA virus. J Bacteriol 175, 3192–3194.

Timms, R.T., Zhang, Z., Rhee, D.Y., Harper, J.W., Koren, I., and Elledge, S.J. (2019). A glycine-specific N-degron pathway mediates the quality control of protein N-myristoylation. Science 365.

Tyanova, S., Temu, T., Sinitcyn, P., Carlson, A., Hein, M.Y., Geiger, T., Mann, M., and Cox, J. (2016). The Perseus computational platform for comprehensive analysis of (prote)omics data. Nat Methods 13, 731–740.

Van Damme, P., Evjenth, R., Foyn, H., Demeyer, K., De Bock, P.J., Lillehaug, J.R., Vandekerckhove, J., Arnesen, T., and Gevaert, K. (2011a). Proteome-derived peptide libraries allow detailed analysis of the substrate specificities of N(alpha)-acetyltransferases and point to hNaa10p as the post-translational actin N(alpha)-acetyltransferase. Mol Cell Proteomics 10, M110 004580.

Van Damme, P., Hole, K., Gevaert, K., and Arnesen, T. (2015). N-terminal acetylome analysis reveals the specificity of Naa50 (Nat5) and suggests a kinetic competition between N-terminal acetyltransferases and methionine aminopeptidases. Proteomics 15, 2436–2446.

Van Damme, P., Hole, K., Pimenta-Marques, A., Helsens, K., Vandekerckhove, J., Martinho, R.G., Gevaert, K., and Arnesen, T. (2011b). NatF contributes to an evolutionary shift in protein N-terminal acetylation and is important for normal chromosome segregation. PLoS Genet 7, e1002169.

Van Damme, P., Kalvik, T.V., Starheim, K.K., Jonckheere, V., Myklebust, L.M., Menschaert, G., Varhaug, J.E., Gevaert, K., and Arnesen, T. (2016). A Role for Human N-alpha Acetyltransferase 30 (Naa30) in Maintaining Mitochondrial Integrity. Molecular & Cellular Proteomics 15, 3361–3372.

Van Damme, P., Lasa, M., Polevoda, B., Gazquez, C., Elosegui-Artola, A., Kim, D.S., De Juan-Pardo, E., Demeyer, K., Hole, K., Larrea, E., et al. (2012). N-terminal acetylome analyses and functional insights of the N-terminal acetyltransferase NatB. P Natl Acad Sci USA 109, 12449–12454.

Van Doren, M., Williamson, A.L., and Lehmann, R. (1998). Regulation of zygotic gene expression in Drosophila primordial germ cells. Curr Biol 8, 243–246.

Varland, S., Aksnes, H., Kryuchkov, F., Impens, F., Van Haver, D., Jonckheere, V., Ziegler, M., Gevaert, K., Van Damme, P., and Arnesen, T. (2018a). N-terminal Acetylation Levels Are Maintained During Acetyl-CoA Deficiency in Saccharomyces cerevisiae. Molecular & Cellular Proteomics 17, 2309–2323.

Varland, S., Myklebust, L.M., Goksoyr, S.O., Glomnes, N., Torsvik, J., Varhaug, J.E., and Arnesen, T. (2018b). Identification of an alternatively spliced nuclear isoform of human N-terminal acetyltransferase Naa30. Gene 644, 27–37.

Varland, S., Osberg, C., and Arnesen, T. (2015). N-terminal modifications of cellular proteins: The enzymes involved, their substrate specificities and biological effects. Proteomics 15, 2385–2401.

Varshavsky, A. (2019). N-degron and C-degron pathways of protein degradation. Proc Natl Acad Sci U S A 116, 358–366.

Wang, T., Birsoy, K., Hughes, N.W., Krupczak, K.M., Post, Y., Wei, J.J., Lander, E.S., and Sabatini, D.M. (2015). Identification and characterization of essential genes in the human genome. Science 350, 1096–1101.

Ward, T., Tai, W., Morton, S., Impens, F., Van Damme, P., Van Haver, D., Timmerman, E., Venturini, G., Zhang, K., Jang, M.Y., et al. (2021). Mechanisms of Congenital Heart Disease Caused by NAA15 Haploinsufficiency. Circ Res 128, 1156–1169.

Warnhoff, K., Murphy, J.T., Kumar, S., Schneider, D.L., Peterson, M., Hsu, S., Guthrie, J., Robertson, J.D., and Kornfeld, K. (2014). The DAF-16 FOXO Transcription Factor Regulates natc-1 to Modulate Stress Resistance in Caenorhabditis elegans, Linking Insulin/IGF-1 Signaling to Protein N-Terminal Acetylation. Plos Genetics 10.

Wenzlau, J.M., Garl, P.J., Simpson, P., Stenmark, K.R., West, J., Artinger, K.B., Nemenoff, R.A., and Weiser-Evans, M.C. (2006). Embryonic growth-associated protein is one subunit of a novel N-terminal acetyltransferase complex essential for embryonic vascular development. Circ Res 98, 846–855.

Wodarz, A., Hinz, U., Engelbert, M., and Knust, E. (1995). Expression of crumbs confers apical character on plasma membrane domains of ectodermal epithelia of Drosophila. Cell 82, 67–76.

Yi, C.H., Pan, H.L., Seebacher, J., Jang, I.H., Hyberts, S.G., Heffron, G.J., Vander Heiden, M.G., Yang, R.L., Li, F.P., Locasale, J.W., et al. (2011). Metabolic Regulation of Protein N-Alpha-Acetylation by Bcl-xL Promotes Cell Survival. Cell 146, 607–620.

Zaffran, S., Astier, M., Gratecos, D., and Semeriva, M. (1997). The held out wings (how) Drosophila gene encodes a putative RNA-binding protein involved in the control of muscular and cardiac activity. Development 124, 2087–2098.

Zheng, J., Edelman, S.W., Tharmarajah, G., Walker, D.W., Pletcher, S.D., and Seroude, L. (2005). Differential patterns of apoptosis in response to aging in Drosophila. Proc Natl Acad Sci U S A 102, 12083–12088.

Zhou, W.H., Xu, J., Tan, M.J., Li, H.M., Li, H., Wei, W.Y., and Sun, Y. (2018). UBE2M Is a Stress-Inducible Dual E2 for Neddylation and Ubiquitylation that Promotes Targeted Degradation of UBE2F. Molecular Cell 70, 1008-+

