## Supplementary figures and images for "N-terminal acetylation shields proteins from degradation and promotes age-dependent motility and longevity"

### Supplemental Figure 1

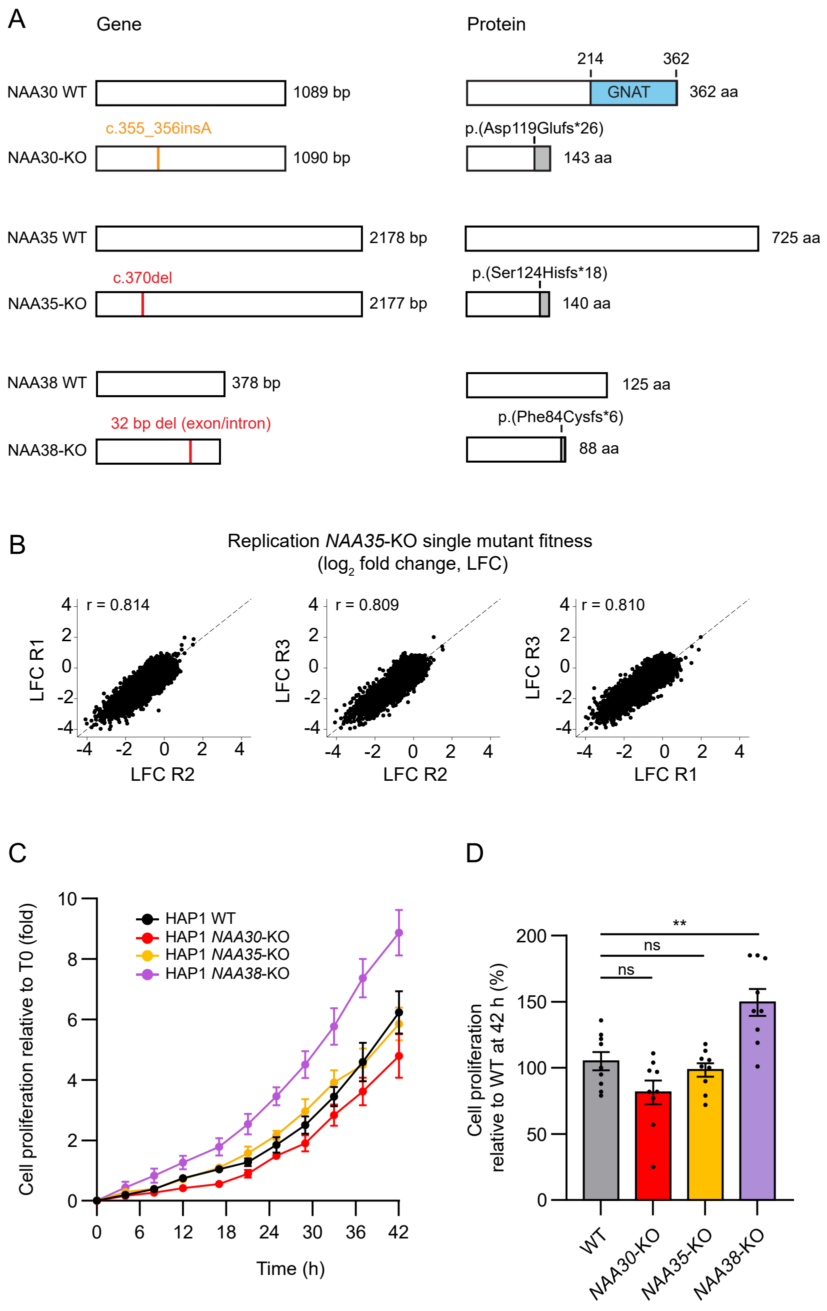

### Supplemental Figure 2

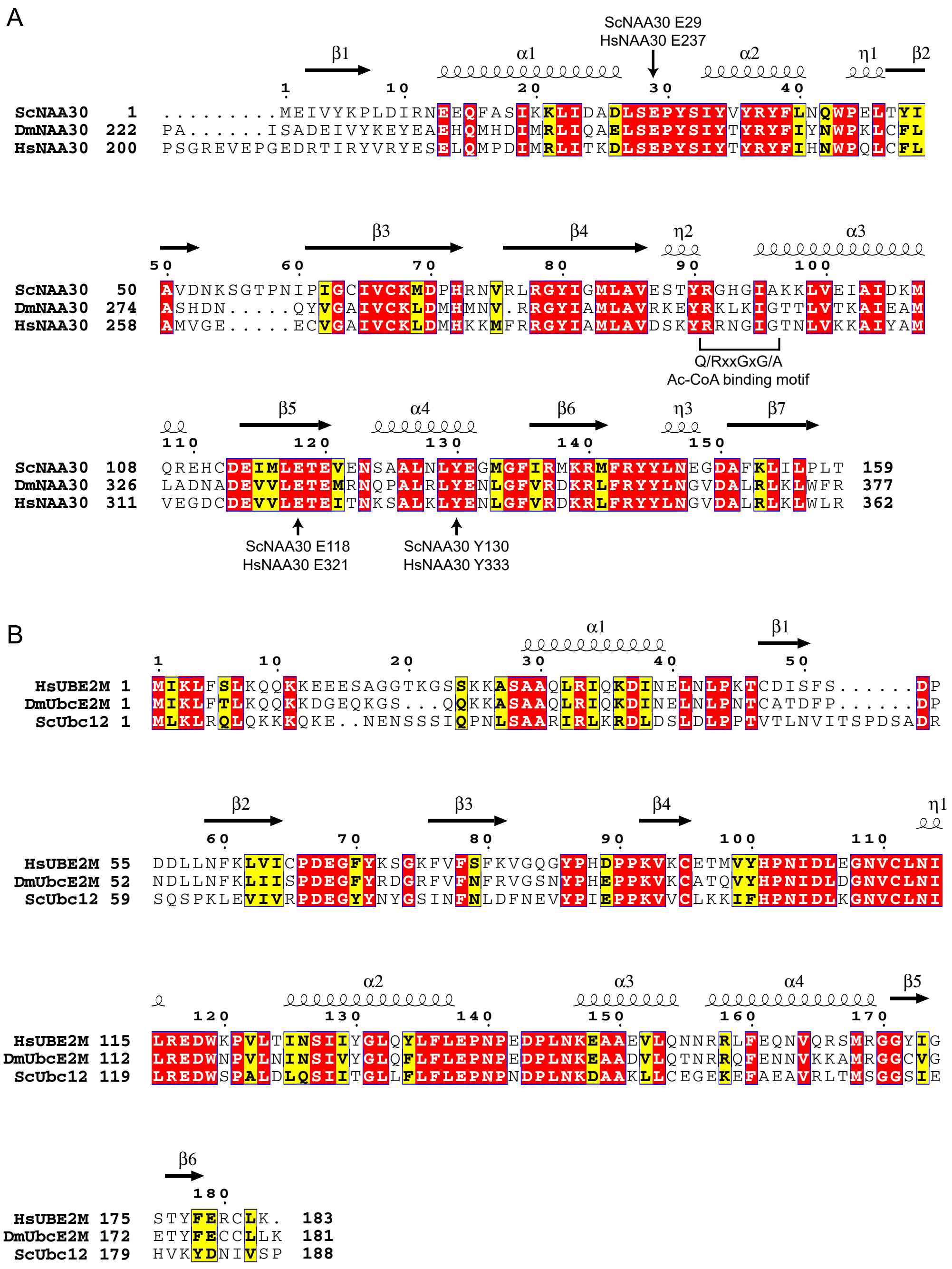

### Supplemental Figure 3

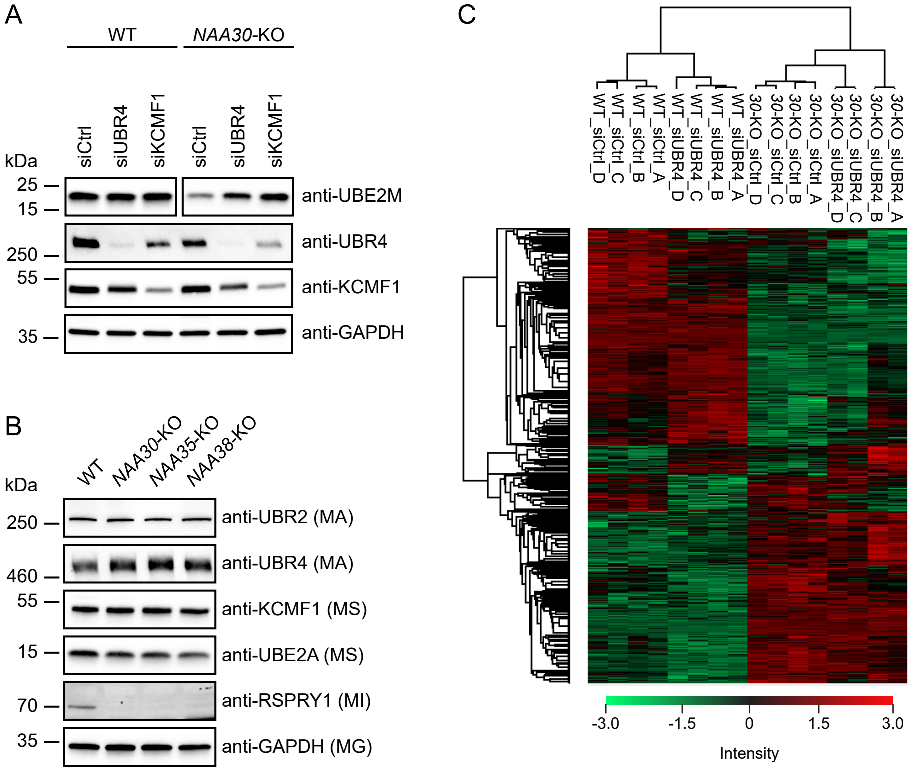

### Supplemental Figure 4

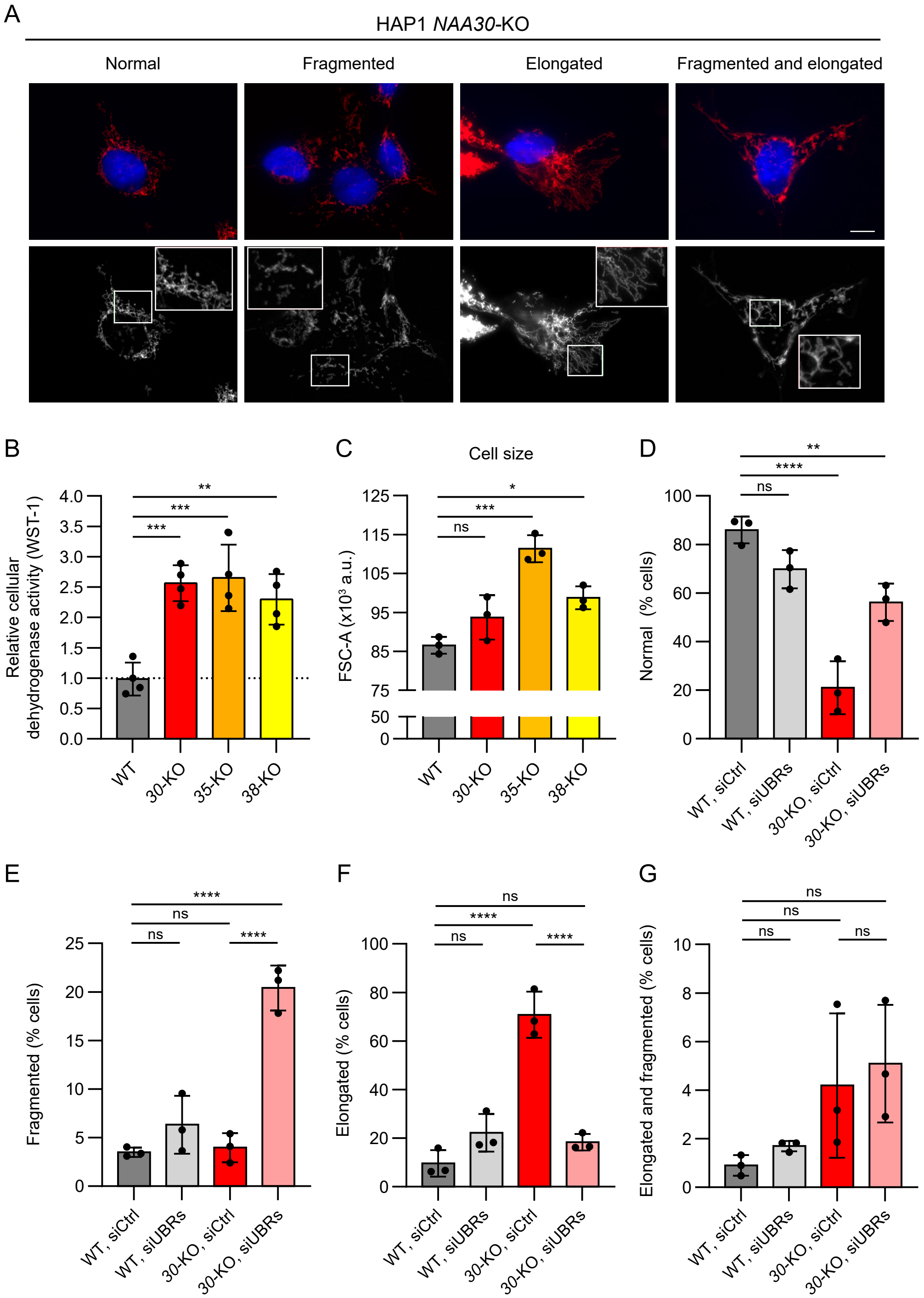

### Supplemental Figure 6

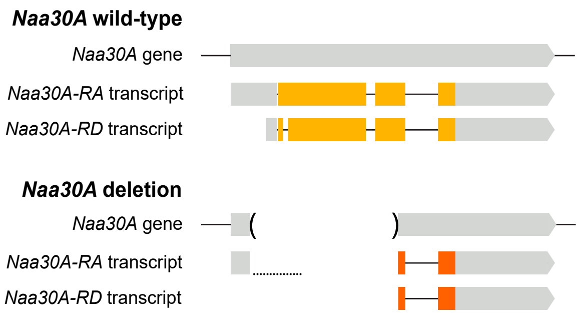

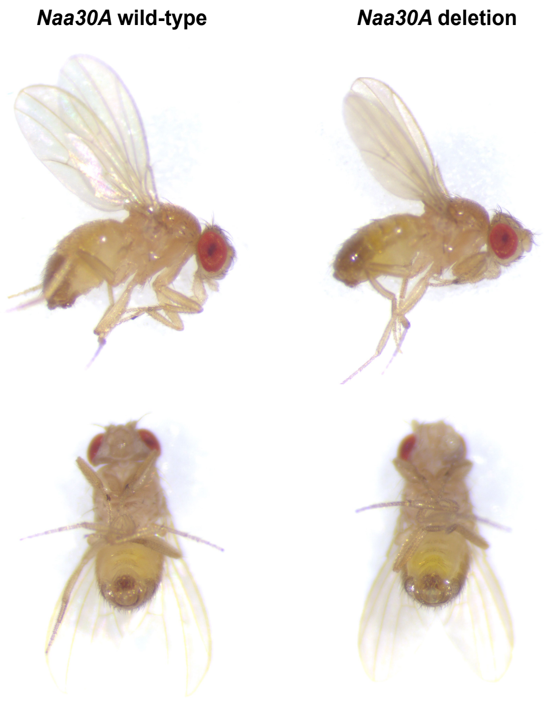

### Supplemental Figure 7

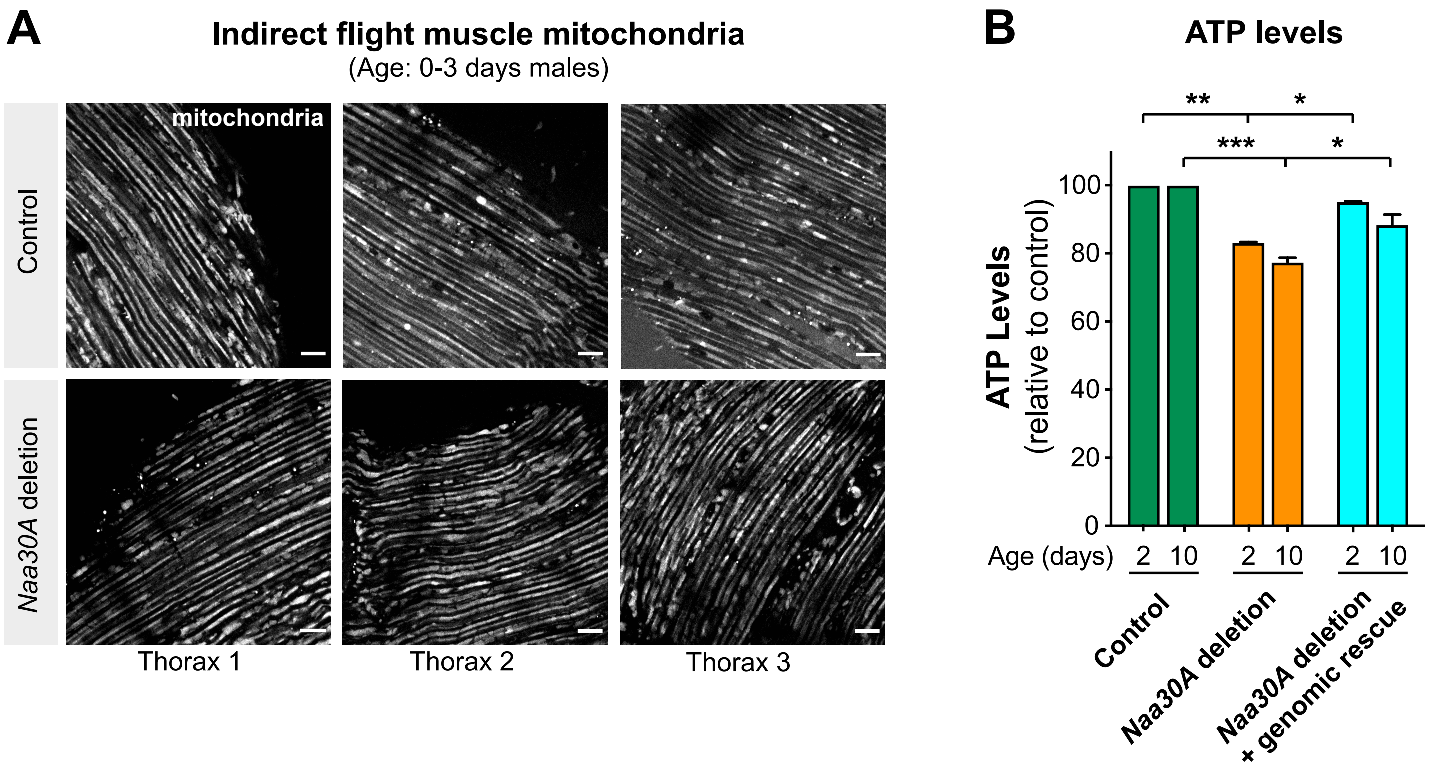
